# Rhizosphere Microbiomes in a Historical Maize/Soybean Rotation System respond to Host Species and Nitrogen Fertilization at Genus and Sub-genus Levels

**DOI:** 10.1101/2020.08.10.244384

**Authors:** Michael A. Meier, Martha G. Lopez-Guerrero, Ming Guo, Marty R. Schmer, Joshua R. Herr, James C. Schnable, James R. Alfano, Jinliang Yang

## Abstract

Root associated microbes are key players in plant health, disease resistance, and nitrogen (N) use efficiency. It remains largely unclear how the interplay of biological and environmental factors affects rhizobiome dynamics in agricultural systems. Here, we quantified the composition of rhizosphere and bulk soil microbial communities associated with maize (Zea mays L.) and soybean (Glycine max L.) in a long-term crop rotation study under conventional fertilization and low N regimes. Over two growing seasons, we evaluated the effects of environmental conditions and several treatment factors on the abundance of rhizosphere and soil colonizing microbial taxa. Time of sampling, host plant species and N fertilization had major effects on microbiomes, while no effect of crop rotation was observed. Using variance partitioning as well as 16S sequence information, we further defined a set of 82 microbial genera and sub-genus groups that show distinct responses to treatment factors. We identified taxa that are highly specific to either maize or soybean rhizospheres, as well as taxa that are sensitive to N fertilization in plant rhizospheres and bulk soil. This study provides insights to harness the full potential of soil microbes in maize and soybean agricultural systems through plant breeding and field management.

## Introduction

Crop rotations of maize and soybean exploit the symbiotic relationship of legumes with nitrogen (N) fixing bacteria. This rotation system has historically been a widespread practice in the U.S and continues to be employed as a supplement to synthetic N fertilizer (Peterson and Varvel, 1989). Soybean-maize (Jagadamma et al., 2008) and other crop rotations in general (Drinkwater et al., 1998; Peralta et al., 2018) have also shown beneficial effects on crop yield, disease resistance, weed management and soil nutrient conservation. Root-colonizing soil microbes may play a role in N use efficiency (Garnett et al., 2009), plant health (Berendsen et al., 2012) and crop performance (Yadav et al., 2018) in agricultural fields. Furthermore, the capacity of plants to recruit a specific set of beneficial microbes can potentially be employed in plant breeding and genetic engineering to improve disease resistance and yield potential of crop plants while reducing the application of exogenous fertilizer and pesticides (Chaparro et al., 2012; Compant et al., 2010; Haichar et al., 2008; Huang et al., 2014).

Soil and rhizosphere microbial communities have been studied in several major crop species including maize (Peiffer et al., 2013), soybean (Mendes et al., 2014), wheat (Donn et al., 2015) and rice (Edwards et al., 2015), as well as in crop rotation systems, including maize-wheat (Rascovan et al., 2016), wheat-maize-soybean (Gdanetz and Trail, 2017) and more complex systems (Peralta et al., 2018). Similarly, the effects of N-fertilization on microbial communities have been studied in maize (Zhu et al., 2016), wheat (Kavamura et al., 2018), and rice (Ikeda et al., 2014). These studies have shown that crop plant species, N-fertilization, and possibly crop rotation affect rhizosphere microbial community structure. However, it is largely unknown how these factors together shape rhizosphere and soil microbial communities in the context of contemporary farm management practices, and how these factors rank in terms of their impact on the abundance of distinct rhizosphere and soil colonizing microbial taxa. For instance, it has been unclear whether maize and soybean planted in succession in the same field would adopt similar root microbiomes in response to soil “memory” induced by the previous year’s crop (Lapsansky et al., 2016), or if the effect of the host plant would outweigh any crop rotation effects.

Here, we leveraged a long-term experimental field with consistent crop rotations (established 1972) and N fertilizer regimes (established 1983) (Peterson and Varvel, 1989; Varvel, 2000) in a two year replicated experiment. Through 16S sequencing of rhizosphere and bulk soil samples and statistical modeling of individual amplicon sequence variants (ASVs), we aim to rank the impact of agriculturally relevant factors, including environmental conditions (year and month of sampling), biological factors (crop plant species), and agricultural practices (N fertilization and crop rotation) on the abundance of rhizosphere and bulk soil colonizing microbes. We further aim to identify microbial taxa that respond to these diverse treatment factors as consistent units. Among these taxa, we aim to identify the key respondents that are specific to either maize or soybean, and taxa that respond to inorganic N-fertilization or the lack thereof.

## Materials & Methods

### Experimental design and sample collection

Maize and soybean plots in a historic long-term crop rotation study at the Eastern Nebraska Research Extension Center near Mead, NE (41.167380, -96.418667) were arranged in a randomized complete block design (Peterson and Varvel, 1989). Detailed site, management, yield and long-term weather information can be accessed at the USDA-ARS Agricultural Collaborative Research Outcomes System (AgCROS) website (https://agcros-usdaars.opendata.arcgis.com/). For this study, plants were sampled from two replicate blocks in each of two subsequent years (2017 and 2018). Each replication included four plots planted with continuous maize (M), continuous soybean (S), maize rotated with soybean (MS), and soybean rotated with maize (SM). Each plot contained a subplot with standard N treatment (180 kg/ha annually for maize, 68 kg/ha for soybean) and a subplot with low N conditions (no added N). From each of those subplots (experimental units), two subsamples, each for plant rhizosphere and bulk soil were collected in June, August, and September (7, 14, and 20 weeks after planting). In total, 384 samples were collected (2 years x 3 months x 2 plant species x 2 crop rotations x 2 N treatments x 2 soil compartments x 2 blocks x 2 subsamples = 384), see Fig. 1. This experimental design made it possible to distinguish 5 experimental factors: year of sampling (year 1 or year 2), month of sampling (early, mid and late season), plant species (maize or soybean), crop rotation (continuous vs. rotated), and N treatment (standard N fertilization or low N conditions). All analyses were conducted separately for rhizosphere soil and bulk soil.

**Fig. 1.**
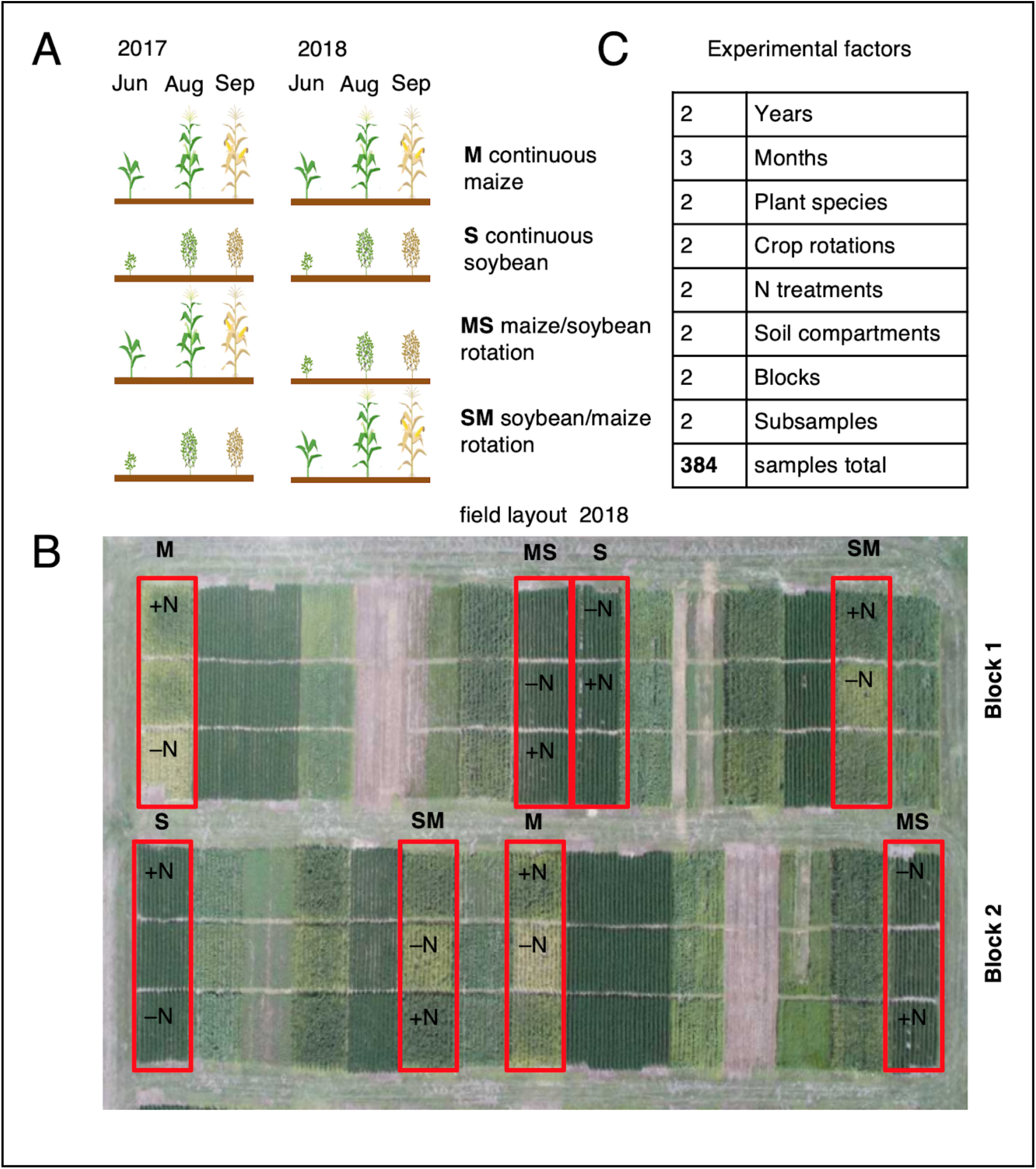
Experimental design. A) Maize (M) and soybean (S) continuous crops as well as crop rotations (MS, SM) were tracked in Jun, Aug, and Sep in two consecutive years. B) Field layout in the second year showing experimental blocks, maize or soybean plots (outlined in red) and subplots with either low (–N) or standard N treatment (+N) separated by alleys. C) Overview of treatment factors analyzed in this study.

### 16S rRNA sequencing and microbial community analysis

Genomic DNA was extracted from n=192 rhizosphere and n=192 bulk soil samples using the DNeasy PowerSoil kit (Qiagen, Hilden, Germany). Paired-end sequencing of a 300-bp sequence spanning the V4 region of the ribosomal 16 S rRNA was generated using the Illumina MiSeq platform (Illumina Inc., San Diego, CA, USA). Overall, sequencing yielded 41.4M raw 16S reads for 384 samples with a median number of 121k reads per sample for rhizosphere and 103k reads per sample for bulk soil samples. ASVs were called using a dada2-based pipeline as described by (Callahan et al., 2016a, 2016b). After a series of quality and abundance filtering steps (see Fig. S1), a final set of 4.3M reads were retained that belong to a curated set of 2,225 unique ASVs derived from both rhizosphere and bulk soil samples. The median read count per sample was 13.1k for rhizosphere and 5.9k for bulk soil samples.

### Grouping of ASVs into taxonomic groups

ASVs were initially grouped at the genus level. This is the lowest taxonomic level where groups of operational taxonomic units (OTUs) or amplicon sequence variants (ASVs) can be reliably annotated using short reads of 16S rDNA alone based on the SILVA reference database (Yilmaz et al., 2014). Sub-genus groups were further identified based on taxonomic clustering of each genus’ ASVs and associated variance partitioning data. For each of 87 genera, a phylogenetic tree of all ASVs was plotted together with the variance scores. This procedure allowed us to identify a total of 105 genera and sub-genus groups that show distinct and unambiguous responses to treatments. 82 groups that had at least five distinct ASVs were used for subsequent analyses. For each set of ASVs that mapped to a genus in which subgroups were identified, open-reference OTU picking was performed in qiime (Caporaso et al., 2010) to cluster ASVs into OTUs. The number of OTUs generated through this OTU picking procedure was compared to the number of groups identified through manual identification of genus subgroups (Table S1).

### Statistical analysis

Variance partitioning was performed on the ASV table with log transformed relative abundances to estimate the contribution of each treatment factor to changes in microbiome composition in rhizosphere and bulk soil. For each of 2,225 ASVs present in rhizospheres and a subset of 2,014 ASVs present in bulk soil, the fraction of total variance explained by each treatment factor was calculated using R package lme4 (Bates et al., 2015) with the model log(ASV relative abundance) ∼ Year + Month + Host species + Crop rotation + Nitrogen + Block + Subsample, where year, month, host species, crop rotation, nitrogen, block, and subsample were all fit as random effects. Differential abundance of taxonomic groups in response to treatments was calculated with R package DESeq2 (Love et al., 2014): starting from the ASV table with raw sequence counts, ASVs were agglomerated into 82 taxonomic groups identified above, and a +1 pseudocount was added to all table values. Unless stated otherwise, n = 96 samples were used for comparisons, e.g. 96 soybean rhizosphere samples vs. 96 maize rhizosphere samples.

For a detailed description of experimental procedures and data availability view supplementary methods.

## Results

### Rhizosphere and soil microbiomes in a historical crop rotation system are highly dynamic over time and across niche environments

Because our field experiments are subject to year and seasonal effects (Fig. S2), our first analysis was to assess how rhizosphere and bulk soil microbiomes vary across early, mid and late season sampling time points in two consecutive years. Principal coordinates analysis (Fig2 A) revealed the time of sampling to be the largest source of variation (PCoA axis 1, 34%), followed by soil compartment rhizosphere vs. bulk soil (PCoA axis 2, 20.4%). Time point variation may be attributable to temperature and precipitation patterns. In particular, the last sampling time point in 2018 occurred soon after a major precipitation event associated with drastic changes in microbial community composition (Fig. S2). Rhizosphere and bulk soil microbiomes are more dissimilar in soybean than in maize with clear separation along axis 2 in the PCoA plot. In both soil compartments, we observed higher microbial diversity in 2018 than in 2017 as measured by the Shannon diversity index (Fig2 B). In addition, both bulk soil and rhizosphere microbiomes tended to increase in diversity as the season progresses (Fig2 B).

### Environment, host plant, and agricultural practice together shape microbial communities

We fit a mixed linear model for each ASV as a response variable in order to reveal in more detail to what degree microbial communities are influenced by different treatment factors (see materials and methods). Through variance partitioning, we calculated the proportion of total variance attributable to each treatment factor (termed “variance scores”) for rhizosphere (2,225 ASVs) and bulk soil (2,014 ASVs). We tallied the number of ASVs that are responsive to treatment – defined here as any ASVs with a variance score above an arbitrary threshold of 5% – to estimate the relative importance of each treatment factor in shaping microbiome composition (Fig. 3). For rhizosphere data, out of n= 2,225 ASVs, we found 1,115 (50.1%) responsive to year and 835 (37.5%) responsive to month above the 5% threshold. For bulk soil data, out of n= 2,014 ASVs, we found 668 (34.2%) responsive to year and 639 (31.7%) responsive to month. These results agreed with our previous observations (Fig. 2) and suggested environmental factors affect microbiome abundance in the rhizosphere more than in bulk soil.

**Fig. 2.**
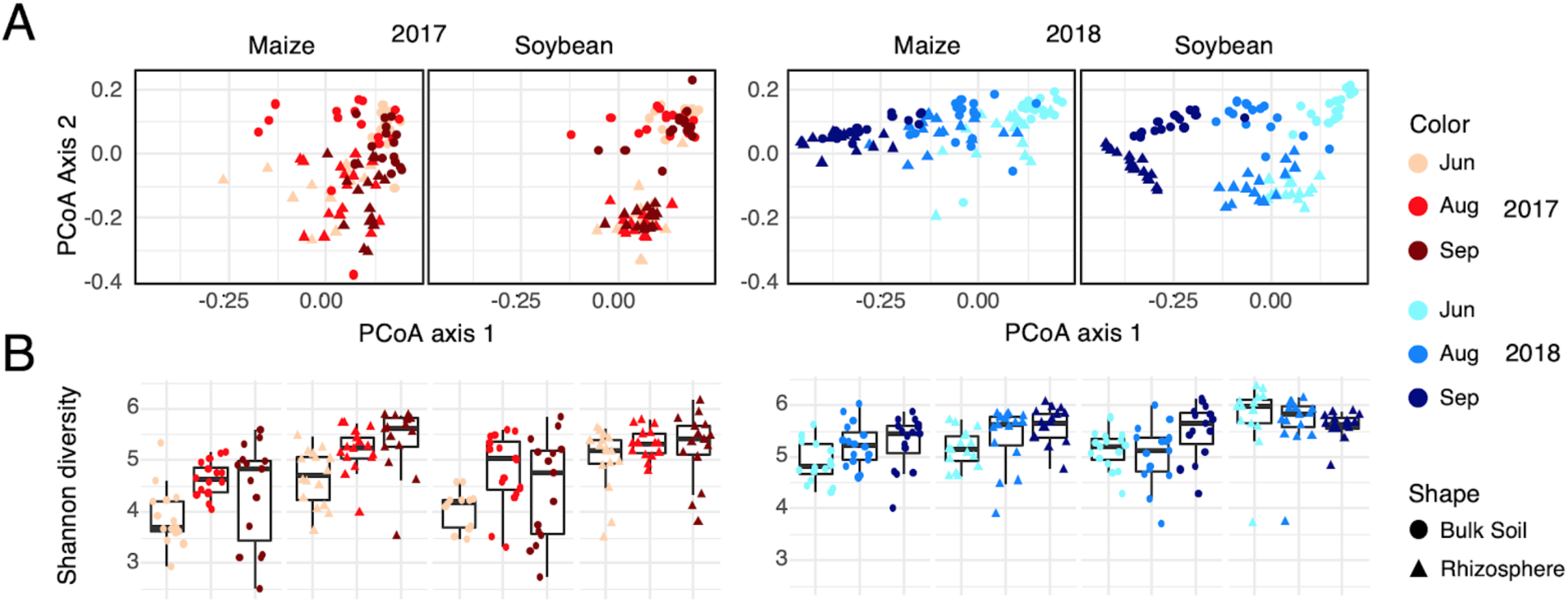
Principal Coordinates Analysis identifies time of sampling and soil compartments as major factors shaping high-level microbial community structure as measured by Shannon diversity index. A) Principal coordinates analysis (PCoA) using weighted unifrac distances, separated into four panels by year and plant species. Colors indicate sampling time point, shapes indicate soil compartments. B) Shannon diversity index plotted for each sample and summarized in box plots grouped by plant species, soil compartment and month.

**Fig 3.**
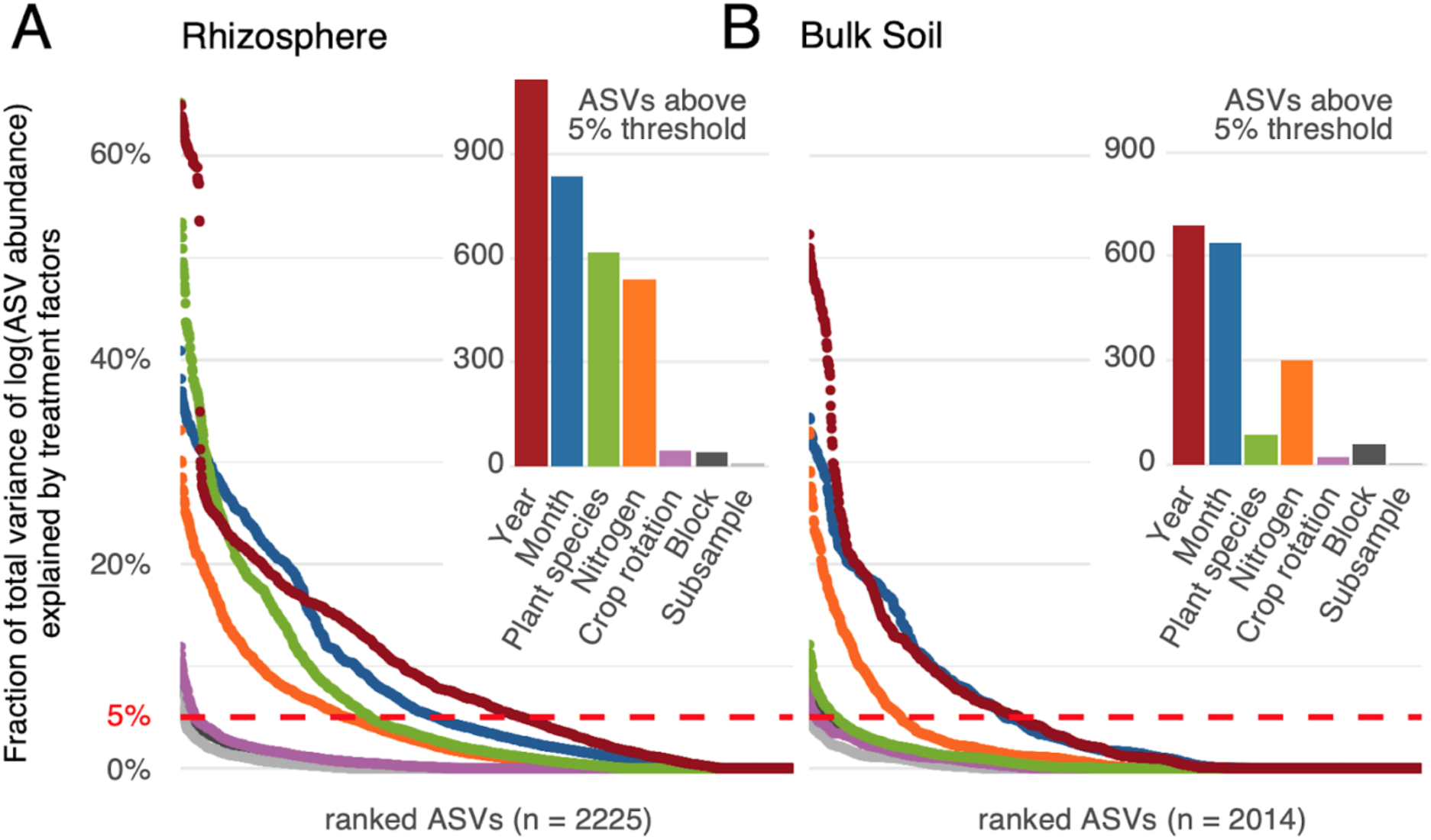
Variance partitioning results for different treatment factors influencing ASV variation in rhizosphere and bulk soil samples. For each treatment factor, percent variance explained (y-axis) was calculated for ASVs in the rhizosphere (**A**) and in bulk soil (**B**). ASVs were ranked by response to treatment factors (x-axis). Red dashed lines denote the 5% arbitrary threshold. The inset figures show the numbers of ASVs exceeding the 5% arbitrary threshold for different treatment factors.

Interestingly, microbial communities responded to host plant species to a statistically significant higher degree in the rhizosphere than the bulk soil (Chi-squared test, p-value = 2.2e-16), with variance scores of 618 ASVs in rhizosphere and only 88 ASVs in bulk soil exceeding 5%. Employing a threshold of 10% reveals a similar pattern with 422 ASVs in the rhizosphere and 9 ASVs in bulk soil exceeding the threshold (Chi-square p-value = 2.2e-16), and patterns were overall consistent at thresholds of 2.5% or 10% (Fig. S3). For 36 ASVs in the rhizosphere, more than 40% of total variance was explained by host plant species whereas no response was observed in bulk soil. These results are consistent with the idea that rhizosphere ecosystems are home to highly specialised microbes that have co-evolved alongside plant hosts, whereas bulk soil harbors more uniform microbial communities.

Among factors related to agricultural practice, we found that 5% or more variability was explained by N treatment in 539 rhizosphere ASVs and in 300 bulk soil ASVs (Chi-square p-value = 3.6e-14), with scores exceeding 20% for 71 and 42 ASVs (Chi-square p-value = 0.03267), respectively. In contrast, response to crop rotation was negligible in both rhizosphere and bulk soil, suggesting that the previous year’s crop has at best a minor effect on microbial community composition in any given year. We detected no noticeable variation due to experimental blocks and subsamples.

### Response to host species and N treatment reveals groups of microbial taxa at sub-genus level

As responses to host plant species and N treatment were apparent at the level of individual ASVs, we hypothesized that the responsive ASVs might be clustered into taxonomic groups. In order to address this hypothesis, we binned ASVs into 87 distinct microbial genera based on SILVA taxonomy annotation. However, by plotting all ASVs within each genus against the variance scores in response to plant host species and N treatment, we noticed a high range of values in some cases (Fig. S4), suggesting that there may be distinct groups of ASVs within the same genus that show different responses to treatments.

To achieve taxonomic resolution beyond the genus level, we constructed a phylogenetic tree of all ASVs in each genus together with the variance scores in response to host plant species and N treatment in the rhizosphere (see materials and methods). Using this approach, we identified subgroups in 12 genera: *Streptomyces, Chitinophaga, Flavobacterium, Pedobacter, Mucilaginibacter, Burkholderia, Pseudomonas, Sphingomonas, Sphingobium, Mesorhizobium, Nitrobacter*, and *RB41* with distinct patterns of variance partitioning (Supplementary file 1). For example, the genus *Burkholderia* (Fig. 4A & 4B), shows two clusters of ASVs (*Burkholderia_*S1, n= 29 ASVs and *Burkholderia_*S2, n= 28 ASVs) that exhibited significantly different variance scores (Wilcoxon rank sum test p-value = 2.2e-16). These clusters are further grouped by phylogeny, which may indicate separate evolutionary lineages. We refer to these groups as sub-genus groups in this study to draw a distinction between groups identified here by 16S phylogeny and variance partitioning, and microbial “species” that are categorized in some 16S sequence databases other than SILVA based on sequencing information alone. In total, a final set of 82 taxonomic groups (genera and sub-genus groups) was defined that responded to treatments as a unit. These groups spanned 64 genera and 12 classes of prokaryotes and contained between 5 and 102 ASVs, displayed in a phylogenetic tree (Fig. 4C) generated based on 300 bp 16S sequences and rooted using the outgroup *Candidtus_Nitrocosmicus* (*Archaea*). This set of 82 taxa was used for subsequent analyses in this study. Total abundances of each group were estimated by the sum of read counts across all samples (Fig. S5).

**Fig 4.**
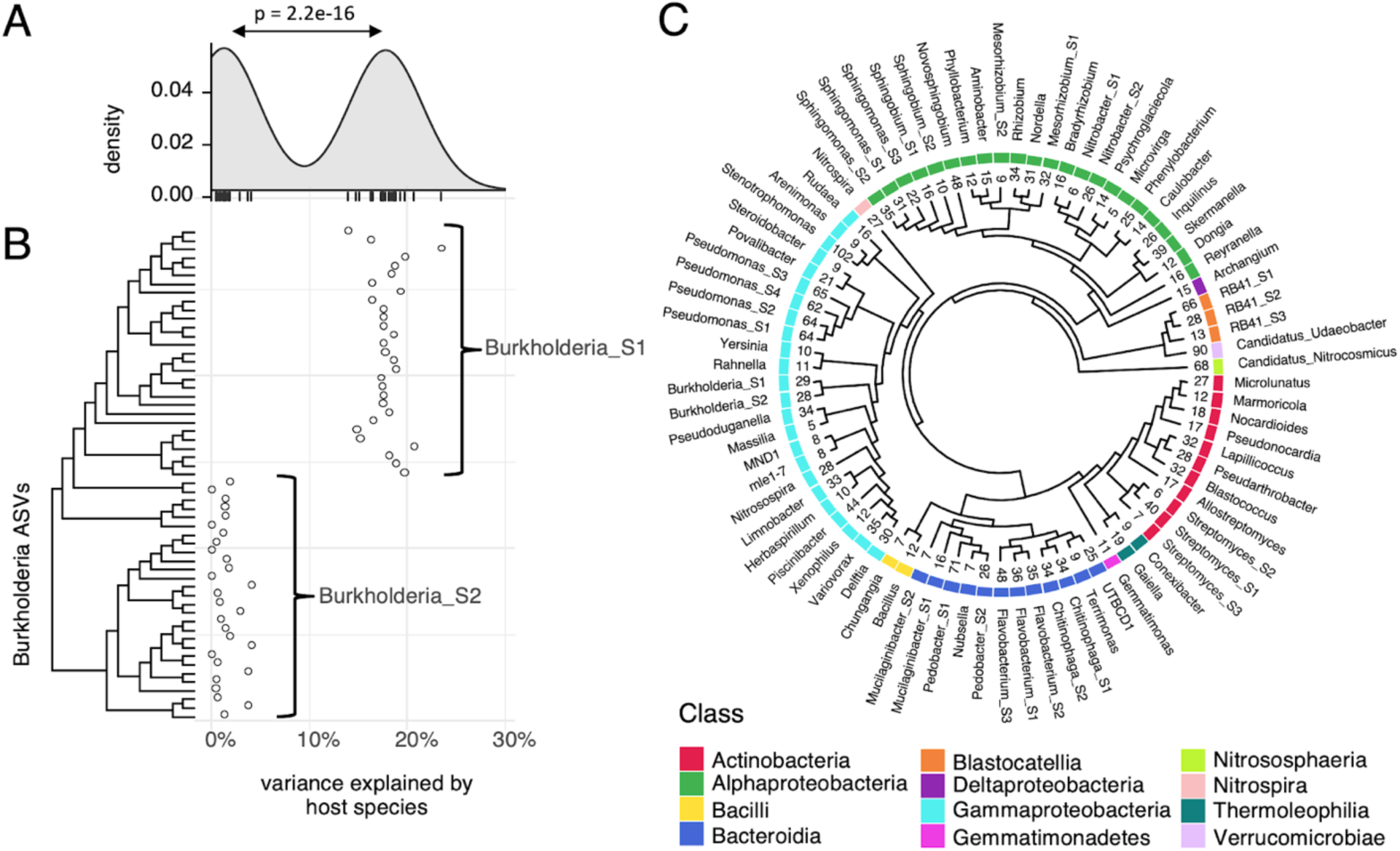
A set of 82 taxonomic groups at the genus and sub-genus level was defined based on 16S sequences and response to treatment factors. A) Variance explained by host species in rhizospheres plotted for 57 ASVs in genus *Burkholderia*. Density plot indicates bimodal distribution. B) Variance scores plotted against phylogenetic tree of ASVs in genus *Burkholderia* reveals sub-genus groups *Burkholderia_S1*, which responds to host plant species, and *Burkholderia_S2*, which is indifferent to host plant species. C) Phylogeny of 82 taxonomic groups analyzed in this study. Numbers above cladogram tips indicate the number of unique ASVs observed in each taxonomic group. Colors indicate class, tip labels indicate genus and sub-genus group (S) where applicable.

To evaluate how our ASV grouping method compares to automated OTU clustering, OTU picking was performed on the sets of ASVs within each of the 12 genera for which we identified sub-genus groups (see materials and methods). The number of sub-genus groups generated by classical OTU picking at a fixed 97% sequence identity threshold was in many cases larger than the number of subgroups identified using our method, which may indicate some redundancy (Table S1). In other cases (including *Burkholderia* in Fig. 4A & 4B), OTU picking failed to identify sub-genus groups altogether, even though variance partitioning data shows a clear distinction in the behavior of groups of ASVs.

### Host plants strongly affect rhizosphere microbial communities and have little influence over bulk soil

Fig. 5 shows 26 taxa that are responsive to host plant species in the rhizosphere, using a 5% variance score as a cutoff. The variance scores are reported together with the Log2 Fold Change (Log2FC) differential abundance of ASV counts in n = 96 soybean vs n = 96 maize rhizosphere samples, and ranked by the response to host plant species. In contrast, no taxonomic groups responded to plant species above the 5% threshold in bulk soil, with the exception of *Rudaea* (see Fig. S6 for complete data).

**Fig. 5.**
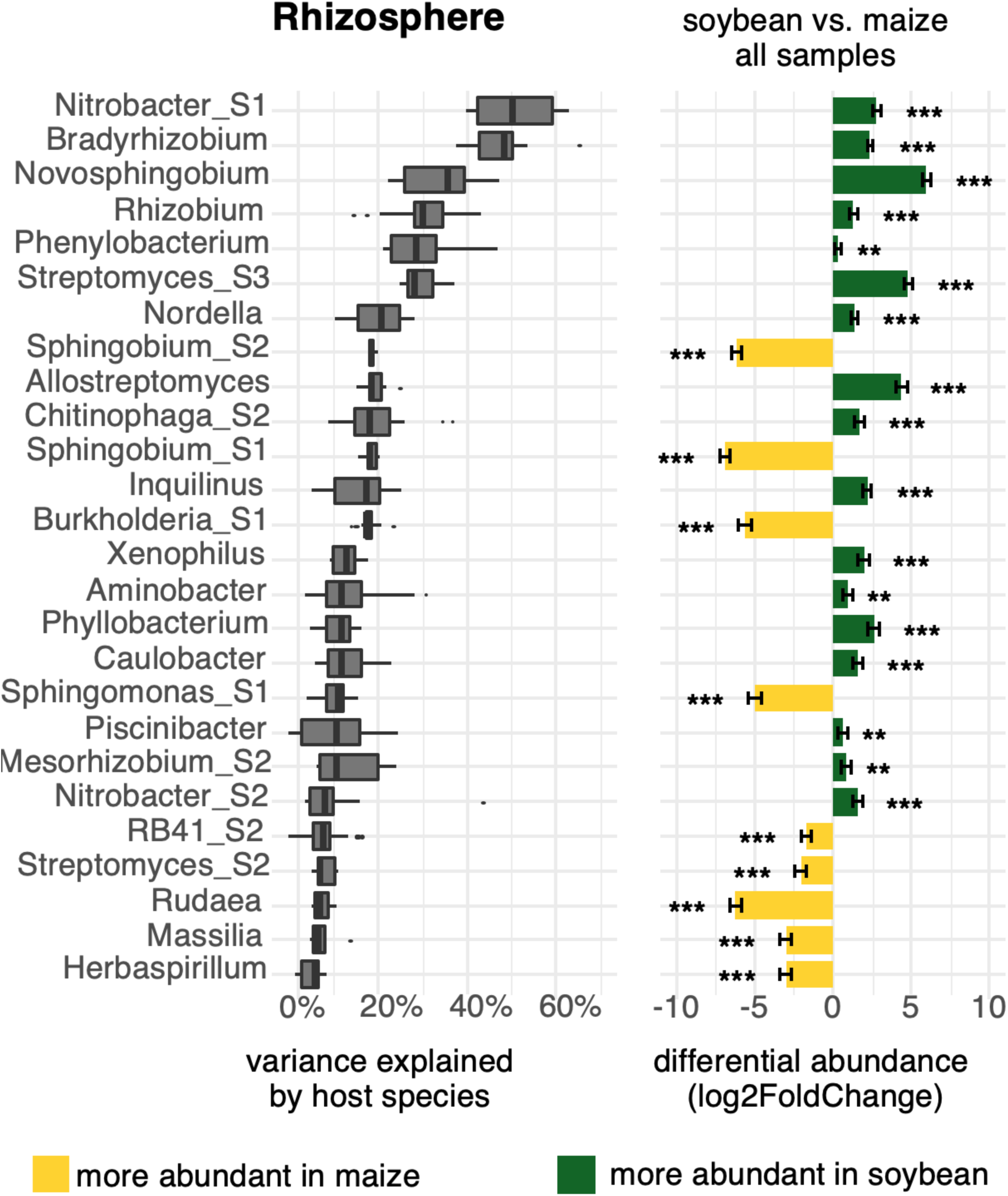
Several microbial groups are enriched in either maize or soybean rhizospheres. Taxonomic groups at the genus and sub-genus level were ranked by the fraction of total variance explained by the host plant species (left). Groups with a median variance score >5% are shown here. Differential abundance of groups, log2(abundance in soybean / abundance in maize) was calculated for n=96 pairs of rhizosphere samples using DESeq2 (right). Bars show mean +/– standard error, asterisks indicate significantly different abundance between soybean and maize at FDR-adjusted p<0.01 (***) and p<0.05 (**).

### Maize and soybean recruit distinct and highly specialized microbial taxa to rhizospheres

Several rhizosphere-dwelling taxa showed a strong response to host plant species. 9 out of the top 10 taxa responding to host plant species are specific to soybean (Fig. 5). These include *Bradyrhizobium, Rhizobium, Nordella, Nitrobacter, Novosphingobium, Phenylobacterium, Streptomyces_S1, Allostreptomyces*, and *Chitinophaga_S2*. Among the three members of the *Sphingomonadaceae* family, *Novosphingobium* (Log2FC = 5.98, FDR adjusted p-value = 1.44e-105) was highly specific to soybean, whereas *Sphingobium_S1* (Log2FC = -6.88 FDR = 1.67e-93), *Sphingobium_S2* (Log2FC = -6.12, FDR = 1.02e-83) and *Sphingomonas_S1* (Log2FC = -5.03, FDR = 6.97e-32) were specific to maize. *Sphingomonas_S2* (Log2FC = -0.77, FDR = 0.0107) shows no substantial host preference. Similarly, within the *Streptomyces* genus, *Streptomyces_S3* (Log2FC = 4.81, FDR = 7.82e-63) was highly specific to soybean whereas *Streptomyces_S2* (Log2FC = -2.00, FDR = 1.94e-07) showed a preference for maize, and *Streptomyces_S1* (Log2FC = -0.15, FDR = 0.5928) was found in roughly equal proportions in soybean and maize. *Burkholderia_S1* (Log2FC = -5.63, FDR = 2.50e-43) was highly specific to maize whereas *Burkholderia_S2* (Log2FC = 0.01 FDR = 0.9786) appears to have no preference (compare also with Fig. 4B).

### Nitrogen treatment affects soil and rhizosphere microbiomes directly and indirectly via host plant effects

Fig. 6 shows microbial taxa that respond to N treatment at a threshold of >5% variance explained. We hypothesized that the N treatment would affect rhizosphere microbiomes of maize and soybean differently, hence differential abundances of microbial taxa were analyzed separately for n=48 low N vs n=48 std N maize rhizosphere samples and for n=48 low N vs n=48 std N soybean rhizosphere samples(Fig 6A). For comparison, differential abundance of microbes between maize and soybean was shown as before (Fig 6A, rightmost panel). For bulk soil, comparisons of n=96 low N vs n=96 std N samples were made with samples from both maize and soybean fields (Fig 6B). The complete data is shown in (Fig. S7). Overall, more taxa were responsive in rhizosphere samples (n=20) than in bulk soil (n=8) at a threshold of >5% variance explained by N treatment. Notably, several taxa responded to N treatment both in bulk soil and in rhizospheres: *Nitrospira, Sphingomonas_S1* & *Sphingomonas_S2, Rudaea, Nocardioides*, and *UTBCD1* (marked bold in Fig. 6). Among these taxa, *UTBCD1* increased under low N whereas the other groups increased under std N in both bulk soil and rhizospheres. Two subgroups of genus *RB41, RB41_S1* and *RB41_S2*, were responsive to N treatment exclusively in bulk soil, whereas *RB41_S3* was responsive in both rhizospheres. *RB41_S1* and *RB41_S2* increased under std N whereas *RB41_S3* was highly increased under low N.

**Fig 6.**
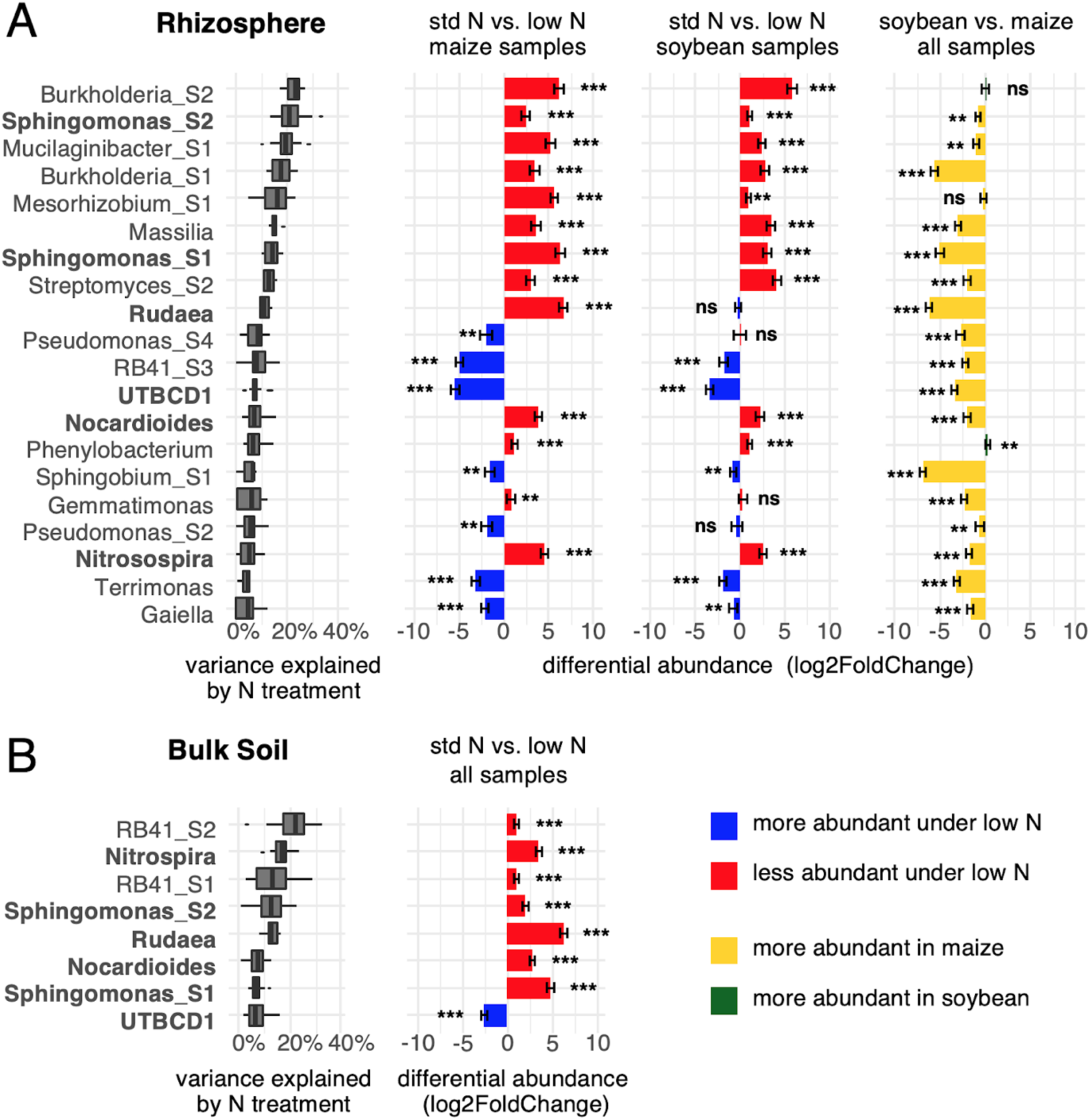
Microbial groups respond to N treatment in rhizospheres of either plant species and in bulk soil. Taxonomic groups at the genus and sub-genus level were ranked by the fraction of total variance explained by N treatment in the rhizosphere (A) and in bulk soil (B). Groups with a median variance score >5% are shown here. Differential abundance of groups, log2(abundance under std N / abundance under low N) was calculated pairwise for n=48 maize and n=48 soybean rhizosphere samples, and for n=96 bulk soil samples using DESeq2. For comparison, differential abundance in soybean vs. maize was shown in rhizosphere (green/yellow bars). Bars show mean +/– standard error, asterisks indicate significantly different abundance between std N and low N at FDR-adjusted p<0.01 (***) and p<0.05 (**). Taxa that showed response to N treatment in both rhizosphere and bulk soil are shown in bold.

Groups that mainly respond to N treatment in both rhizospheres include *Burkholderia_S1* & *Burkholderia_S2, Mucilaginibacter_S1, Mesorhizobium_S1, Massilia, Streptomyces_S2, Pseudomonas_S2* & *Pseudomonas_S4, RB41_S3, Phenylobacterium, Sphingobium_S1, Gemmatimonas, Terrimonas* and *Gaiella*.

These data suggest that N fertilization has a direct effect on the 6 microbial taxa that respond in both rhizosphere and bulk soil environments, as well as an indirect effect on taxa that only respond in rhizospheres, which is likely induced by changes in the host plant rhizosphere.

### Maize rhizosphere microbiomes are affected by N-deficiency

Differential abundance of microbial groups tends to be more extreme in maize than in soybean. This was noticed by calculating the means of absolute Log2 Fold Changes (low N ASV counts vs std N ASV counts) for maize and soybean in rhizospheres (maize mean Log2FC 1.945735 vs soybean mean Log2FC 0.9755595, Welch two sample t-test p-value 1.54e-05) as well as in bulk soil (maize mean Log2FC 1.51321 vs soybean mean Log2FC 0.8643147, p-value 0.001722). The vast majority of taxa responding to N treatment occur in greater numbers in maize than in soybean rhizospheres (Fig 6A, rightmost panel). While responses to N treatments are generally more pronounced in maize rhizospheres than in soybean rhizospheres, the direction of the changes seems to be consistent between host plant species, with a few notable exceptions: *Rudaea* are more abundant under standard N treatment than under low N in maize rhizospheres (and in bulk soil), whereas no response to N treatment was observed in soybean rhizospheres. Similarly, *Pseudomonas_S4* and *Pseudomonas_S2* increase in abundance under low N in maize rhizospheres but not in soybean rhizospheres.

Maize showed a severe N-deficiency phenotype, especially late in the season. This is known to dramatically change root architecture and exudation patterns (Gaudin et al., 2011). In contrast, soybean plants are more tolerant to a wide range of N fertilizer, which is reflected in a more stable root microbiome. Together, these data show that variation in N levels likely has a direct effect on soil microbes as well as an indirect effect through the impact of N levels on plant health and root exudation, which is most apparent in maize.

## Discussion

### Environmental factors, plant species and N treatments affect rhizosphere microbiota

Through statistical modeling of individual ASVs and variance partitioning we identified three major factors influencing rhizosphere microbial communities: time of sampling, plant species and N treatment. Year-to-year variation due to different weather conditions is common in agricultural experiments, and soil microbial communities are known to be affected by changes in temperature or humidity (Ullah et al., 2019; van der Voort et al., 2016, Fig. S2). Seasonal variation has an additional biological cause as host plant physiology – including root exudation – changes significantly as plants mature (Shi et al., 2015). Apart from environmental factors, the host plant species is the most important factor shaping rhizosphere microbiomes. Genetic distance between plant species (Fitzpatrick et al., 2018) and between genotypes of the same species (Bouffaud et al., 2014) seems to correlate with differences in microbial communities. N fertilization had an effect on both rhizosphere and soil microbial communities, which has been observed before in maize (Zhu et al., 2016). Our data show that both host plant genetics and N fertilization are major factors influencing microbial communities in maize/soybean agricultural systems. It may thus be possible to modify the composition of microbial communities in the field through plant breeding and the mode of fertilizer application, respectively.

Our data do not support a major effect of crop rotation on bulk soil or rhizosphere microbiomes when compared to other environmental and experimental factors. Thus more targeted experiments are required to discern any changes in bulk soil and rhizosphere microbiomes in response to different cropping histories at agricultural field sites.

### ASVs and variance data enable unprecedented taxonomic resolution

A common practice in observational microbiome studies is to cluster 16S amplicon sequences into operational taxonomic units (OTUs) in bins of 97% sequence similarity, and conclusions about microbial communities are drawn often at the level of bacterial phyla or classes (Bragina et al., 2015), and rarely at lower taxonomic ranks such as families (Santos-Medellín et al., 2017). However, in a highly competitive environment such as the plant rhizosphere we would expect to find highly specialized groups of microbes that react differently to a variety of treatments and any such effects would not be apparent at higher taxonomic ranks. Moreover, OTUs may not correspond to any established taxonomic rank or experimentally distinguishable group of microbes that can be studied as a unit (Yilmaz et al., 2014).

To circumvent the problems inherent to OTU clustering, we employed variance partitioning on individual amplicon sequence variants (ASVs) and used these data to complement DNA sequence information. This novel approach allowed us to identify biologically relevant taxonomic groups at the genus and sub-genus level. Importantly, we showed that traditional OTU picking would have under- or overestimated the number of sub-genus groups in most cases (Table S1). Most interestingly, the two subgroups of *Burkholderia* identified in this study, which show significantly different responses to host plant species (Fig. 4A & 4B), would have been missed entirely with traditional OTU picking. Thus, we demonstrated that multifactorial experimental designs may be exploited to improve taxonomic resolution in microbiome studies using both 16S sequence information and variance partitioning data. It may be worthwhile to formalize and automate this process using appropriate statistical tools or machine learning approaches, and to re-analyze previously published data sets whenever there are treatment factors involved that could be used to distinguish groups of microbes.

### Host plant species are a key predictor of rhizosphere microbial communities

Overall, higher microbial diversity was observed in rhizospheres than in bulk soil, which is consistent with previous studies (Prashar et al., 2014). Also in accordance with previous research (Wang et al., 2017), we observed strong responses to host plant species in both maize and soybean rhizospheres and no response in bulk soil sampled only a few centimeters away from root surfaces. An immediate effect of host plants on bulk soil microbiomes is not expected as root exudate concentrations decline exponentially and reach virtually zero only 7 mm into the soil (Kuzyakov et al., 2003).

The top 6 taxa responding to plant host species are specific to soybean. Unsurprisingly, they include N fixing bacteria such as *Bradyrhizobium, Rhizobium* and closely related *Nordella*. These were previously identified as key components of soybean microbiomes (Sugiyama et al., 2014). Nitrobacter is closely related to Bradyrhizobium and involved in Nitrite oxidation (Boon and Laudelout, 1962). *Novosphingobium, Phenylobacterium, Streptomyces* and *Allostreptomyces* have no known role in the N cycle. One notable observation was that *Novosphingobium* is highly specific to soybean, and *Sphingobium* and *Sphingomonas* are specific to maize, while all three genera are members of the *Sphingomonadaceae* family. This demonstrates once again the need for adequate taxonomic resolution when comparing microbial communities.

*Novosphingobium* has been found in the rhizosphere of *Arabidopsis* (Lin et al., 2014), maize (Kampfer et al., 2015), lettuce (Schreiter et al., 2014) and rice (Zhang et al., 2016), and to our knowledge it has not previously been reported as a prominent member of soybean rhizospheres. It remains to be confirmed whether *Novosphingobium* can be found in soybean rhizospheres in different geographic locations and in different soybean cultivars. *Sphingomonas* has been isolated previously from maize rhizospheres and proposed as a good candidate for microbial fertilizers due to N-fixation capabilities (Sun et al., 2010). A previous study (Li et al., 2014) has found *Sphingobium* to be significantly enriched in the maize rhizosphere compared to bulk soil, which was consistent with our findings. Sphingobium has also been found in rhizospheres of other grasses such as sorghum (Kochar and Singh, 2016) and common reed (Toyama et al., 2009), as well as in distantly related plants such as pine trees (Lee et al., 2019) and Kumquat (Young et al., 2008). Members of the Sphingobium genus were shown to degrade phenolic compounds such as the biocide pentachlorophenol (Dams et al., 2007) and to solubilize inorganic phosphates (Yongbin Li et al., 2017). Furthermore, An aryloxyalkanoate dioxygenase gene derived from *Sphingobium herbicidivorans* has been successfully expressed in maize to confer resistance to a broad range of herbicides (Wright et al., 2010).

Together, our results show that the host plant strongly influences microbial communities in the rhizosphere, with minimal effect on bulk soil, and that specific taxonomic groups at the genus and sub-genus level are highly adapted to either host plant. These data are consistent with the idea that maize and soybean rhizospheres are colonized by highly specialized groups of microbes that are likely in a symbiotic relationship with the host plant and may be relevant to plant health and performance.

### N treatment affects rhizosphere microbiomes both directly and indirectly via host plant effects

The vast majority of taxa responding to N fertilizer are more abundant in maize rhizospheres than in soybean rhizospheres, whereas soybean-specific taxa generally do not respond to N treatments (see Fig. 6A, rightmost panel and Fig. S6). This may be because maize shows a severely stressed phenotype under N-deficiency, especially late in the season, which induces large-scale changes to root architecture, including root hair length and density (Gaudin et al., 2011). N-limited conditions have also been shown to alter plant root exudate profiles (Baudoin et al., 2003; Haase et al., 2007). In contrast, soybean plants are hardly affected if fields are not fertilized. Thus, two factors shape microbial communities in agricultural systems: direct application of N fertilizer to the soil, which should affect both rhizosphere and bulk soil microbes, and changes due to altered root architecture and exudation patterns in response to N deficiency, which should mainly affect rhizosphere microbiomes. In accordance with this, we found more taxa affected by N treatment in rhizospheres than in bulk soil. Microbial taxa directly affected by N treatment are likely the ones that show a response to N treatment in both rhizosphere and bulk soil samples (marked in bold in Fig 6). All other taxa are likely affected indirectly, and reduced abundance under N deficiency may be due to reduced vigor of the host plant rather than due to a simple lack of inorganic N to consume.

These findings also support the idea that plant rhizospheres are colonized by highly specialized groups of microbes that are intimately tied to the host.

Taxa that increase in abundance under standard N fertilization are often capable of directly metabolizing ammonia or nitrate. *Rudaea*, a member of the *Xanthomonadaceae* family, has not been reported in maize or soybean rhizospheres but has been linked to nitrification in wastewater (Dong et al., 2016). Similarly, *Gemmatimonas, Nitrospira, Mesorhizobium, Burkholderia, Rudaea*, and *RB41* were shown to be key players in N assimilation (Morrissey et al., 2018). Burkholderia and Sphingomonas decrease in abundance under low N conditions in both maize and soybean rhizospheres, even though many members of the genus have N-fixing capabilities (Caballero-Mellado et al., 2007; Sun et al., 2010). This may indicate that the reduced abundance could also be due to changes in the rhizosphere environment other than a direct lack of N. This reinforces the idea that rhizosphere microbiomes are primarily shaped by host plant effects and to a lesser degree by external treatments such as N fertilization.

Taxa that increase in abundance under low N conditions in plant rhizospheres may be able to take advantage of reduced plant vigor under N-deficiency. Conversely, we suggest that some microbes may also be actively recruited by plants if they confer a growth or disease resistance benefit under low N stress conditions. The *Pseudomonas* genus contains both opportunistic pathogens and stranis with plant-growth promoting activity (Santoyo et al., 2012) and some groups have previously been observed in maize rhizospheres under low N conditions. *Terrimonas, Gaiella* and *Gemmatimonas* have been observed in maize rhizospheres before (Correa-Galeote et al., 2016), although their function is unknown. *UTBCD1* (*Chitinophagaceae*) and *RB41_S3* (*Pyromonadaceae*), both uncultured bacteria, increased the most under low N conditions (Liljeroth et al., 1990). Overall, surprisingly little is known about these taxa that respond positively to N-deficiency in rhizospheres and it remains to be determined whether they are simple opportunists, whether they cause disease, or whether they actively respond to changes in root exudate profiles under low N conditions, and if so, whether they have plant-growth promoting capabilities that could be exploited to improve agricultural production.

## Conclusions

In this study, we observed that rhizosphere and bulk soil microbiomes are primarily shaped by seasonal effects due to environmental changes, host plant species, and N treatment, whereas crop rotation of maize and soybean seems to be of minor importance. This suggests that maize and soybean rhizosphere microbiomes can potentially be manipulated through targeted plant breeding and farm management. We defined a set of 82 taxonomic groups at the genus and sub-genus level based on both 16S sequence information and responses to treatment variables. This allowed us to identify biologically meaningful groups of microbes that are relevant in maize and soybean production. We found groups of microbes that are highly adapted to either the maize (e.g. *Sphingobium*) or the soybean host (e.g. *Novosphingobium*), which may be relevant to plant health and performance. Lastly, we showed that N fertilization or the lack thereof has a direct effect on the abundance of several groups of microbes in bulk soil and rhizospheres as well as an indirect effect via reduced host plant vigor that is most apparent in maize. The findings presented in this work enhance our understanding of the key factors that influence rhizobiome compositions in two major crop plants under conventional and N-limited farming practices. Further research in this direction may open avenues to sustainably improve crop performance in agricultural industry.

## Supporting information

Subgroups of microbial genera

Sample Metadata

ASV table

Variance scores

Taxonomy Table

## Supplementary Data

**Fig. S1.**
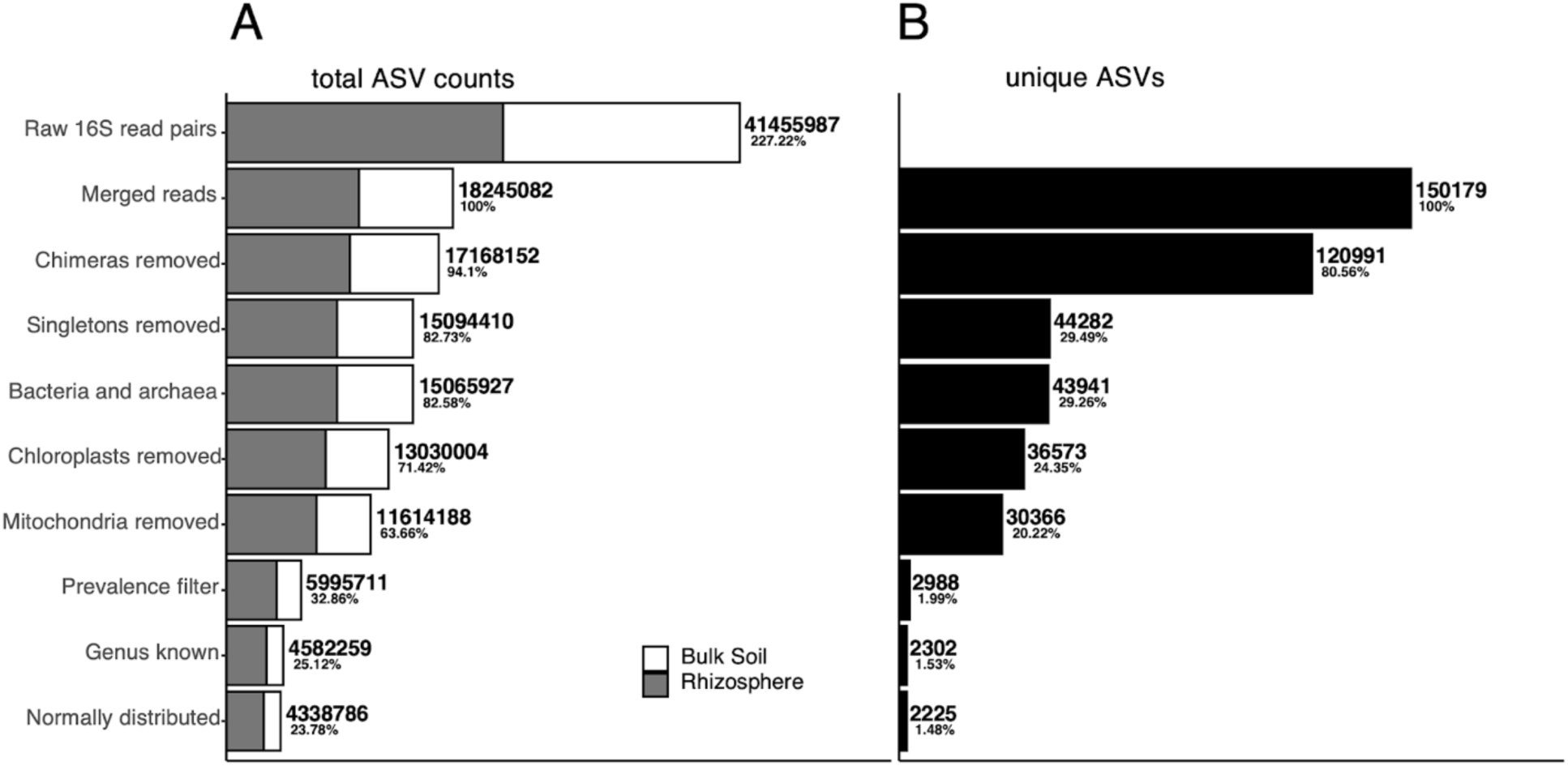
ASV counts at each filtering step in total of 384 samples. A) total ASV counts B) total unique ASVs.

**Fig S2.**
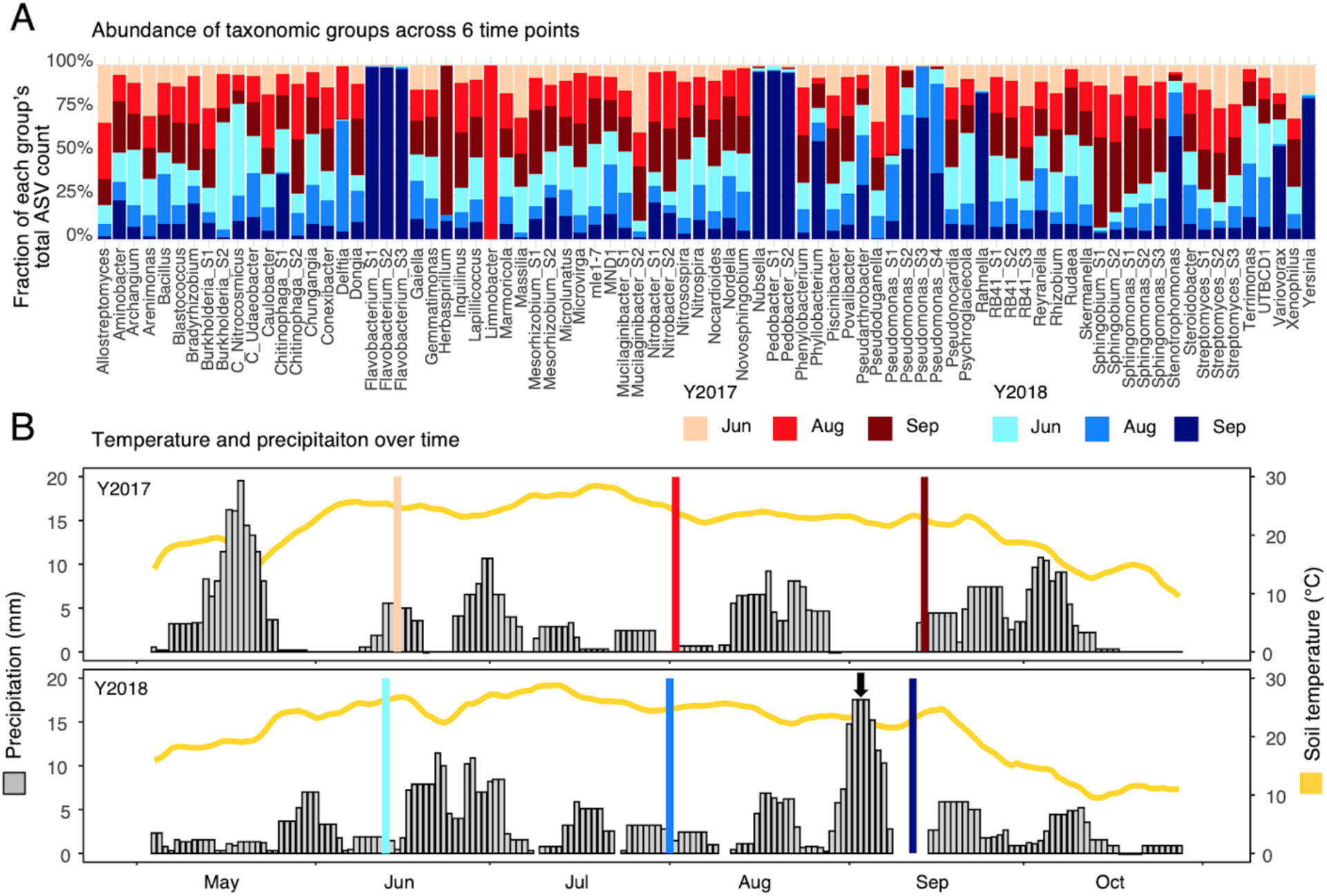
Microbial groups vary in abundance over time. Variation can be partly explained by weather effects. A) Percentage of total ASV count per taxonomic group broken down by time point. B) Soil temperature (yellow line) and precipitation data (grey bars) at USDA field site near Memphis NE. Vertical bars indicate sampling days. These data reveal unusually high counts of several taxa (*Phyllobacterium, Flavobacterium, Nubsella, Pedobacter, Pseudomonas, Rahnella, Stenotrophomonas, Variovorax, Yersinia*) in September 2018. This is possibly an environmental effect due to a heavy rainfall event only days before sample collection (arrow).

**Fig. S3.**
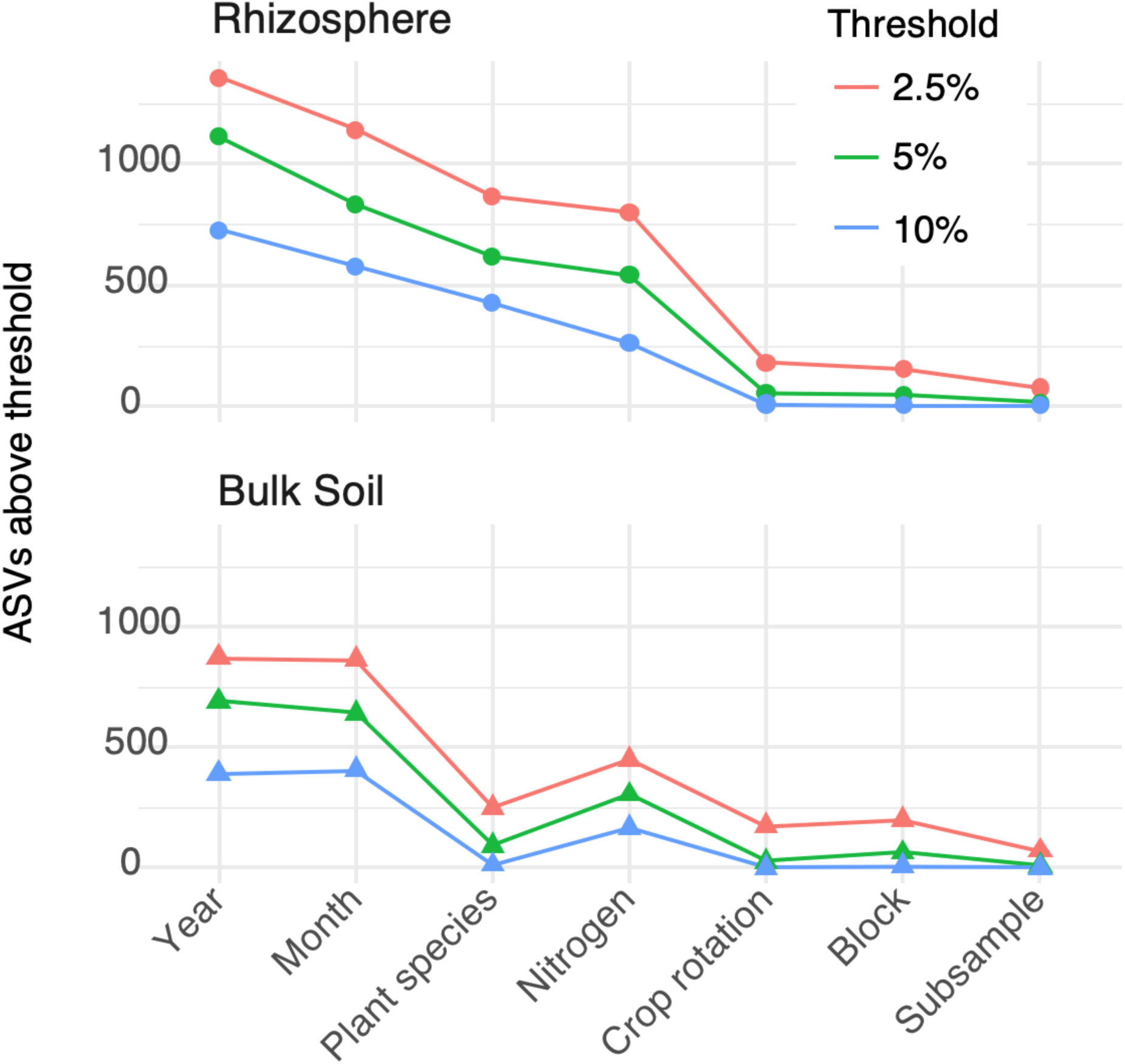
Halving or doubling the chosen 5% arbitrary threshold reveals the same ranking of treatment factors in rhizosphere and bulk soil.

**Fig. S4.**
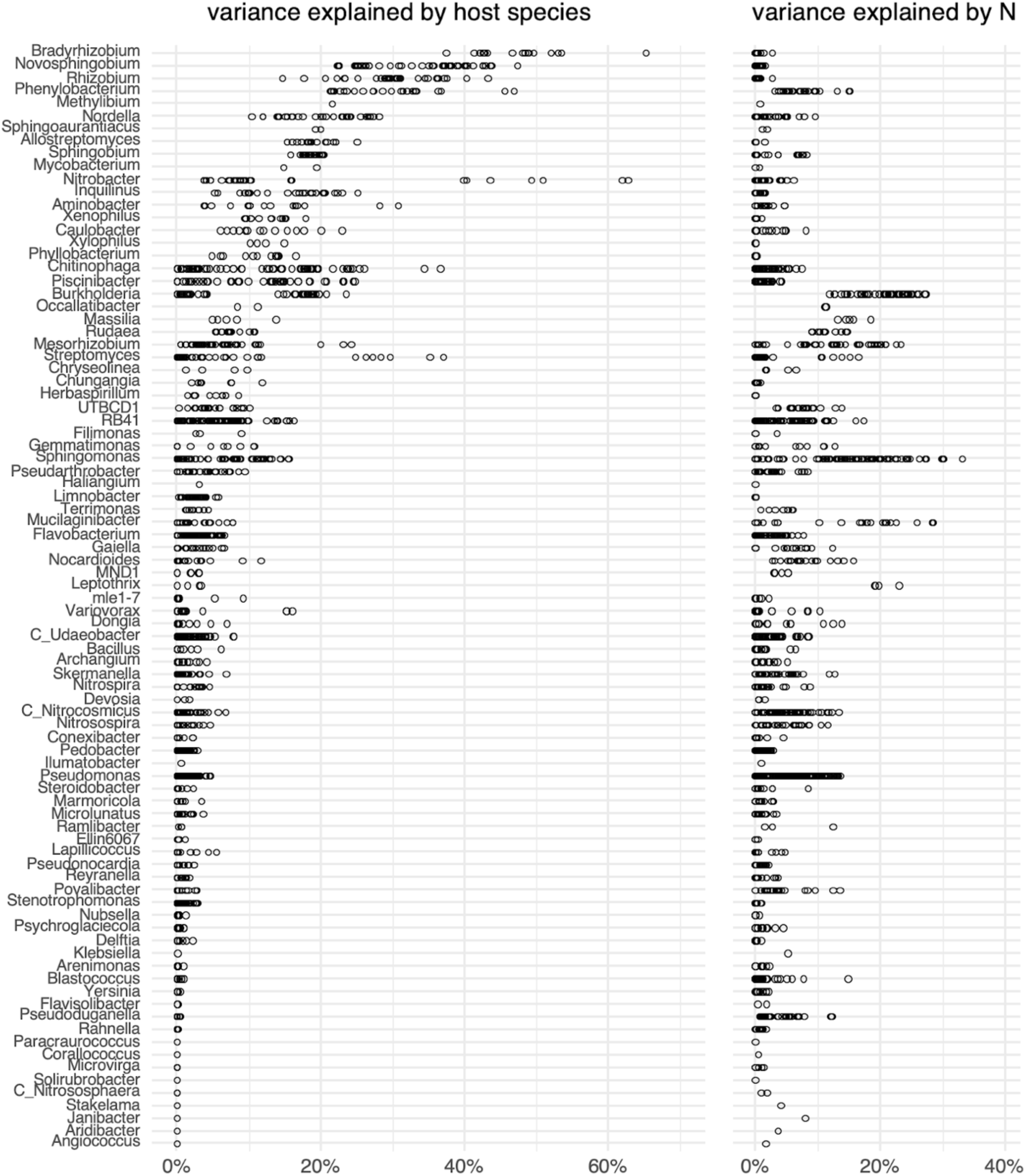
Rhizosphere Variability explained by plant species (left) and N treatment (right) for each genus. Variation in this plot may indicate that subgroups in each genus respond differently to treatments.

**Fig S5.**
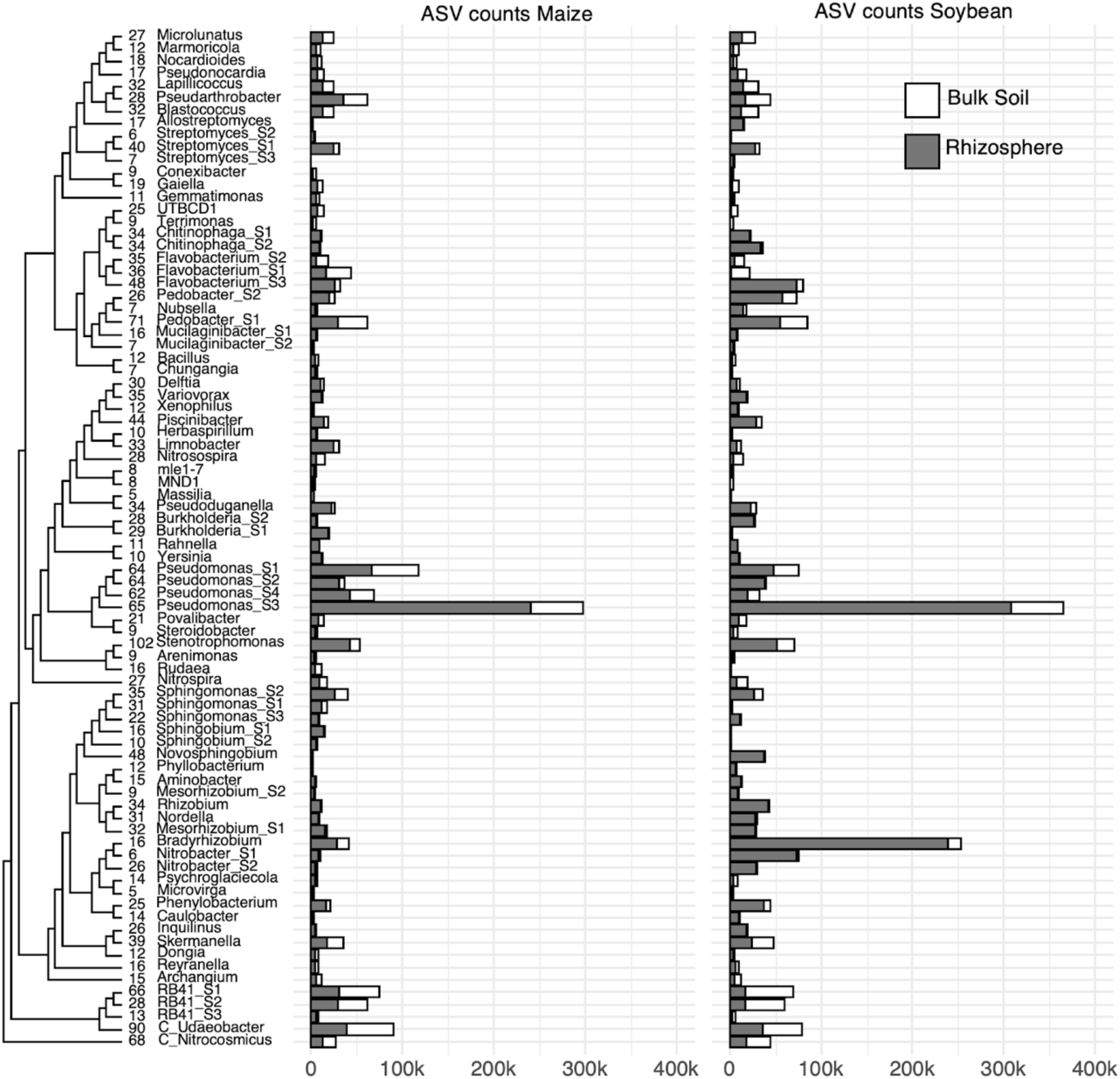
Total ASV counts in maize and soybean bulk soil and rhizosphere for all 82 taxonomic groups. Numbers above cladogram tips indicate the number of unique ASVs observed in each taxonomic group. *Pseudomonas_S3* and in soybean *Bradyrhizobium* are the most abundant, mainly in the rhizosphere.

**Fig S6.**
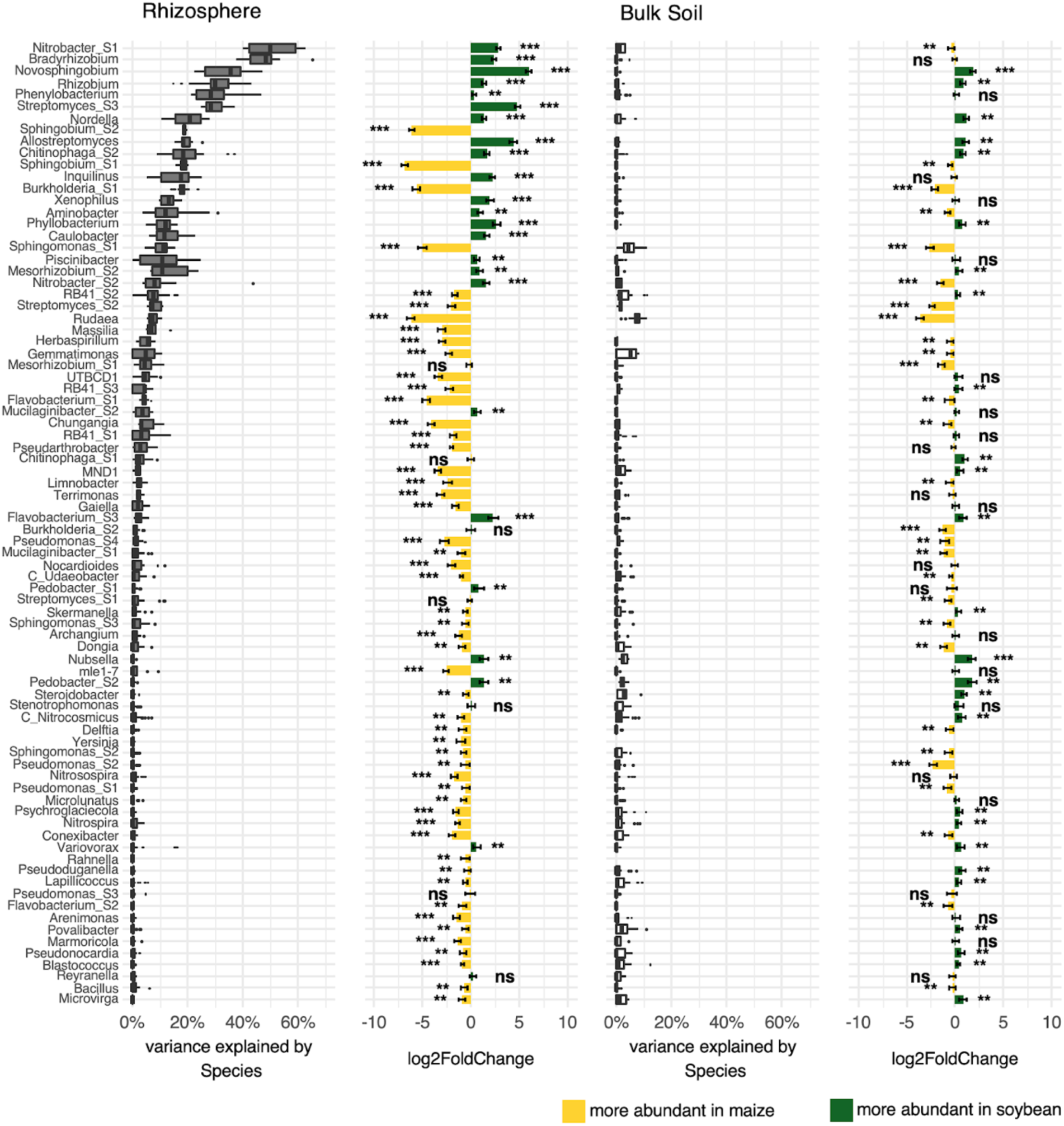
Variance explained by host plant species, all 82 taxonomic groups. Notice that the host plant effect on bulk soil is minor.

**Fig S7.**
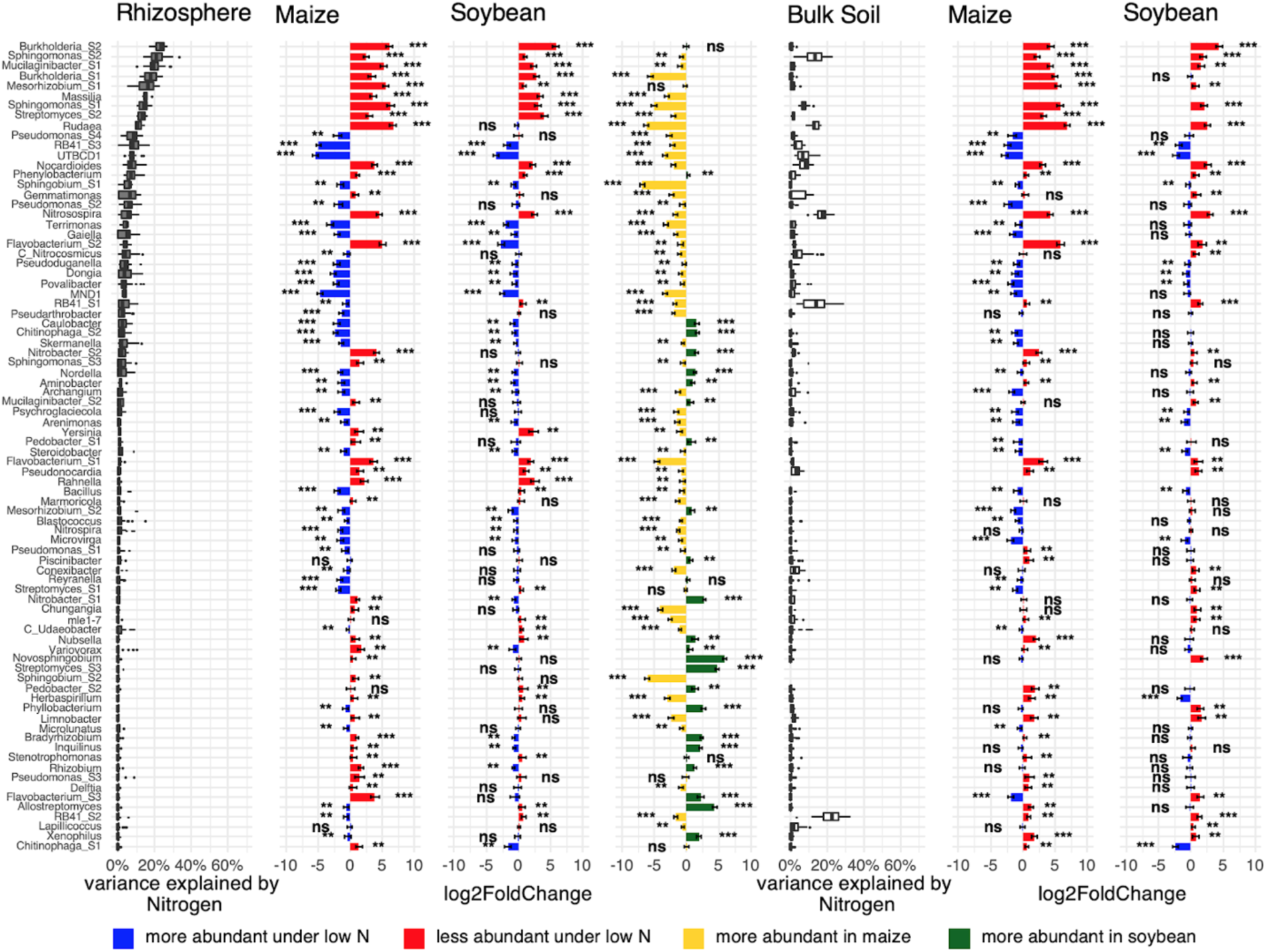
Variance explained by N treatment, all 82 taxonomic groups. Notice that ranking of groups is different in bulk soil.

**Table S1:**
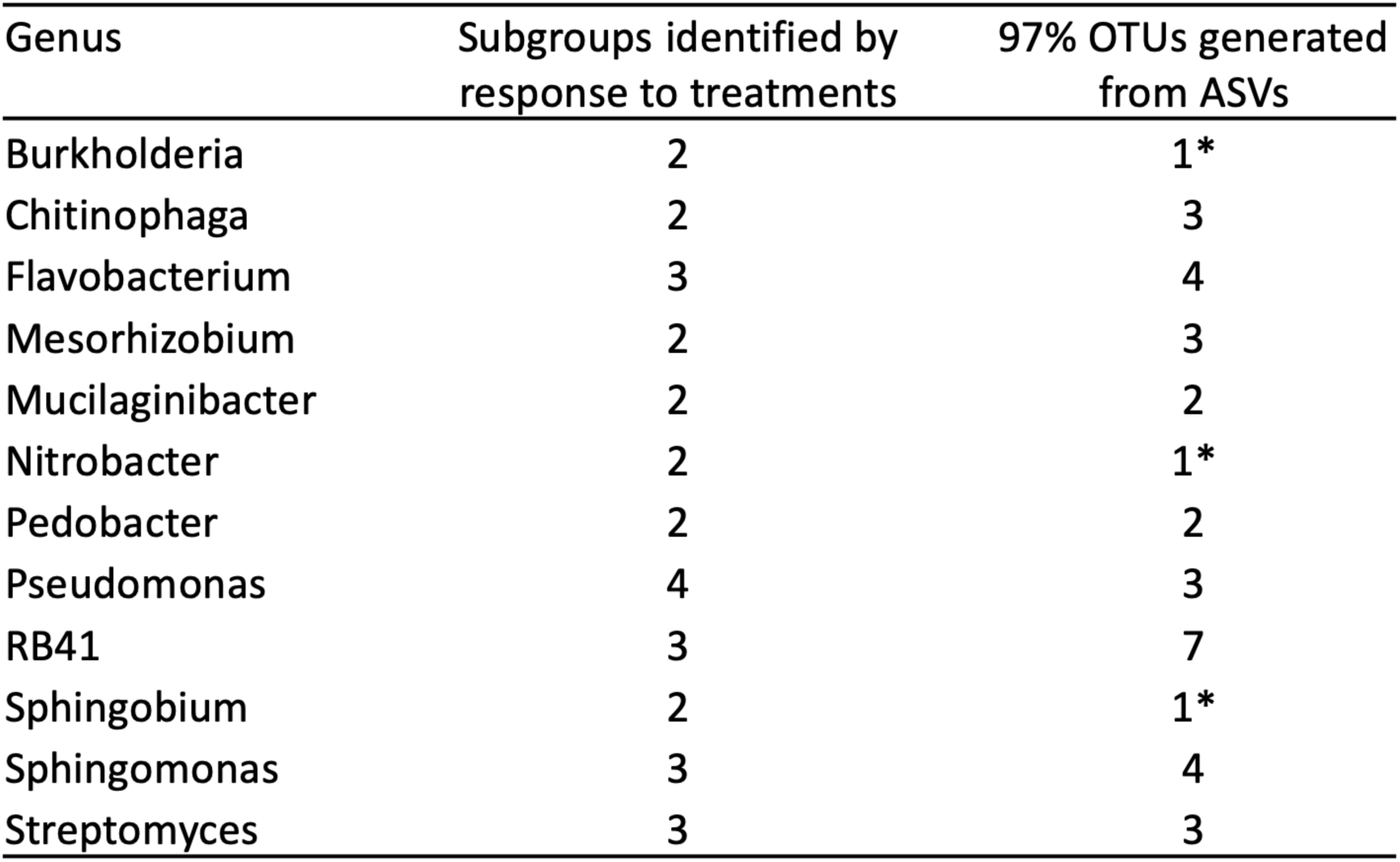
OTUs clustered from ASVs in each genus do not reproduce genus subgroups identified through variance partitioning data. OTU clustering at a fixed 97% identity threshold may fail to identify groups that respond to treatment factors as a unit. In some cases, 97% OTUs do not offer sufficient resolution to distinguish sub-genus groups at all (*).

| Genus | Subgroups identified by response to treatments | 97% OTUs generated from ASVs |
| --- | --- | --- |
| Burkholderia | 2 | 1* |
| Chitinophaga | 2 | 3 |
| Flavobacterium | 3 | 4 |
| Mesorhizobium | 2 | 3 |
| Mucilaginibacter | 2 | 2 |
| Nitrobacter | 2 | 1* |
| Pedobacter | 2 | 2 |
| Pseudomonas | 4 | 3 |
| RB41 | 3 | 7 |
| Sphingobium | 2 | 1* |
| Sphingomonas | 3 | 4 |
| Streptomyces | 3 | 3 |

## References

Bates, D., Mächler, M., Bolker, B., and Walker, S. (2015). Fitting Linear Mixed-Effects Models Using lme4. J. Stat. Softw. 67.

Berendsen, R.L., Pieterse, C.M.J., and Bakker, P.A.H.M. (2012). The rhizosphere microbiome and plant health. Trends Plant Sci. 17, 478–486.

Bouffaud, M.-L., Poirier, M.-A., Muller, D., and Moënne-Loccoz, Y. (2014). Root microbiome relates to plant host evolution in maize and other Poaceae: Poaceae evolution and root bacteria. Environ. Microbiol. 16, 2804–2814.

Bragina, A., Berg, C., and Berg, G. (2015). The core microbiome bonds the Alpine bog vegetation to a transkingdom metacommunity. Mol. Ecol. 24, 4795–4807.

Callahan, B.J., Sankaran, K., Fukuyama, J.A., McMurdie, P.J., and Holmes, S.P. (2016a). Bioconductor Workflow for Microbiome Data Analysis: from raw reads to community analyses. F1000Research 5, 1492.

Callahan, B.J., McMurdie, P.J., Rosen, M.J., Han, A.W., Johnson, A.J.A., and Holmes, S.P. (2016b). DADA2: High-resolution sample inference from Illumina amplicon data. Nat. Methods 13, 581–583.

Chaparro, J.M., Sheflin, A.M., Manter, D.K., and Vivanco, J.M. (2012). Manipulating the soil microbiome to increase soil health and plant fertility. Biol. Fertil. Soils 48, 489–499.

Compant, S., Clément, C., and Sessitsch, A. (2010). Plant growth-promoting bacteria in the rhizo- and endosphere of plants: Their role, colonization, mechanisms involved and prospects for utilization. Soil Biol. Biochem. 42, 669–678.

Dams, R.I., Paton, G.I., and Killham, K. (2007). Rhizoremediation of pentachlorophenol by Sphingobium chlorophenolicum ATCC 39723. Chemosphere 68, 864–870.

Donn, S., Kirkegaard, J.A., Perera, G., Richardson, A.E., and Watt, M. (2015). Evolution of bacterial communities in the wheat crop rhizosphere: Rhizosphere bacteria in field-grown intensive wheat crops. Environ. Microbiol. 17, 610–621.

Drinkwater, L.E., Wagoner, P., and Sarrantonio, M. (1998). Legume-based cropping systems have reduced carbon and nitrogen losses. Nature 262–265.

Edwards, J., Johnson, C., Santos-Medellín, C., Lurie, E., Podishetty, N.K., Bhatnagar, S., Eisen, J.A., and Sundaresan, V. (2015). Structure, variation, and assembly of the root-associated microbiomes of rice. Proc. Natl. Acad. Sci. 112, E911–E920.

Fitzpatrick, C.R., Copeland, J., Wang, P.W., Guttman, D.S., Kotanen, P.M., and Johnson, M.T.J. (2018). Assembly and ecological function of the root microbiome across angiosperm plant species. Proc. Natl. Acad. Sci. 115, E1157–E1165.

Garnett, T., Conn, V., and Kaiser, B.N. (2009). Root based approaches to improving nitrogen use efficiency in plants. Plant Cell Environ. 32, 1272–1283.

Gaudin, A.C.M., Mcclymont, S.A., Holmes, B.M., Lyons, E., and Raizada, M.N. (2011). Novel temporal, fine-scale and growth variation phenotypes in roots of adult-stage maize (Zea mays L.) in response to low nitrogen stress: Nitrogen stress on maize roots. Plant Cell Environ. 34, 2122–2137.

Gdanetz, K., and Trail, F. (2017). The Wheat Microbiome Under Four Management Strategies, and Potential for Endophytes in Disease Protection. Phytobiomes J. 1, 158–168.

Haichar, F. el Z., Marol, C., Berge, O., Rangel-Castro, J.I., Prosser, J.I., Balesdent, J., Heulin, T., and Achouak, W. (2008). Plant host habitat and root exudates shape soil bacterial community structure. ISME J. 2, 1221–1230.

Huang, X.-F., Chaparro, J.M., Reardon, K.F., Zhang, R., Shen, Q., and Vivanco, J.M. (2014). Rhizosphere interactions: root exudates, microbes, and microbial communities. Botany 92, 267–275.

Ikeda, S., Sasaki, K., Okubo, T., Yamashita, A., Terasawa, K., Bao, Z., Liu, D., Watanabe, T., Murase, J., Asakawa, S., et al. (2014). Low Nitrogen Fertilization Adapts Rice Root Microbiome to Low Nutrient Environment by Changing Biogeochemical Functions. Microbes Environ. 29, 50–59.

Jagadamma, S., Lal, R., Hoeft, R.G., Nafziger, E.D., and Adee, E.A. (2008). Nitrogen fertilization and cropping system impacts on soil properties and their relationship to crop yield in the central Corn Belt, USA. Soil Tillage Res. 98, 120–129.

Kavamura, V.N., Hayat, R., Clark, I.M., Rossmann, M., Mendes, R., Hirsch, P.R., and Mauchline, T.H. (2018). Inorganic Nitrogen Application Affects Both Taxonomical and Predicted Functional Structure of Wheat Rhizosphere Bacterial Communities. Front. Microbiol. 9, 1074.

Kuzyakov, Y., Raskatov, A., and Kaupenjohann, M. (2003). Turnover and distribution of root exudates of Zea mays. Plant Soil 254, 317–327.

Li, X., Rui, J., Mao, Y., Yannarell, A., and Mackie, R. (2014). Dynamics of the bacterial community structure in the rhizosphere of a maize cultivar. Soil Biol. Biochem. 68, 392–401.

Love, M.I., Huber, W., and Anders, S. (2014). Moderated estimation of fold change and dispersion for RNA-seq data with DESeq2. Genome Biol. 15, 550.

Mendes, L.W., Kuramae, E.E., Navarrete, A.A., van Veen, J.A., and Tsai, S.M. (2014). Taxonomical and functional microbial community selection in soybean rhizosphere. ISME J. 8, 1577–1587.

Peiffer, J.A., Spor, A., Koren, O., Jin, Z., Tringe, S.G., Dangl, J.L., Buckler, E.S., and Ley, R.E. (2013). Diversity and heritability of the maize rhizosphere microbiome under field conditions. Proc. Natl. Acad. Sci. 110, 6548–6553.

Peralta, A.L., Sun, Y., McDaniel, M.D., and Lennon, J.T. (2018). Crop rotational diversity increases disease suppressive capacity of soil microbiomes. Ecosphere 9, e02235.

Peterson, T.A., and Varvel, G.E. (1989). Crop Yield as Affected by Rotation and Nitrogen Rate.Soybean. Agron. J. 81, 727.

Prashar, P., Kapoor, N., and Sachdeva, S. (2014). Rhizosphere: its structure, bacterial diversity and significance. Rev. Environ. Sci. Biotechnol. 13, 63–77.

Rascovan, N., Carbonetto, B., Perrig, D., Díaz, M., Canciani, W., Abalo, M., Alloati, J., González-Anta, G., and Vazquez, M.P. (2016). Integrated analysis of root microbiomes of soybean and wheat from agricultural fields. Sci. Rep. 6, 28084.

Santos-Medellín, C., Edwards, J., Liechty, Z., Nguyen, B., and Sundaresan, V. (2017). Drought Stress Results in a Compartment-Specific Restructuring of the Rice Root-Associated Microbiomes. MBio 8, e00764–17.

Sun, J.-G., Zhang, Y.-C., Xu, J., and Hu, H.-Y. (2010). Isolation, identification and inoculation effect of nitrogen-fixing bacteria <I>Sphingomonas</I> GD542 from maize rhizosphere: Isolation, identification and inoculation effect of nitrogen-fixing bacteria <I>Sphingomonas</I> GD542 from maize rhizosphere. Chin. J. Eco-Agric. 18, 89–93.

Varvel, G.E. (2000). Crop Rotation and Nitrogen Effects on Normalized Grain Yields in a Long-Term Study. Agron. J. 92, 938.

Wang, P., Marsh, E.L., Ainsworth, E.A., Leakey, A.D.B., Sheflin, A.M., and Schachtman, D.P. (2017). Shifts in microbial communities in soil, rhizosphere and roots of two major crop systems under elevated CO2 and O3. Sci. Rep. 7, 15019.

Yadav, A.N., Kumar, V., Dhaliwal, H.S., Prasad, R., and Saxena, A.K. (2018). Microbiome in Crops: Diversity, Distribution, and Potential Role in Crop Improvement. In Crop Improvement Through Microbial Biotechnology, (Elsevier), pp. 305–332.

Yilmaz, P., Parfrey, L.W., Yarza, P., Gerken, J., Pruesse, E., Quast, C., Schweer, T., Peplies, J., Ludwig, W., and Glöckner, F.O. (2014). The SILVA and “All-species Living Tree Project (LTP)” taxonomic frameworks. Nucleic Acids Res. 42, D643–D648.

Zhu, S., Vivanco, J.M., and Manter, D.K. (2016). Nitrogen fertilizer rate affects root exudation, the rhizosphere microbiome and nitrogen-use-efficiency of maize. Appl. Soil Ecol. 107, 324–333.

