## Supplementary Methods for "Rhizosphere Microbiomes in a Historical Maize/Soybean Rotation System respond to Host Species and Nitrogen Fertilization at Genus and Sub-genus Levels"

Experimental field

Root and soil samples for this study were collected in 2017 and in 2018 from rain-fed experimental plots managed by USDA-ARS that had consistent long-term crop rotations and nitrogen fertilizer regimes in place for close to four decades. Plots are located at the University of Nebraska Agricultural research and Development Center near Mead, NE at the geographical location [41°10'00.7"N 96°25'07.2"W]. In brief, continuous maize (Zea mays L., Pioneer P1751AMT), continuous soybean (Glycine max L., Pioneer P31T11R), maize/soybean and other crop rotations were arranged in a randomized complete block design. Subplots were randomly assigned either low nitrogen (no extra fertilizer) medium nitrogen or high nitrogen treatments (180 kg/ha ammonium nitrate annually for maize, 68 kg/ha for soybean). Details about the rotation study are described by Peterson and Varvel. (Peterson and Varvel, 1989; Varvel, 2000).

Rhizobiome sampling

Root and soil samples were collected in June, early August, and September (7, 14, and 20 weeks after planting) in both 2017 and 2018. Samples were taken from maize and soybean plants in two different cropping systems (continuous and rotated) with two replicate plots per plant species and cropping system. Continuous crops were continuous maize (plots 115, 208), continuous soybean (plots 107, 215), crop rotations were maize in 2017/soybean in 2018 (plots 108 & 201 ) and soybean in 2017/maize in 2018 (plots 102, 210). On each of the 8 plots, samples were taken from both the low nitrogen and high nitrogen subplot. On each subplot, two subsamples were taken for a total of 4 replicates per plant species / cropping system / nitrogen treatment combination. Each replicate sample in turn consisted of pooled material from two adjacent plants that were randomly selected from the inner rows of each subplot.

Plant roots were dug up to a depth of 30 cm and rootstocks were manually shaken to remove loosely adherent soil. This soil was collected from two adjacent plants, homogenized and 15 ml was collected in a 50 ml tube. We called this fraction bulk soil. To gather a representative root sample, roots from two plants were cut into 5 cm pieces, homogenized and one 50 ml tube was filled with random root material. In soybean, root nodules were removed before cutting the roots to decrease the abundance of (well studied) nodule-forming rhizobium symbionts. Also, as soybean roots are relatively small early in the growing season up to 5 adjacent plants were pooled to gather enough root material for the June time points.

Sample processing

Root and bulk soil samples were immediately put on ice, transferred to the lab and processed within 24 h. 30 ml phosphate buffer (46 mM NaH2PO4, 60 mM NA2HPO4, 200 μl/L Silwet-77) was added to the root samples in 50 ml tubes and samples were vortexed horizontally for 3 min at 8000 rpm to shake tightly adherent soil off the roots. This fraction we call rhizosphere. The rhizosphere soil suspension was filtered through a 100 μm nylon cell strainer (Celltreat Scientific Products, Pepperell, MA, USA) to remove residual plant material. The filtrate was centrifuged for 10 min at 4000 g. Supernatant was discarded and the soil pellet was stored at -20°C. For bulk soil samples, 30 ml phosphate buffer was added to 15 ml soil collected from the field and the sample was vortexed, filtered and processed in the same way as the rhizosphere samples.

In total, 2 crop species x 2 crop rotations x 2 N treatments x 2 years x 3 time points x 4 replicates x 2 fractions amounted to 384 samples.

DNA isolation and quantification

DNA was isolated using the DNeasy PowerSoil kit (Qiagen, Hilden, Germany) for both rhizosphere and bulk soil samples, starting with up to 250 mg material. Differing from the standard protocol, all samples were eluted in 50 μl TE (10 mM Tris-HCl, 1 mM disodium EDTA, pH 8.0) passed twice over the ion exchange column to increase DNA concentration. DNA was quantified fluorometrically using the QuantiFluor® dsDNA System (Promega, Madison, WI, USA). DNA extraction was repeated with fresh material if DNA concentration was less than 1 ng/µl.

16 S library preparation and sequencing

DNA samples were processed at the University of Minnesota Genomics Center (Minneapolis, MN, USA). In brief, the 16S rRNA V4 region was amplified using V4_515F_Nextera and V4_806R_Nextera primers and sequencing library preparation as described by Gohl (Gohl et al., 2016). For the 2018 samples, oligonucleotide PCR blockers (PNA Bio INC, Thousand Oaks, CA, USA) targeting mitochondrial and chloroplast sequences were applied in the primary V4 amplification to reduce amplification of templates derived from eukaryotes. Up to 120 barcoded samples were pooled per sequencing run and sequenced on an Illumina MiSeq platform.

Raw read processing and construction of ASV table

16S sequencing reads were processed in R 3.5.2 using a workflow described by Callahan (Callahan et al., 2016b), which employs the package dada2 1.10.1(Callahan et al., 2016a). Cluster computing resources at the UNL Holland Computing Center were used for computationally demanding steps. In brief, ~300 bp raw sequencing reads were trimmed using *filterAndTrim()* at 240 bp (forward reads) and 200 bp (reverse reads), respectively. Amplicon sequence variants (ASV) were inferred using *dada()* and forward and reverse reads were merged with *mergePairs()*. A sequence table was generated using *makeSequenceTable()* and chimaeras were removed using *removeBimeraDenovo()*. Taxonomy was assigned to ASVs with *assignTaxonomy()* using the SILVA database version 132 as a reference. SILVA was our taxonomy of choice because it is a relatively large 16S sequence database compared to alternative databases, it is regularly maintained and updated and it is widely used in ecological research, making our results comparable to other 16S studies. (Balvočiūtė and Huson, 2017)

Taxonomic training data formatted for DADA2 (silva_nr_v132_train_set.fa.gz) was obtained from<https://zenodo.org/record/1172783#.XhN6UxdKh24>, as referenced by<https://benjjneb.github.io/dada2/training.html> on GitHub. 16S reads and sample data were prepared in an R Phyloseq object for further processing.

ASV table filtering

Raw ASV reads were subjected to a series of filters to produce a final ASV table with biologically relevant 16S sequences: 1) Removed singleton 16S reads. 2) removed sequences that did not map to either Bacteria or Archaea. 3) Removed chloroplast sequences. 4) Removed mitochondrial sequences. 5) Prevalence filter: A range of prevalence filters was tested with x observations in n number of samples.

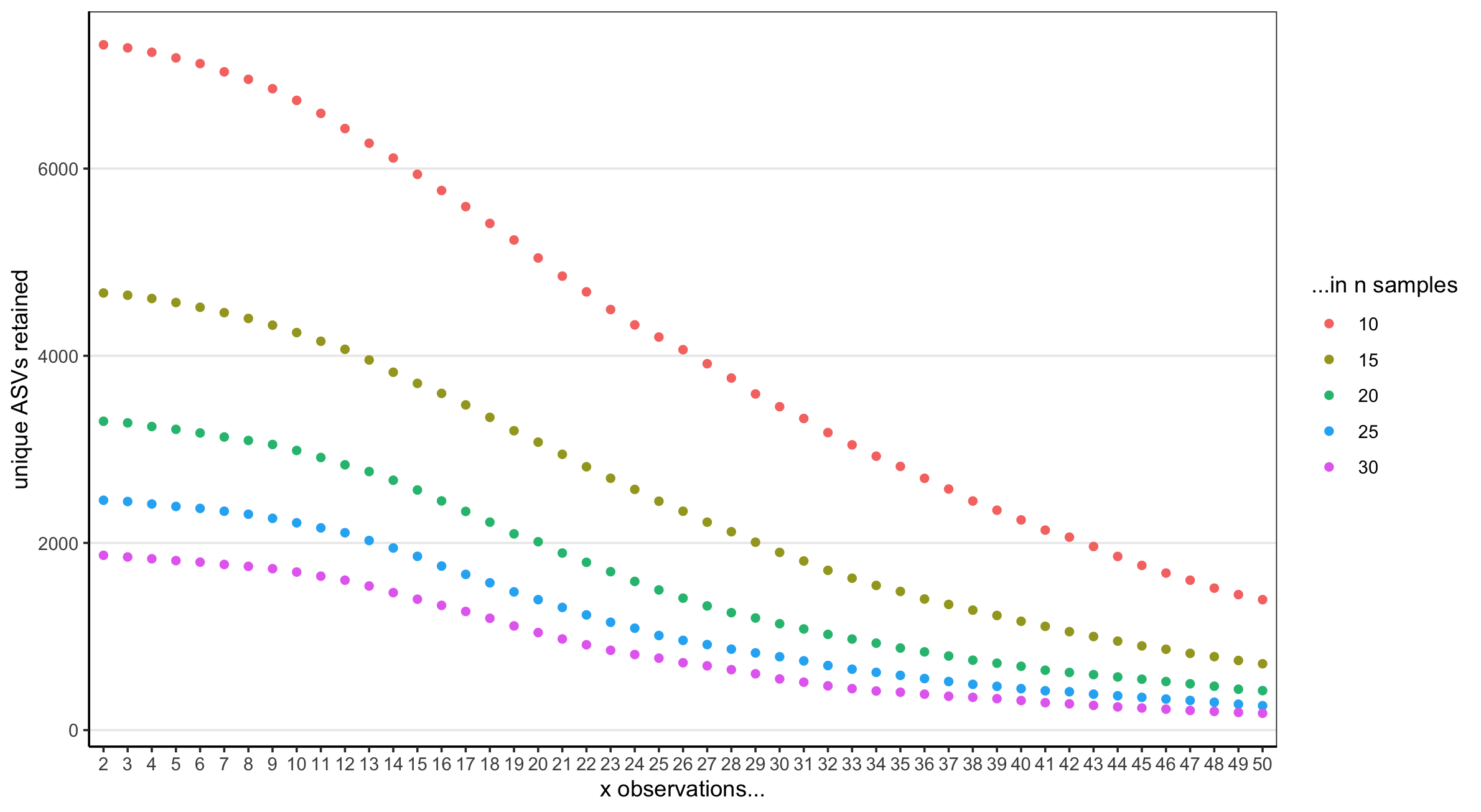

Prevalence filtering of ASV table. To find a workable prevalence filter, various thresholds “x observations in n samples” were tested. We decided to set the threshold to retain ASVs to 10 observations in at least 20 samples right before rapid loss of ASVs is observed (green curve).

To eliminate the vast majority of low-abundance sequences, ASVs were retained that had at least 10 observations in at least 20 samples. 6) Removed any reads that failed to be classified down to the genus level using SILVA taxonomy. This excludes reads where taxonomy is unclear or ambiguous and ensures efficient grouping of ASVs at the genome level later on. 7) Removed any ASVs where log transformed relative abundances were not normally distributed across 384 samples. The ASV table from step 6 was converted to relative abundances and values were transformed with the natural logarithm. Using the R “stats” package, normality was assessed for each ASV individually using the Shapiro-Wilks test shapiro.test(). Shapiro p-values were adjusted with the Benjamini-Hochberg procedure to account for false discovery rate. ASVs were discarded if p_adj < 0.05, meaning a significant difference from a normal distribution. The resulting set of 2225 ASVs was used to generate a phylogenetic tree using mafft v. 7.404 for multiple alignment and fasttree v. 2.1 and the phylogenetic tree was attached to the phyloseq object. Lastly, 11 samples were removed from the data set that had fewer than 1000 total ASV counts. The final ASV table thus contained 2225 ASVs x 373 samples. This table was again converted to relative abundances and log transformed.

### Statistical analysis

Variance partitioning was performed on the above ASV table using R package lme4 using the model log(ASV relative abundance) ~ Year + Month + Host Species + Crop Rotation + Nitrogen + Block + Subsample with all random factors. Subsample with all random factors. Genus subgroups were identified visually by plotting variance partitioning data against phylogenetic trees of ASVs in each genus generated with R packages ggtree and ape. To compare genus subgroup selection to OTUs, FASTA files were generated using R package seqinr from all ASVs in each genus for which we defined subgroups. OTUs were generated from FASTA files using pick_open_reference_otus.py implemented in Qiime/1.9 (Caporaso et al., 2010) with default settings. Differential abundance analysis was performed using R package DESeq2 using the raw ASV table with plain sequence counts (not relative abundance or log transformed) with a +1 pseudocount added to all table values.

### Data Availability

ASV_table, sample data, taxonomy and partitioned variance scores of the set of 2,225 ASVs are available in supplementary data. Scripts used to analyze the data are available on GitHub (<https://github.com/mandmeier/USDA_CornSoy>). 16S sequences are available at (will be updated).
