## Supplementary material for "Rhizosphere Microbiomes in a Historical Maize/Soybean Rotation System respond to Host Species and Nitrogen Fertilization at Genus and Sub-genus Levels": Subgroups of microbial genera

### Identification of sub-genus groups

- Phylogenetic tree of 64 genera with >5 ASVs plotted against variance explained by host species and nitrogen
- Sub-genus groups were identified for 12 genera

### Genera with sub-genus groups

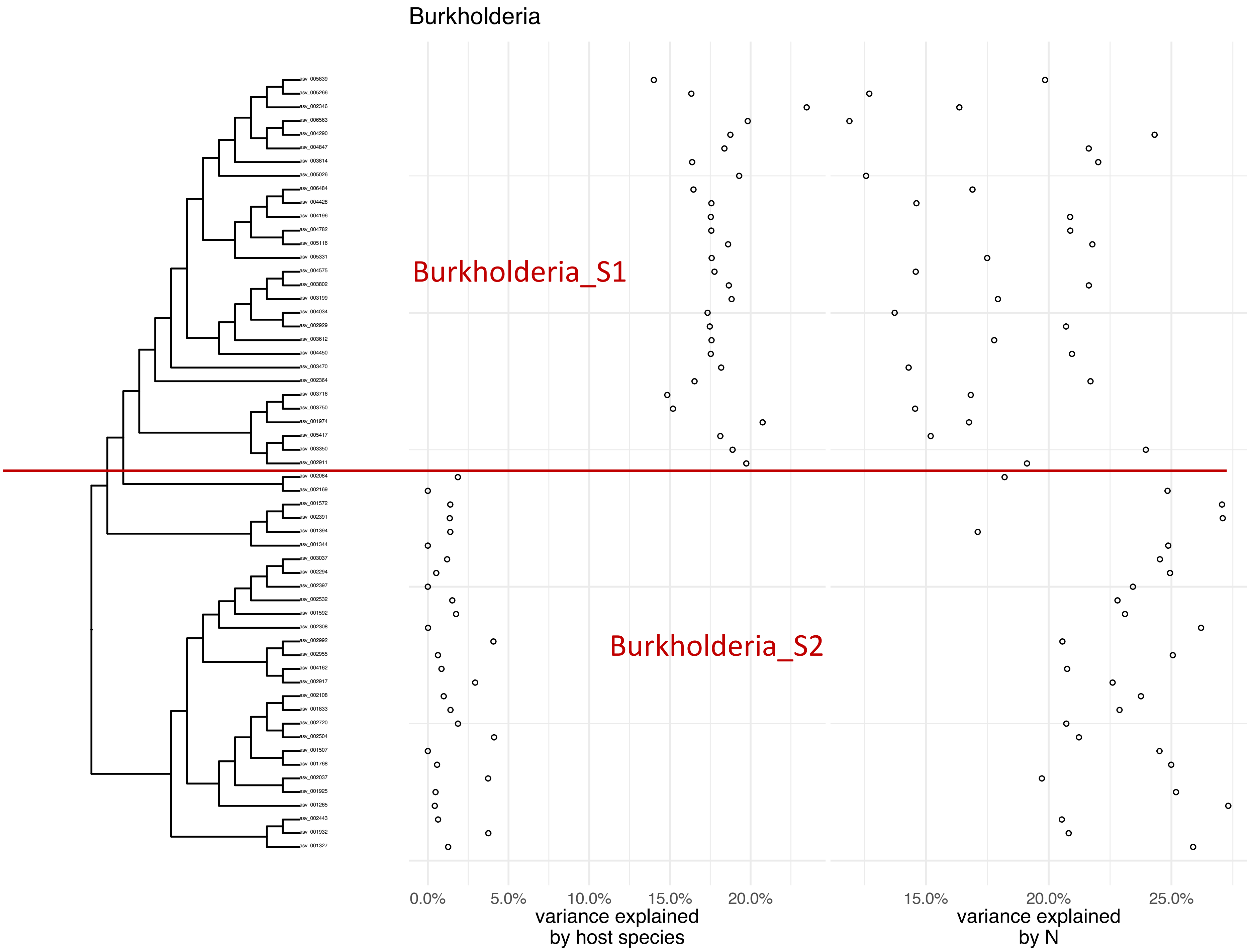

Burkholderia
Burkholderia_S1
Burkholderia_S2
asv_005839
asv_005266
asv_002346
asv_006563
asv_004290
asv_004847
asv_003814
asv_005026
asv_006484
asv_004428
asv_004196
asv_004782
asv_005116
asv_005331
asv_004575
asv_003802
asv_003199
asv_004034
asv_002929
asv_003612
asv_004450
asv_003470
asv_002364
asv_003716
asv_003750
asv_001974
asv_005417
asv_003350
asv_002911
asv_002084
asv_002169
asv_001572
asv_002391
asv_001394
asv_001344
asv_003037
asv_002294
asv_002397
asv_002532
asv_001592
asv_002308
asv_002992
asv_002955
asv_004162
asv_002917
asv_002108
asv_001833
asv_002720
asv_002504
asv_001507
asv_001768
asv_002037
asv_001925
asv_001265
asv_002443
asv_001932
asv_001327
0.0%
5.0%
10.0%
15.0%
20.0%
variance explained
by host species
15.0%
20.0%
25.0%
variance explained
by N

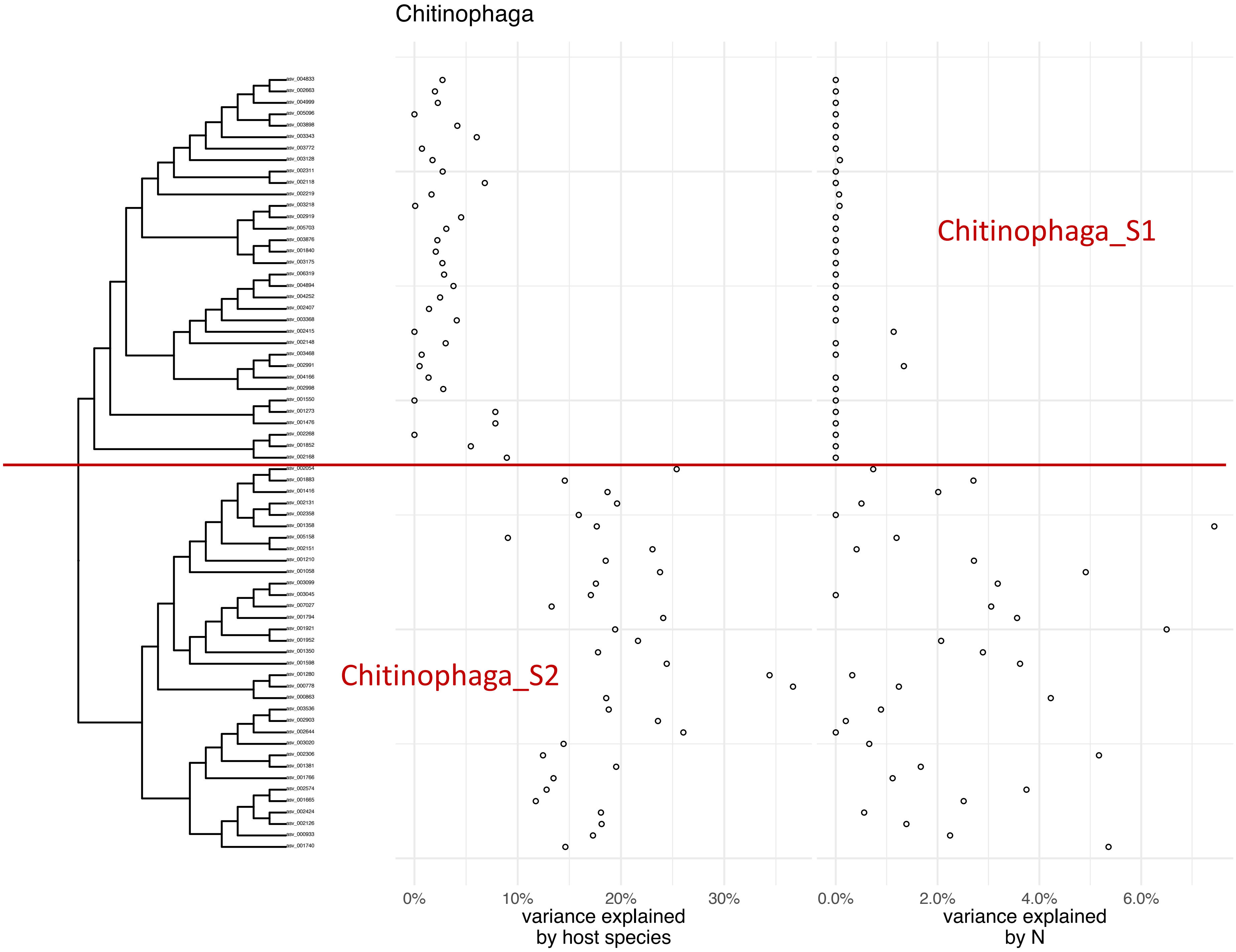
Chitinophaga
asv_004833
asv_002663
asv_004999
asv_005096
asv_003898
asv_003343
asv_003772
asv_003128
asv_002311
asv_002118
asv_002219
asv_003218
asv_002919
asv_005703
asv_003876
asv_001840
asv_003175
asv_006319
asv_004894
asv_004252
asv_002407
asv_003368
asv_002415
asv_002148
asv_003468
asv_002991
asv_004166
asv_002998
asv_001550
asv_001273
asv_001476
asv_002268
asv_001852
asv_002168
asv_002054
asv_001883
asv_001416
asv_002131
asv_002358
asv_001358
asv_005158
asv_002151
asv_001210
asv_001058
asv_003099
asv_003045
asv_007027
asv_001794
asv_001921
asv_001952
asv_001350
asv_001598
asv_001280
asv_000778
asv_000863
asv_003536
asv_002903
asv_002644
asv_003020
asv_002306
asv_001381
asv_001766
asv_002574
asv_001665
asv_002424
asv_002126
asv_000933
asv_001740
Chitinophaga_S1
Chitinophaga_S2
0%
10%
20%
30%
variance explained
by host species
0.0%
2.0%
4.0%
6.0%
variance explained
by N

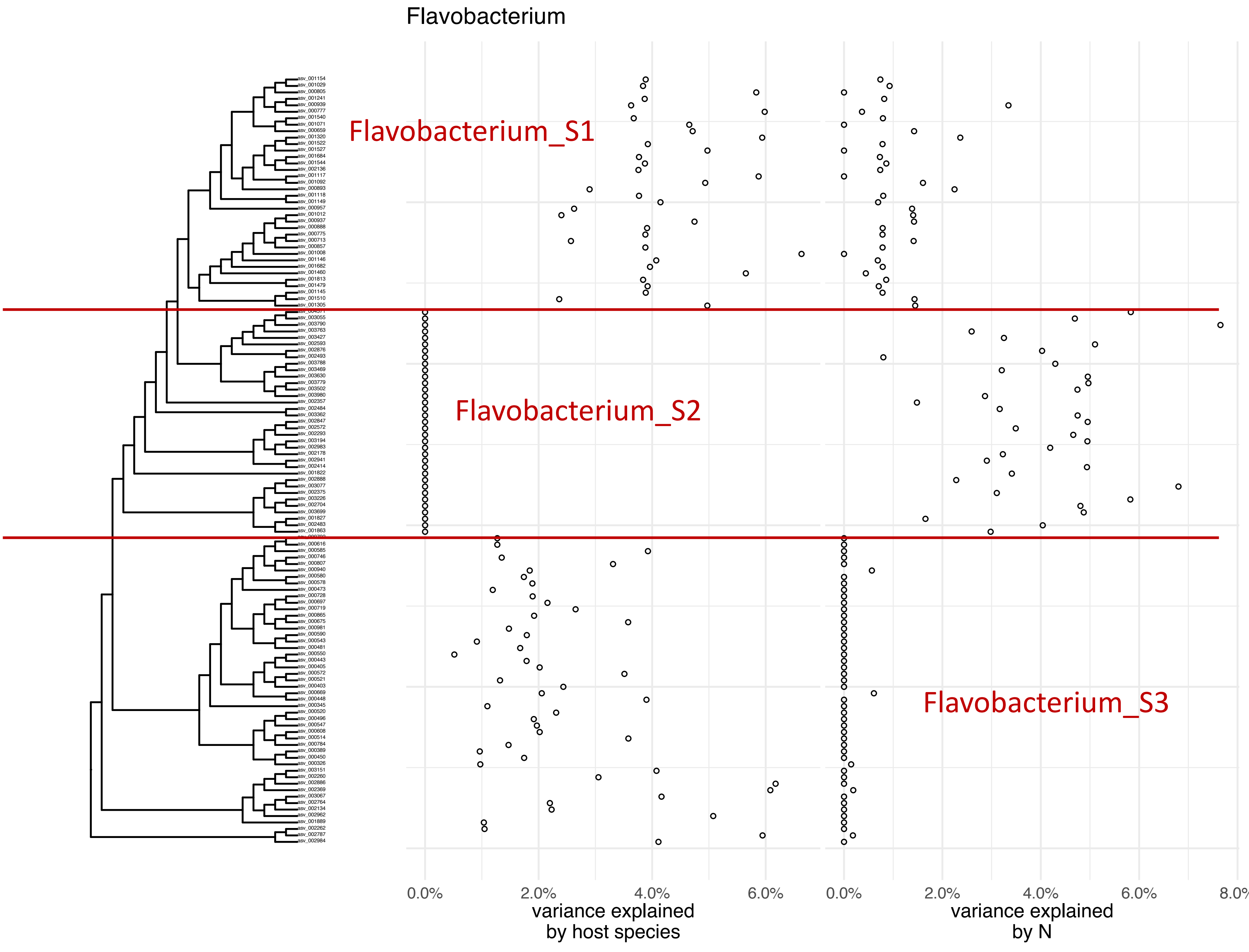
Flavobacterium
Flavobacterium_S1
Flavobacterium_S2
Flavobacterium_S3
0.0%
2.0%
4.0%
6.0%
variance explained
by host species
0.0%
2.0%
4.0%
6.0%
8.0%
variance explained
by N

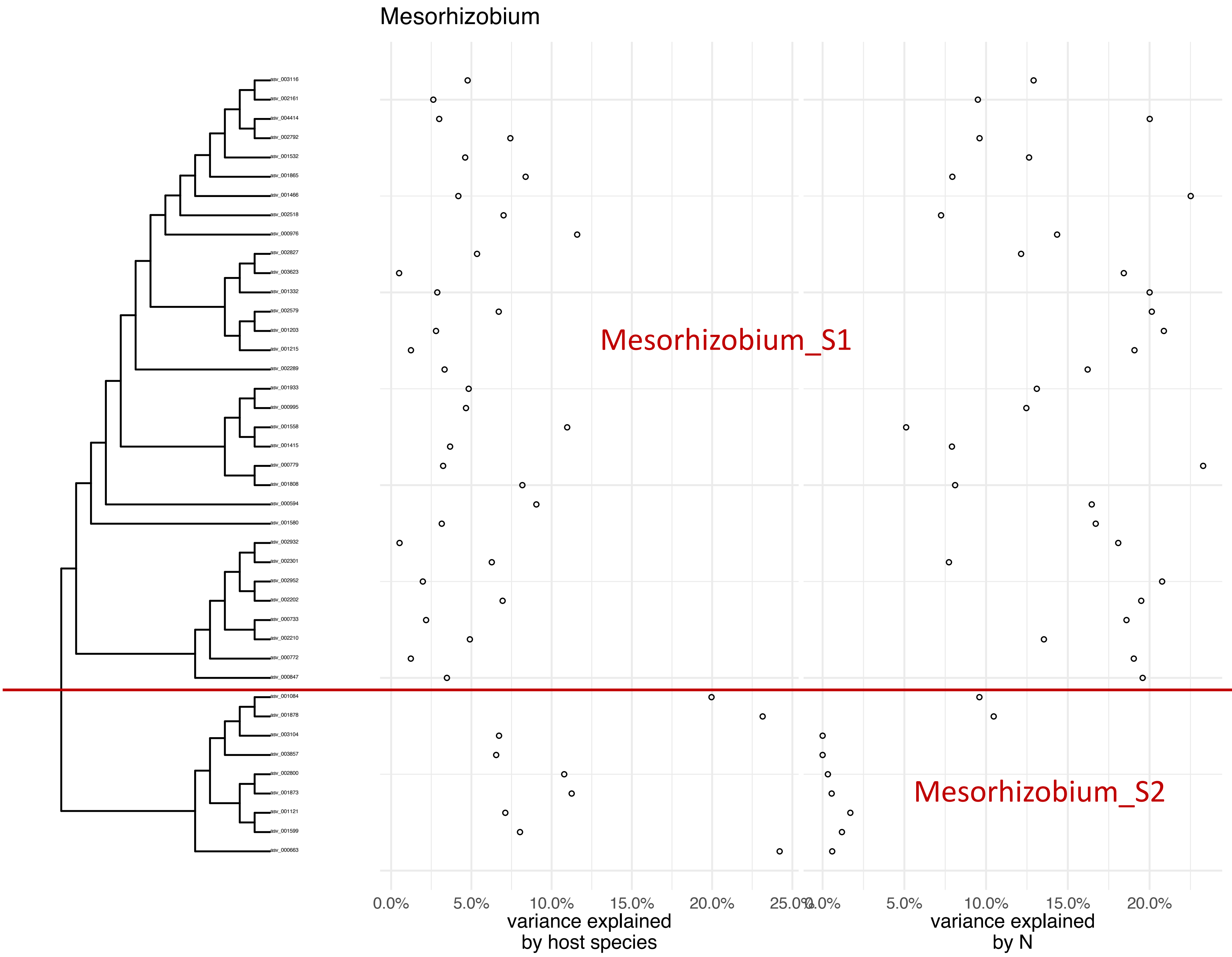
Mesorhizobium
asv_003116
asv_002161
asv_004414
asv_002792
asv_001532
asv_001865
asv_001466
asv_002518
asv_000976
asv_002827
asv_003623
asv_001332
asv_002579
asv_001203
asv_001215
asv_002289
asv_001933
asv_000995
asv_001558
asv_001415
asv_000779
asv_001808
asv_000594
asv_001580
asv_002932
asv_002301
asv_002952
asv_002202
asv_000733
asv_002210
asv_000772
asv_000847
asv_001084
asv_001878
asv_003104
asv_003857
asv_002800
asv_001873
asv_001121
asv_001599
asv_000663
Mesorhizobium_S1
Mesorhizobium_S2
0.0%
5.0%
10.0%
15.0%
20.0%
25.0%
variance explained by host species
0.0%
5.0%
10.0%
15.0%
20.0%
variance explained by N

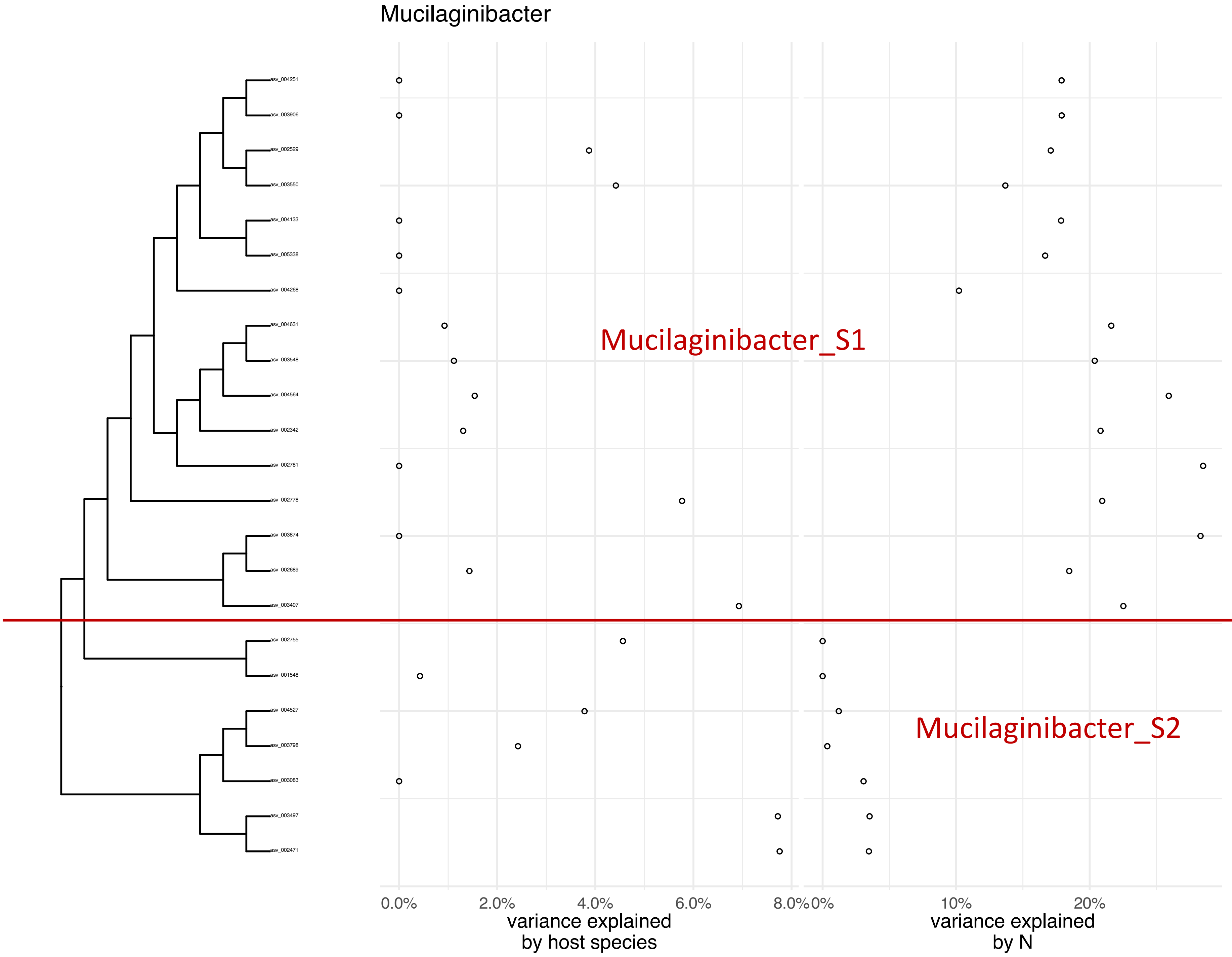
Mucilaginibacter
asv_004251
asv_003906
asv_002529
asv_003550
asv_004133
asv_005338
asv_004268
asv_004631
asv_003548
asv_004564
asv_002342
asv_002781
asv_002778
asv_003874
asv_002689
asv_003407
asv_002755
asv_001548
asv_004527
asv_003798
asv_003083
asv_003497
asv_002471
Mucilaginibacter_S1
Mucilaginibacter_S2
0.0%
2.0%
4.0%
6.0%
8.0%
variance explained by host species
0%
10%
20%
variance explained by N

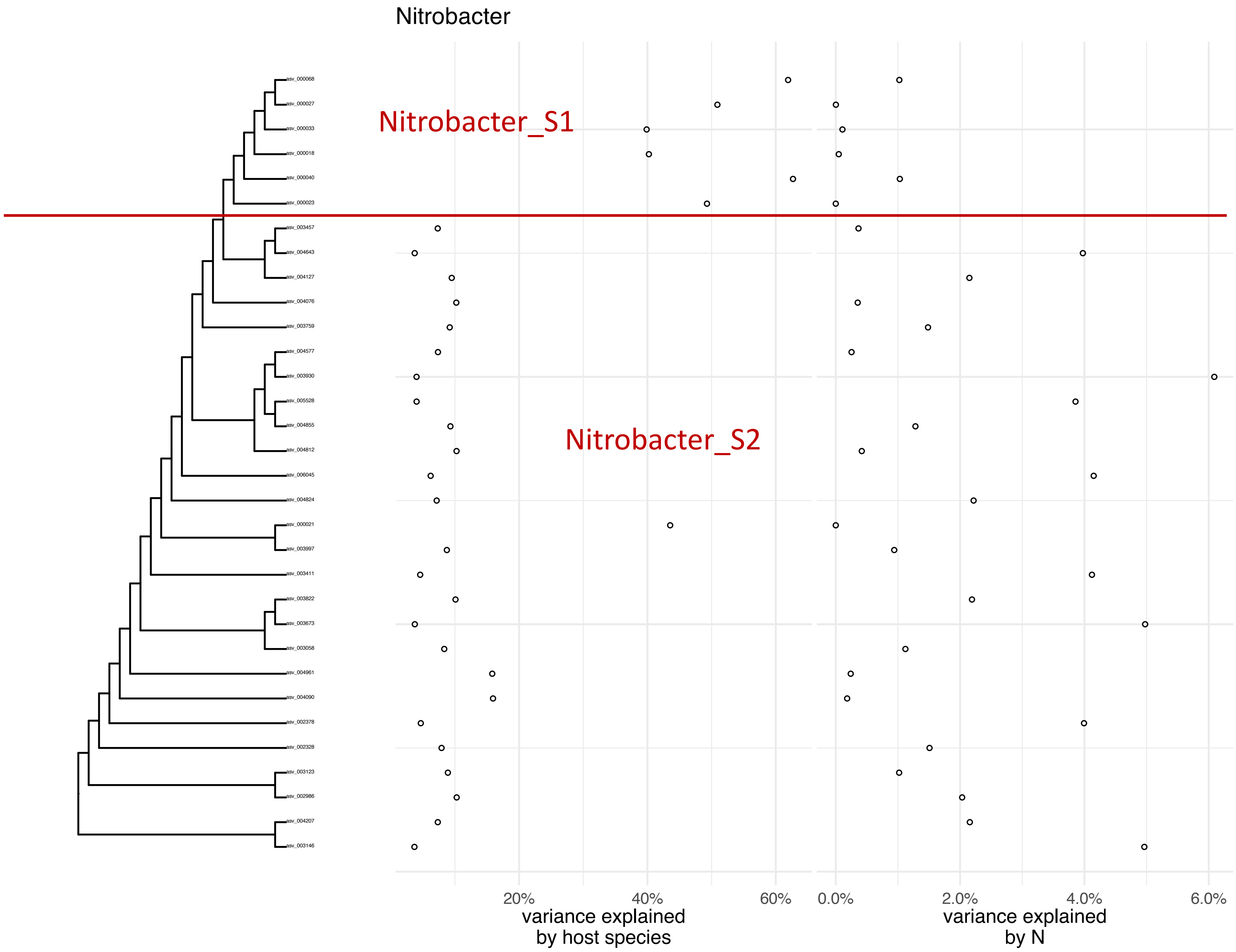
Nitrobacter
Nitrobacter_S1
Nitrobacter_S2
asv_000068
asv_000027
asv_000033
asv_000018
asv_000040
asv_000023
asv_003457
asv_004643
asv_004127
asv_004076
asv_003759
asv_004577
asv_003930
asv_005528
asv_004855
asv_004812
asv_006045
asv_004824
asv_000021
asv_003997
asv_003411
asv_003822
asv_003673
asv_003058
asv_004961
asv_004090
asv_002378
asv_002328
asv_003123
asv_002986
asv_004207
asv_003146
20%
40%
60%
variance explained by host species
0.0%
2.0%
4.0%
6.0%
variance explained by N

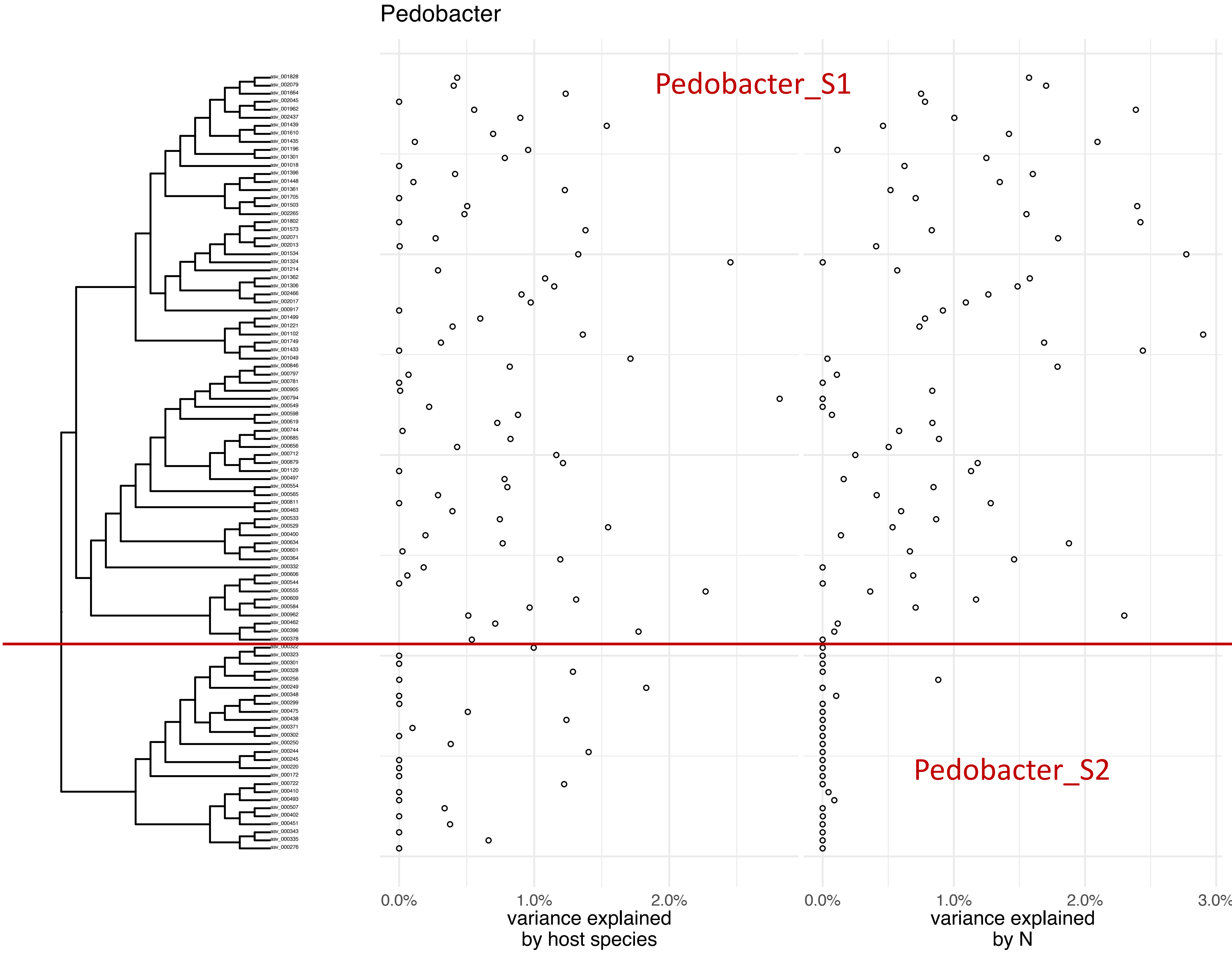

Pedobacter
Pedobacter_S1
Pedobacter_S2
0.0%
1.0%
2.0%
variance explained
by host species
0.0%
1.0%
2.0%
3.0%
variance explained
by N

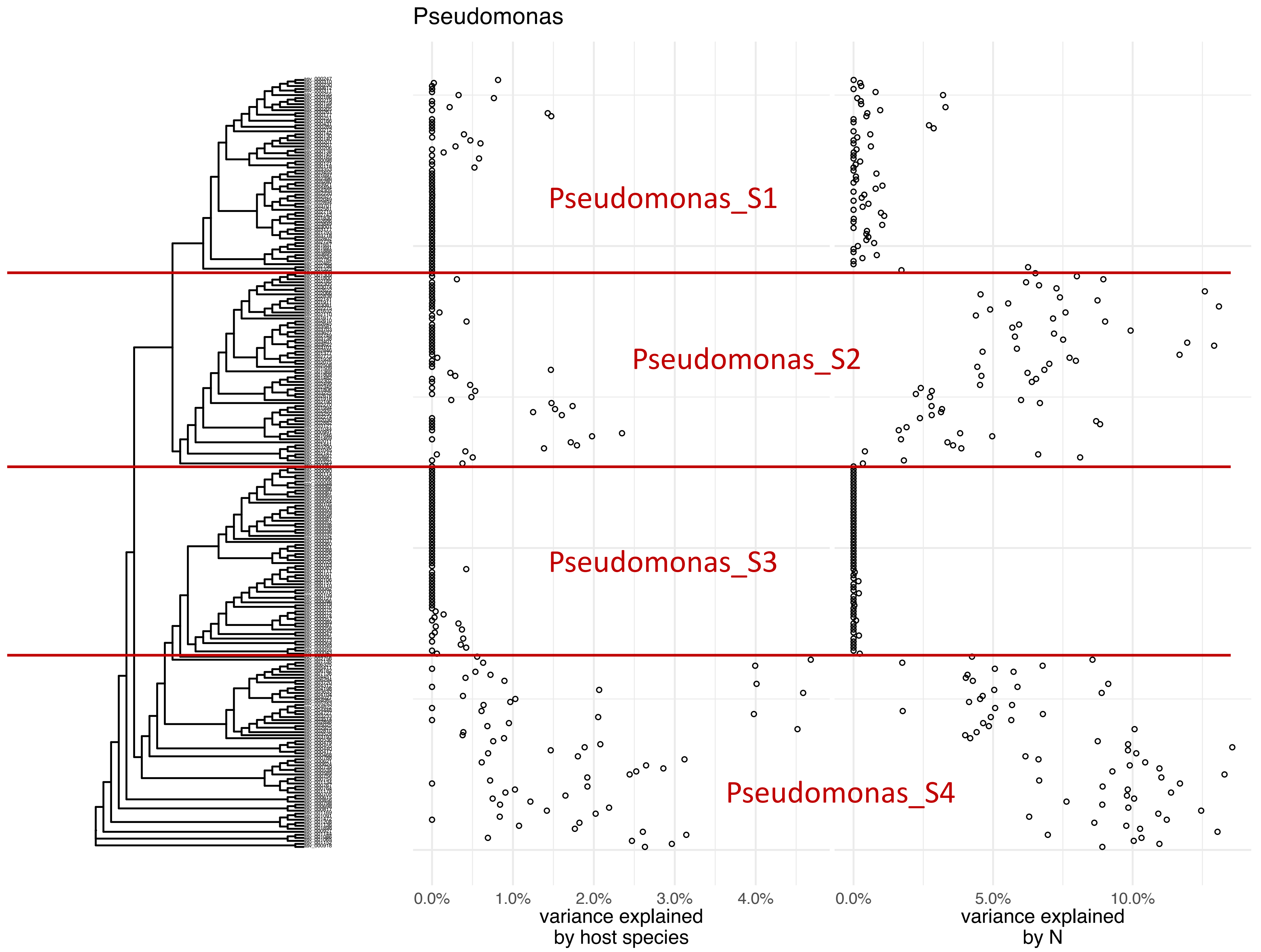

Pseudomonas
Pseudomonas_S1
Pseudomonas_S2
Pseudomonas_S3
Pseudomonas_S4
0.0%
1.0%
2.0%
3.0%
4.0%
variance explained by host species
0.0%
5.0%
10.0%
variance explained by N

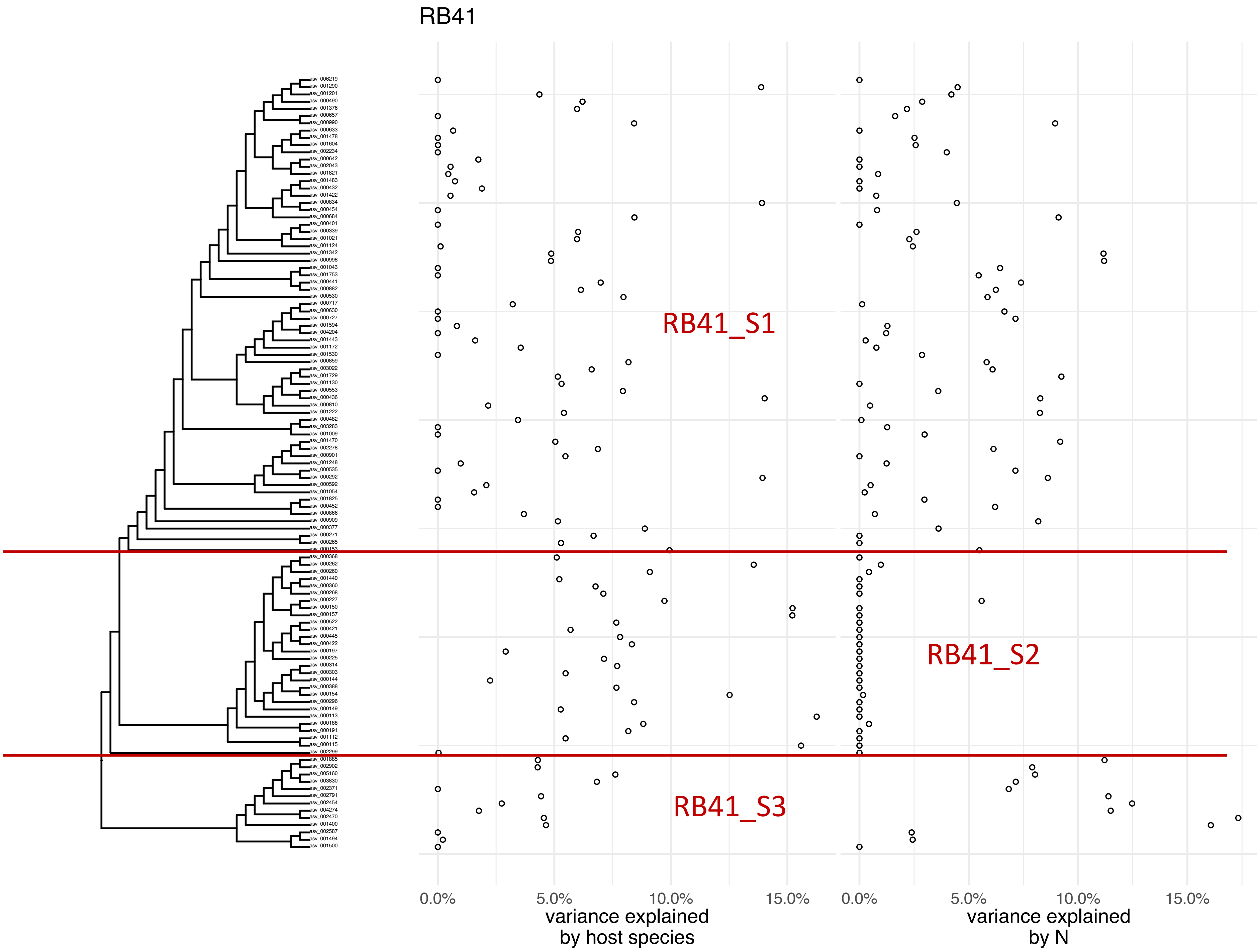
RB41
RB41_S1
RB41_S2
RB41_S3
0.0%
5.0%
10.0%
15.0%
variance explained
by host species
0.0%
5.0%
10.0%
15.0%
variance explained
by N

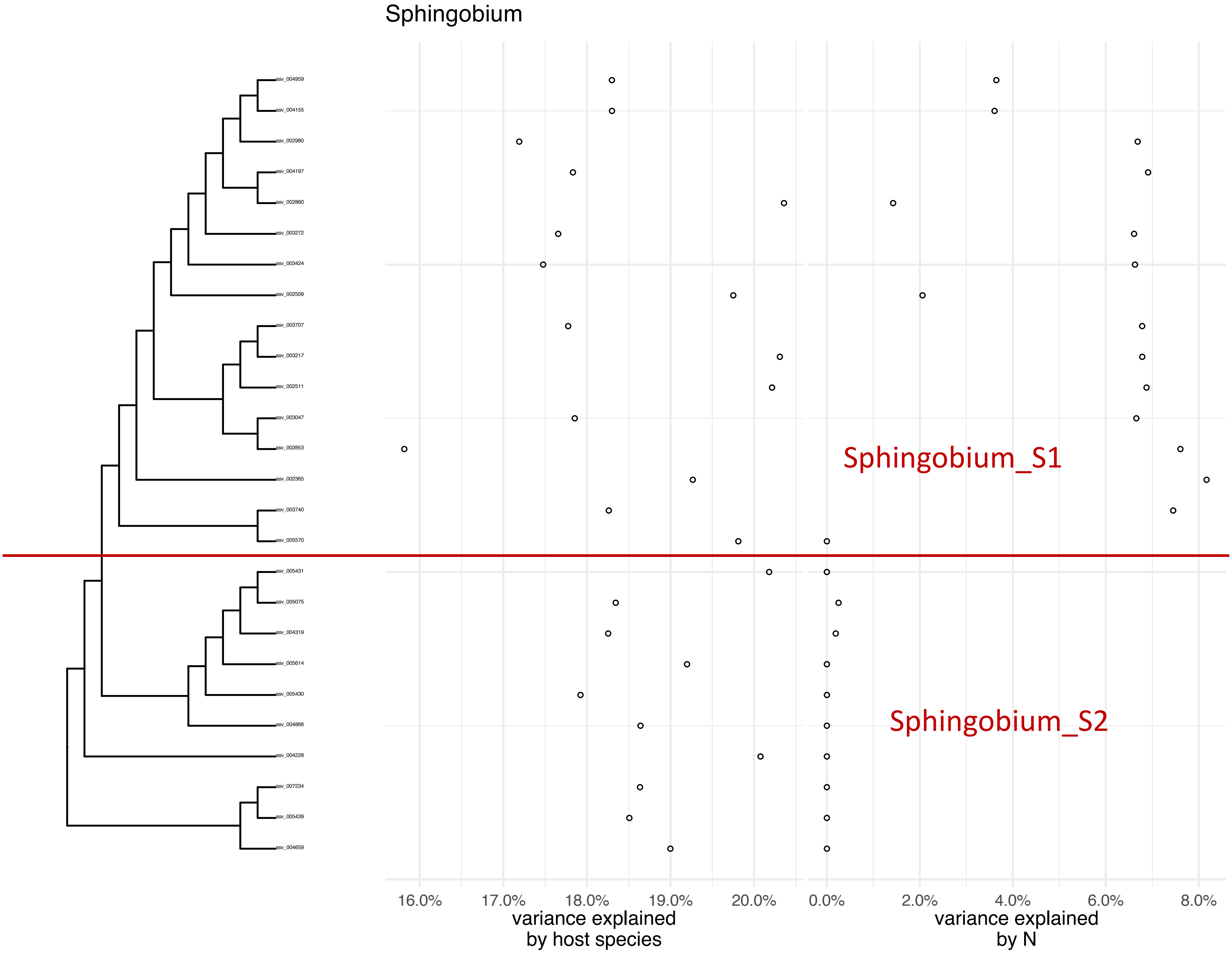
Sphingobium
asv_004959
asv_004155
asv_002980
asv_004197
asv_002860
asv_003272
asv_003424
asv_002509
asv_003707
asv_003217
asv_002511
asv_003047
asv_002853
asv_002365
asv_003740
asv_005570
asv_005431
asv_005075
asv_004319
asv_005614
asv_005430
asv_004866
asv_004228
asv_007234
asv_005439
asv_004659
Sphingobium_S1
Sphingobium_S2
16.0%
17.0%
18.0%
19.0%
20.0%
variance explained
by host species
0.0%
2.0%
4.0%
6.0%
8.0%
variance explained
by N

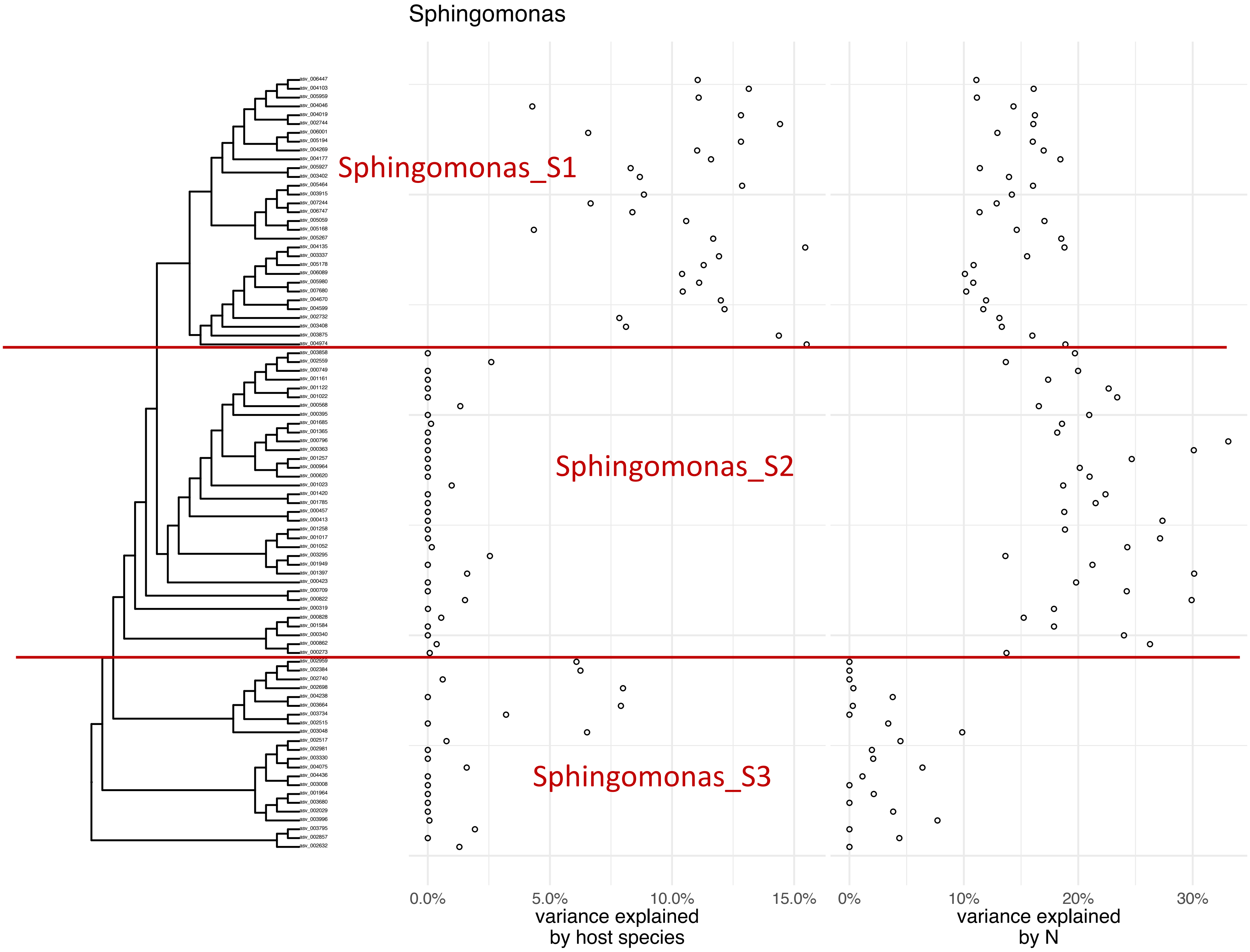
Sphingomonas
Sphingomonas_S1
Sphingomonas_S2
Sphingomonas_S3
0.0%
5.0%
10.0%
15.0%
variance explained
by host species
0%
10%
20%
30%
variance explained
by N

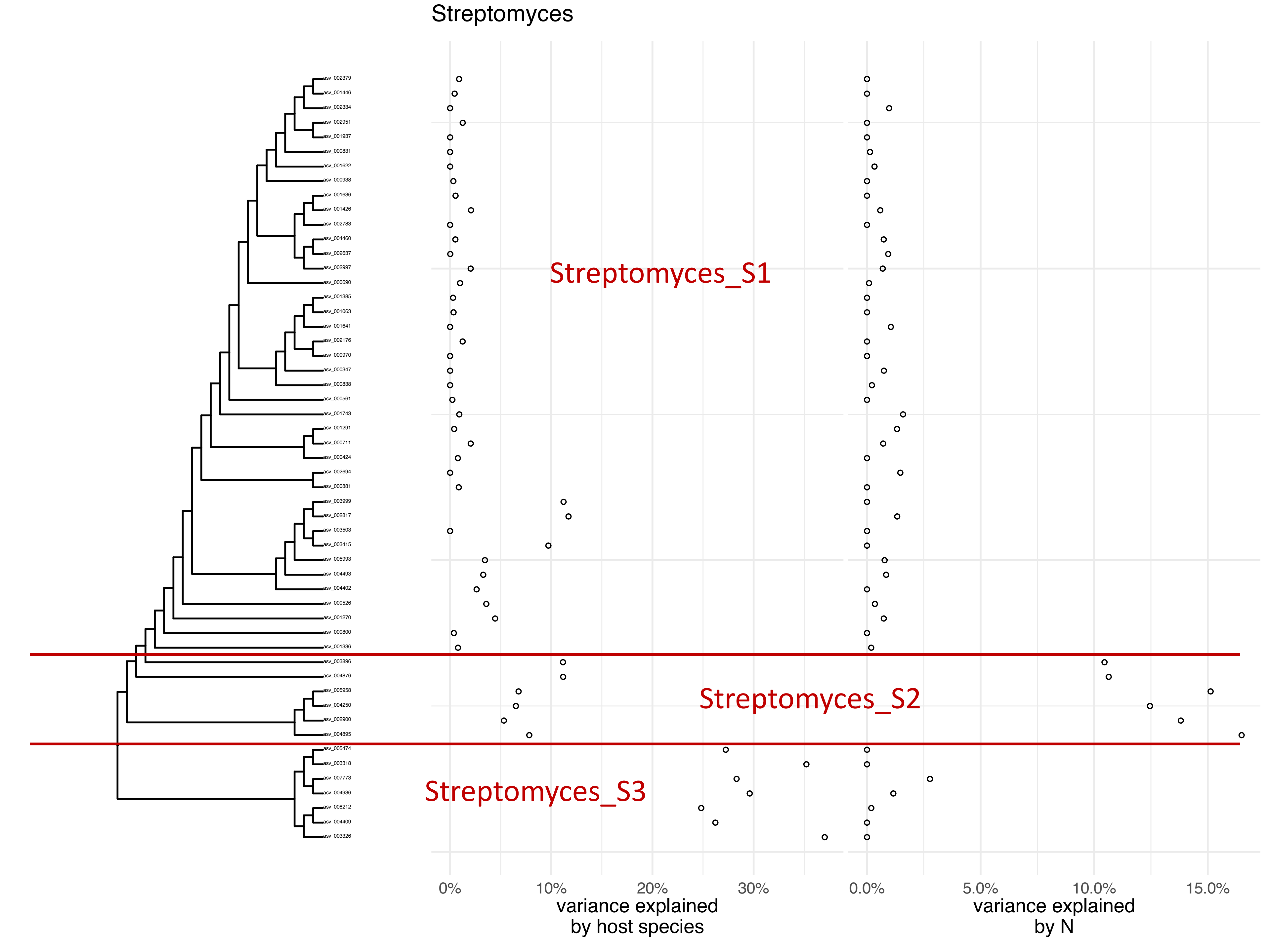
Streptomyces
asv_002379
asv_001446
asv_002334
asv_002951
asv_001937
asv_000831
asv_001622
asv_000938
asv_001636
asv_001426
asv_002783
asv_004460
asv_002637
asv_002997
asv_000690
asv_001385
asv_001063
asv_001641
asv_002176
asv_000970
asv_000347
asv_000838
asv_000561
asv_001743
asv_001291
asv_000711
asv_000424
asv_002694
asv_000881
asv_003999
asv_002817
asv_003503
asv_003415
asv_005993
asv_004493
asv_004402
asv_000526
asv_001270
asv_000800
asv_001336
asv_003896
asv_004876
asv_005958
asv_004250
asv_002900
asv_004895
asv_005474
asv_003318
asv_007773
asv_004936
asv_008212
asv_004409
asv_003326
Streptomyces_S1
Streptomyces_S2
Streptomyces_S3
0%
10%
20%
30%
variance explained by host species
0.0%
5.0%
10.0%
15.0%
variance explained by N

### Genera without sub-genus groups

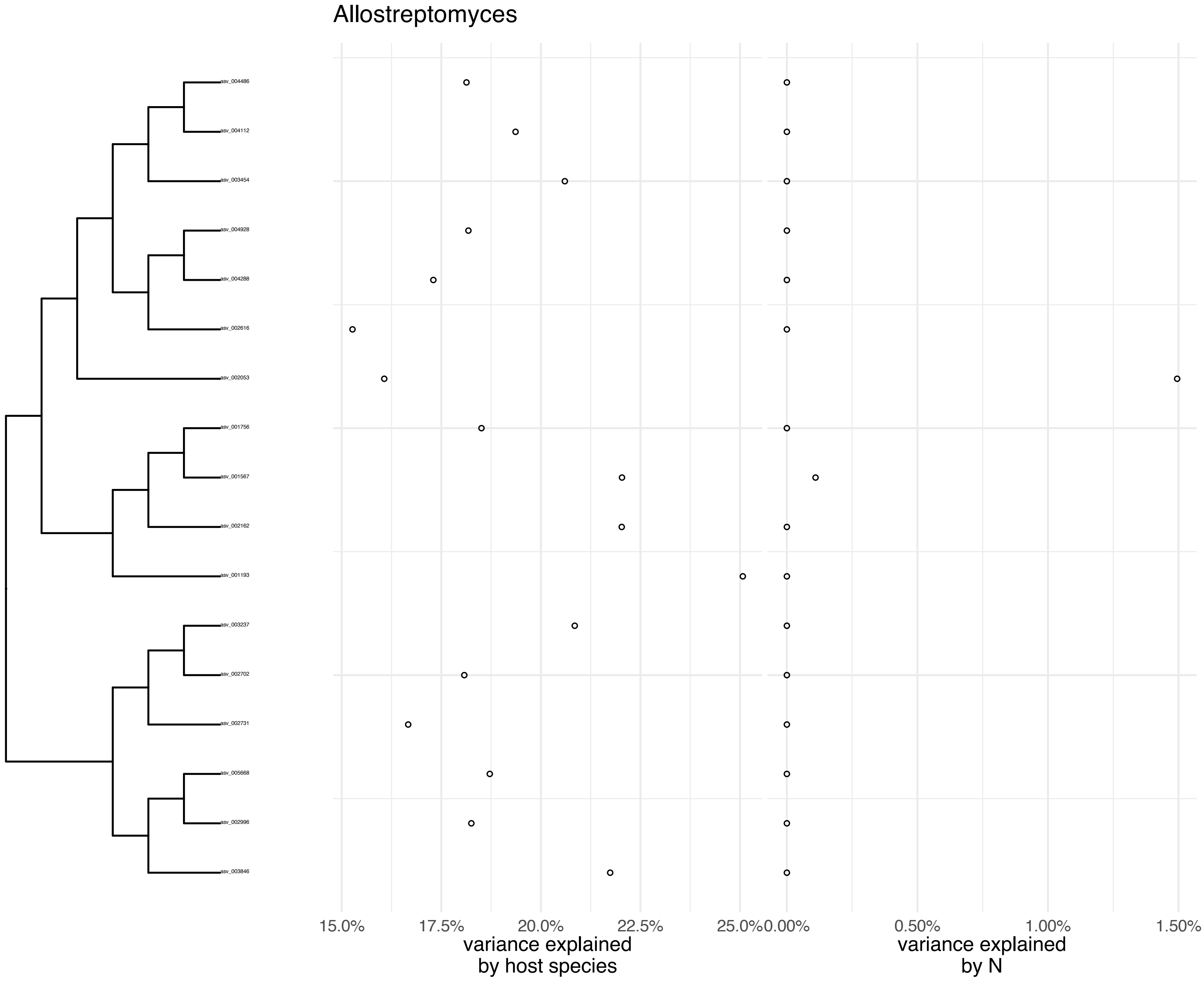
Allostreptomyces
asv_004486
asv_004112
asv_003454
asv_004928
asv_004288
asv_002616
asv_002053
asv_001756
asv_001567
asv_002162
asv_001193
asv_003237
asv_002702
asv_002731
asv_005668
asv_002996
asv_003846
15.0%
17.5%
20.0%
22.5%
25.0%
0.00%
0.50%
1.00%
1.50%
variance explained
by host species
variance explained
by N

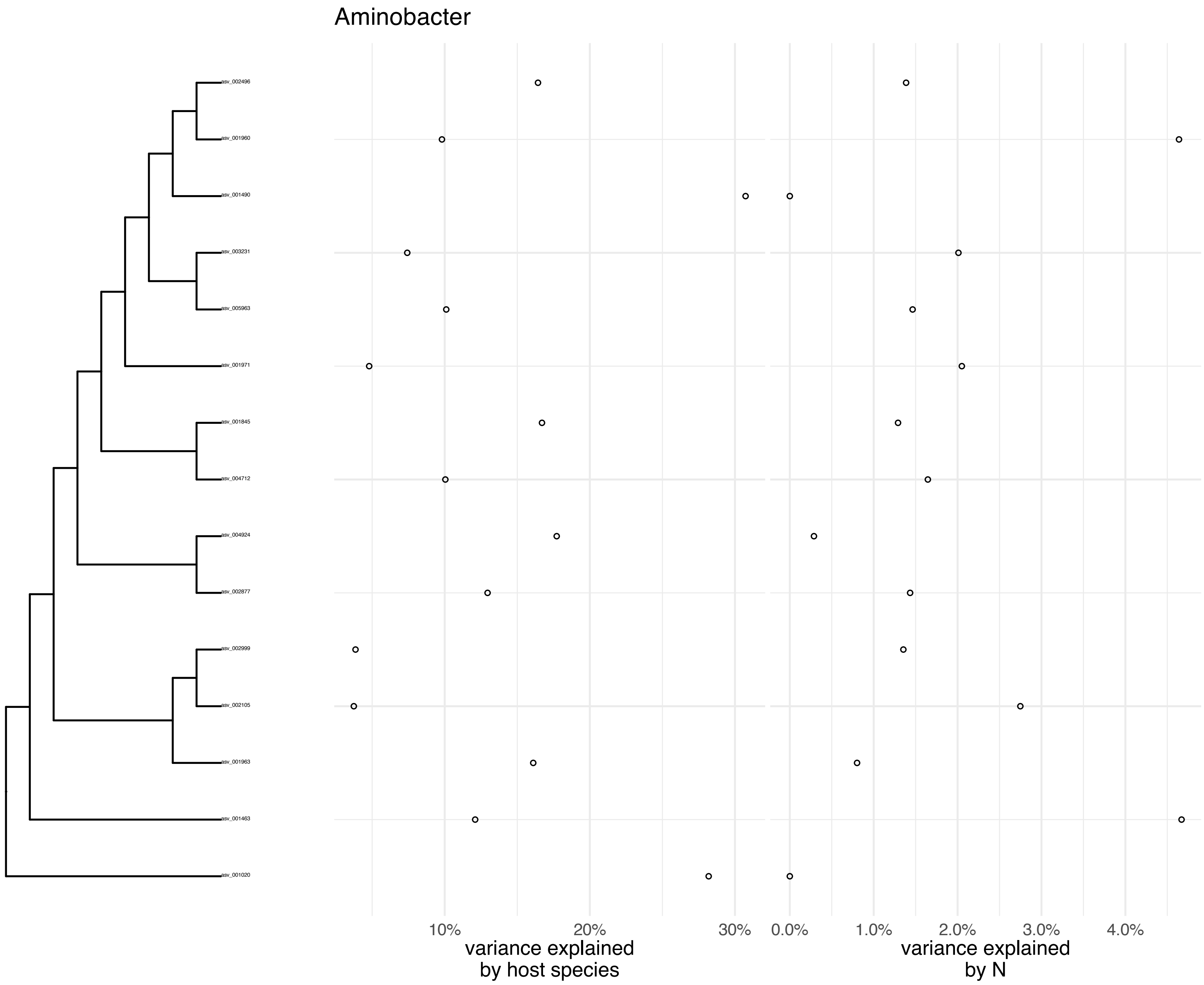
Aminobacter
asv_002496
asv_001960
asv_001490
asv_003231
asv_005963
asv_001971
asv_001845
asv_004712
asv_004924
asv_002877
asv_002999
asv_002105
asv_001963
asv_001463
asv_001020
10%
20%
30%
variance explained by host species
0.0%
1.0%
2.0%
3.0%
4.0%
variance explained by N

Archangium

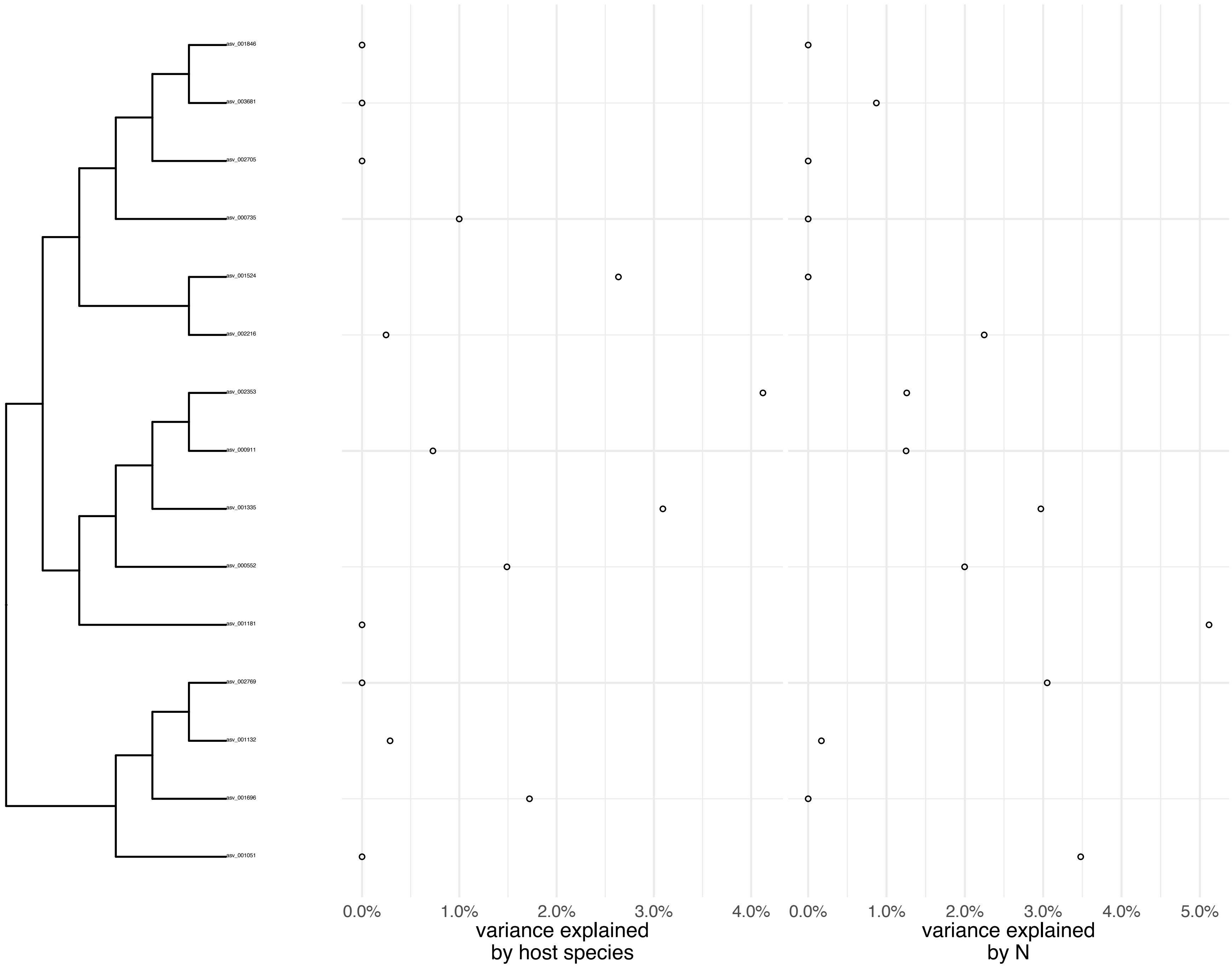
asv_001846
asv_003681
asv_002705
asv_000735
asv_001524
asv_002216
asv_002353
asv_000911
asv_001335
asv_000552
asv_001181
asv_002769
asv_001132
asv_001696
asv_001051
0.0%
1.0%
2.0%
3.0%
4.0%
variance explained by host species
0.0%
1.0%
2.0%
3.0%
4.0%
5.0%
variance explained by N

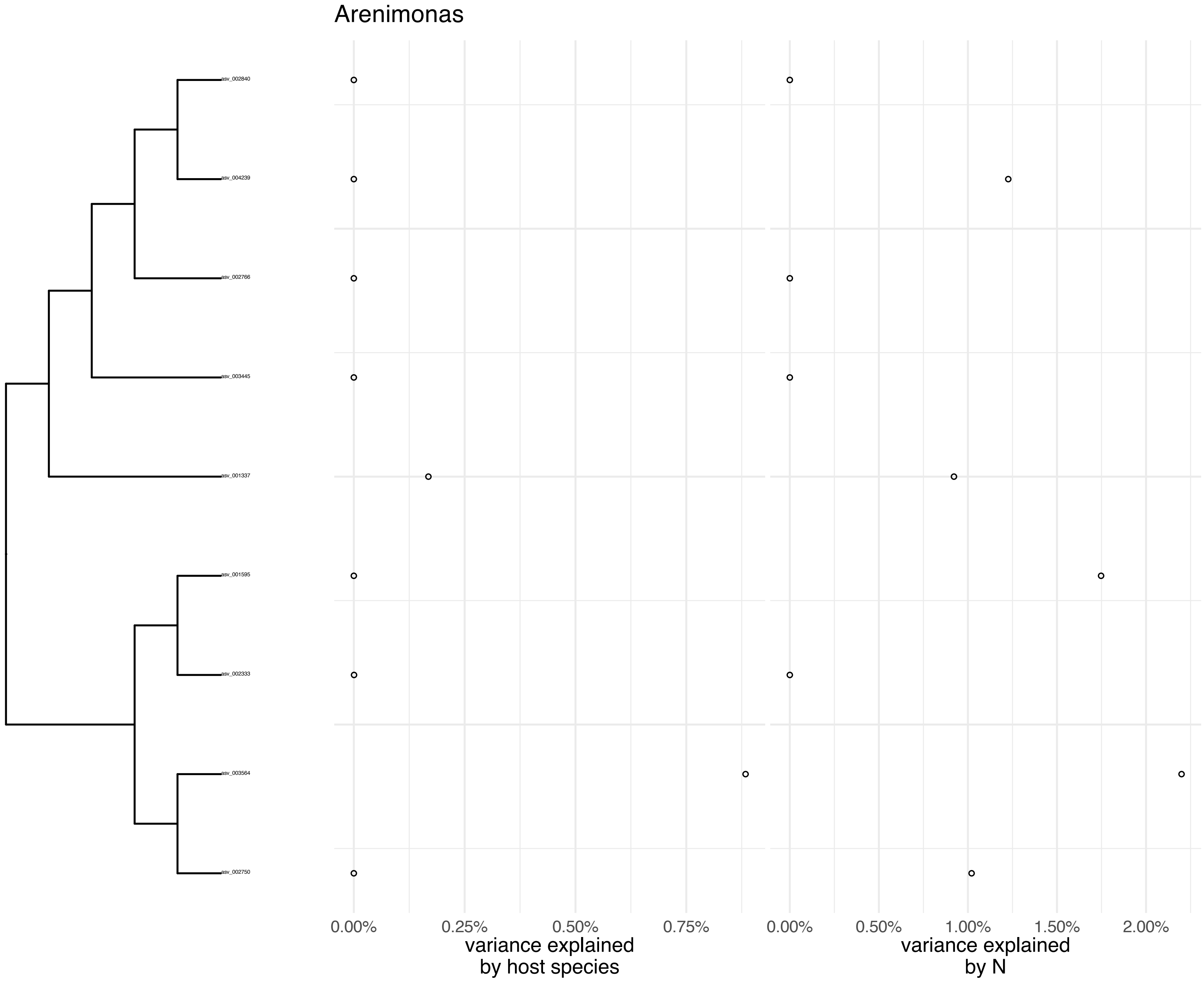

Arenimonas
asv_002840
asv_004239
asv_002766
asv_003445
asv_001337
asv_001595
asv_002333
asv_003564
asv_002750
0.00%
0.25%
0.50%
0.75%
variance explained
by host species
0.00%
0.50%
1.00%
1.50%
2.00%
variance explained
by N

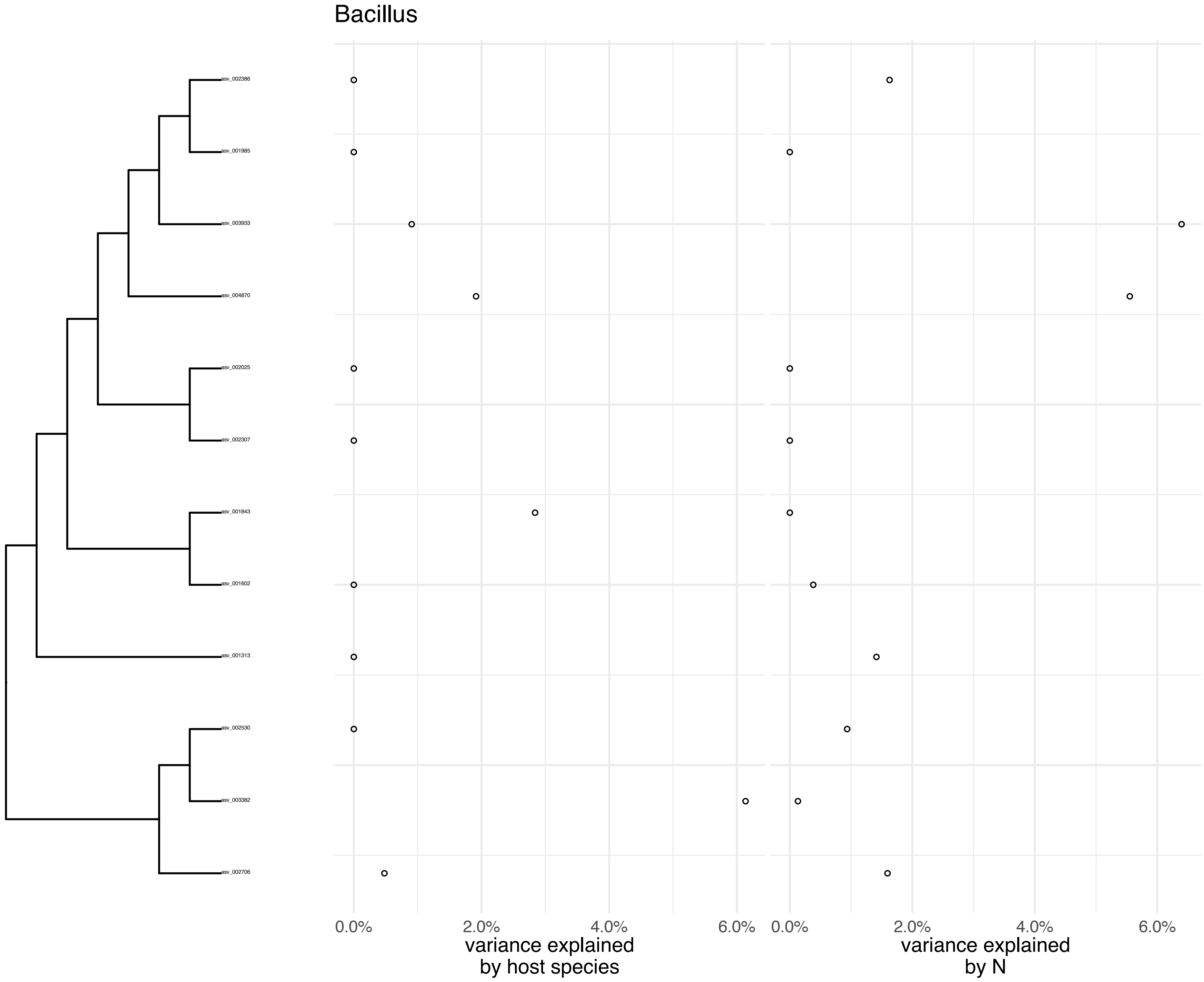
Bacillus
asv_002386
asv_001985
asv_003933
asv_004870
asv_002025
asv_002307
asv_001843
asv_001602
asv_001313
asv_002530
asv_003382
asv_002706
0.0%
2.0%
4.0%
6.0%
variance explained by host species
0.0%
2.0%
4.0%
6.0%
variance explained by N

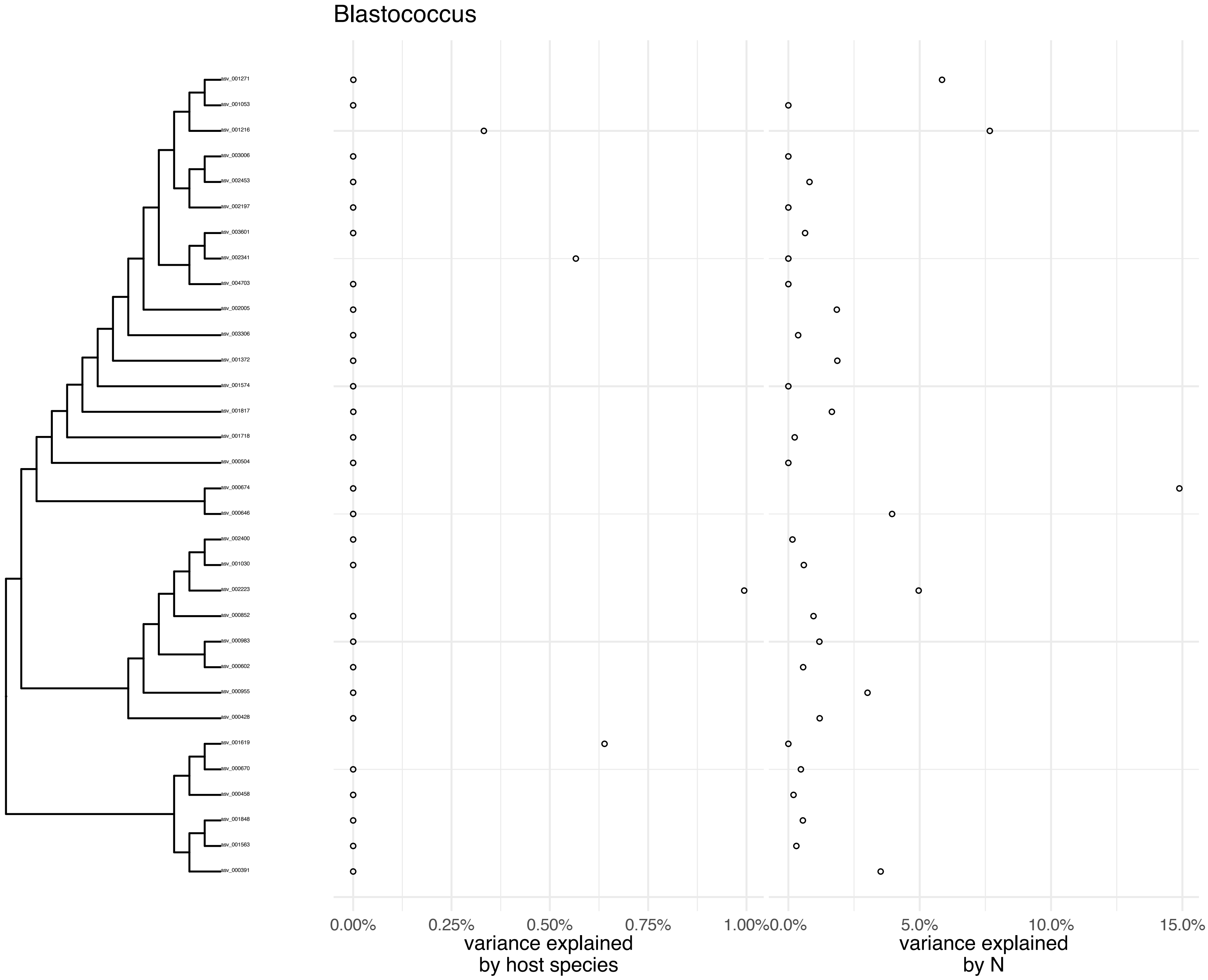
Blastococcus
asv_001271
asv_001053
asv_001216
asv_003006
asv_002453
asv_002197
asv_003601
asv_002341
asv_004703
asv_002005
asv_003306
asv_001372
asv_001574
asv_001817
asv_001718
asv_000504
asv_000674
asv_000646
asv_002400
asv_001030
asv_002223
asv_000852
asv_000983
asv_000602
asv_000955
asv_000428
asv_001619
asv_000670
asv_000458
asv_001848
asv_001563
asv_000391
0.00%
0.25%
0.50%
0.75%
1.00%
variance explained by host species
0.0%
5.0%
10.0%
15.0%
variance explained by N

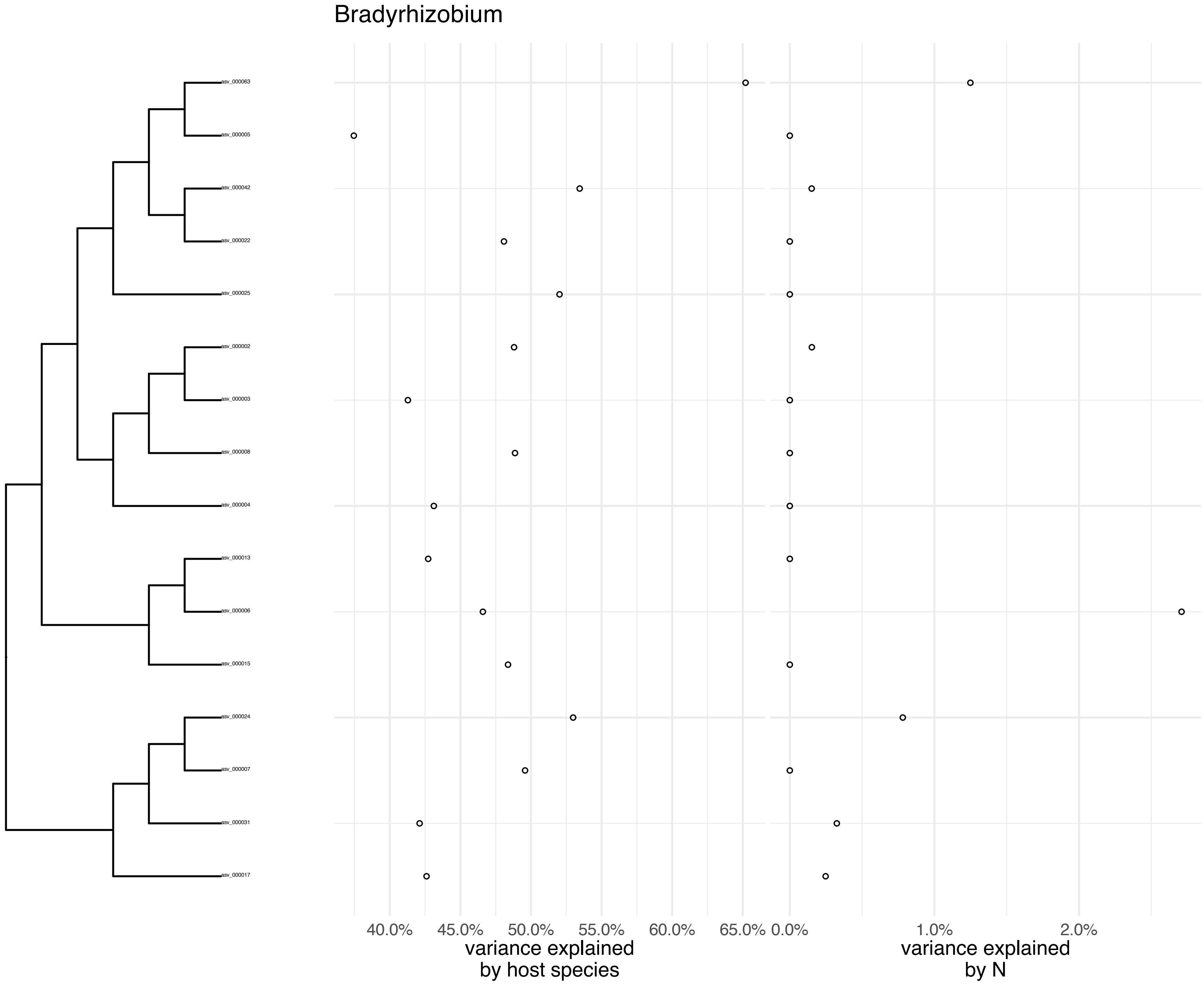

Bradyrhizobium
asv_000063
asv_000005
asv_000042
asv_000022
asv_000025
asv_000002
asv_000003
asv_000008
asv_000004
asv_000013
asv_000006
asv_000015
asv_000024
asv_000007
asv_000031
asv_000017
40.0%
45.0%
50.0%
55.0%
60.0%
65.0%
0.0%
1.0%
2.0%
variance explained by host species
variance explained by N

Candidatus_Nitrocosmicus

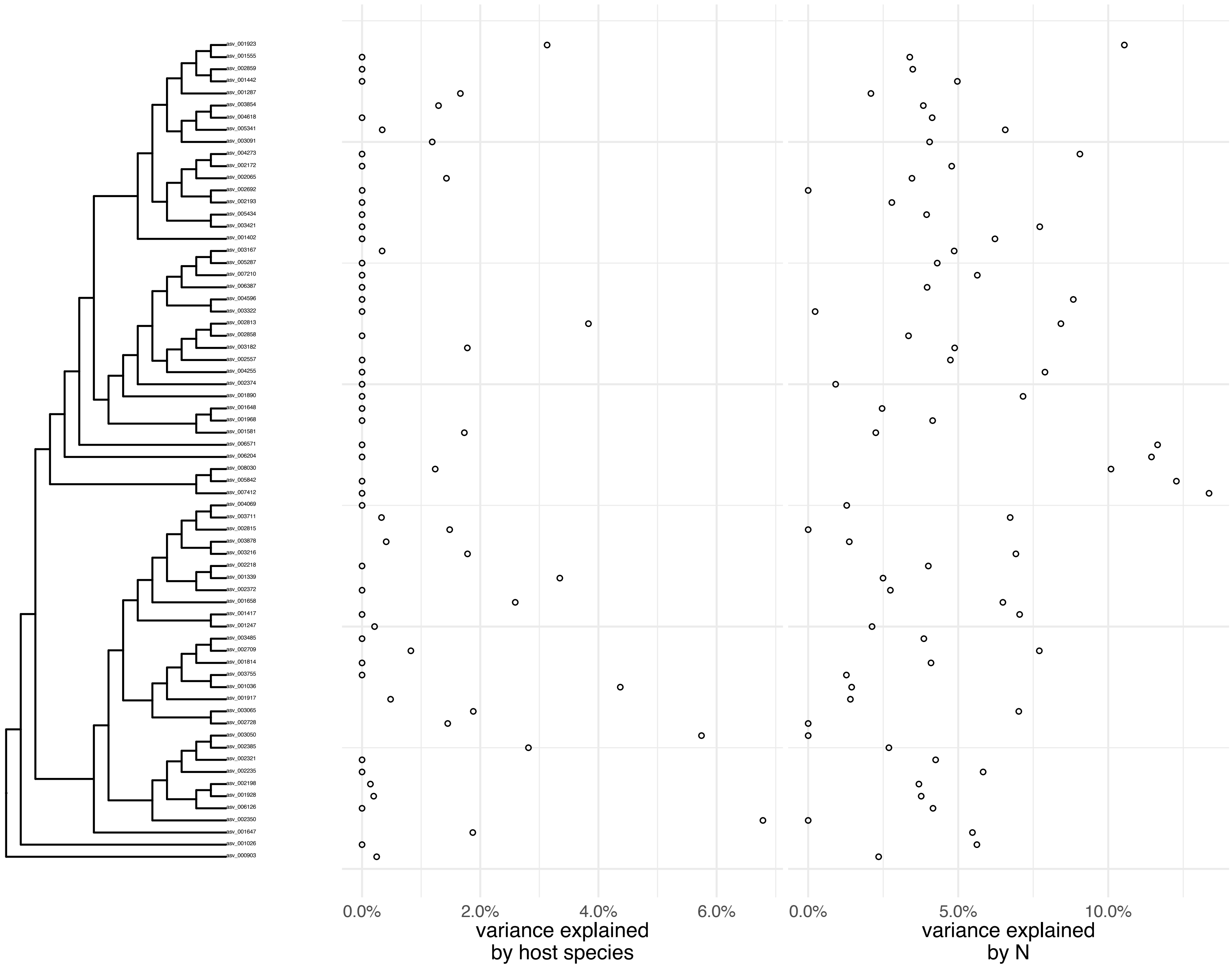

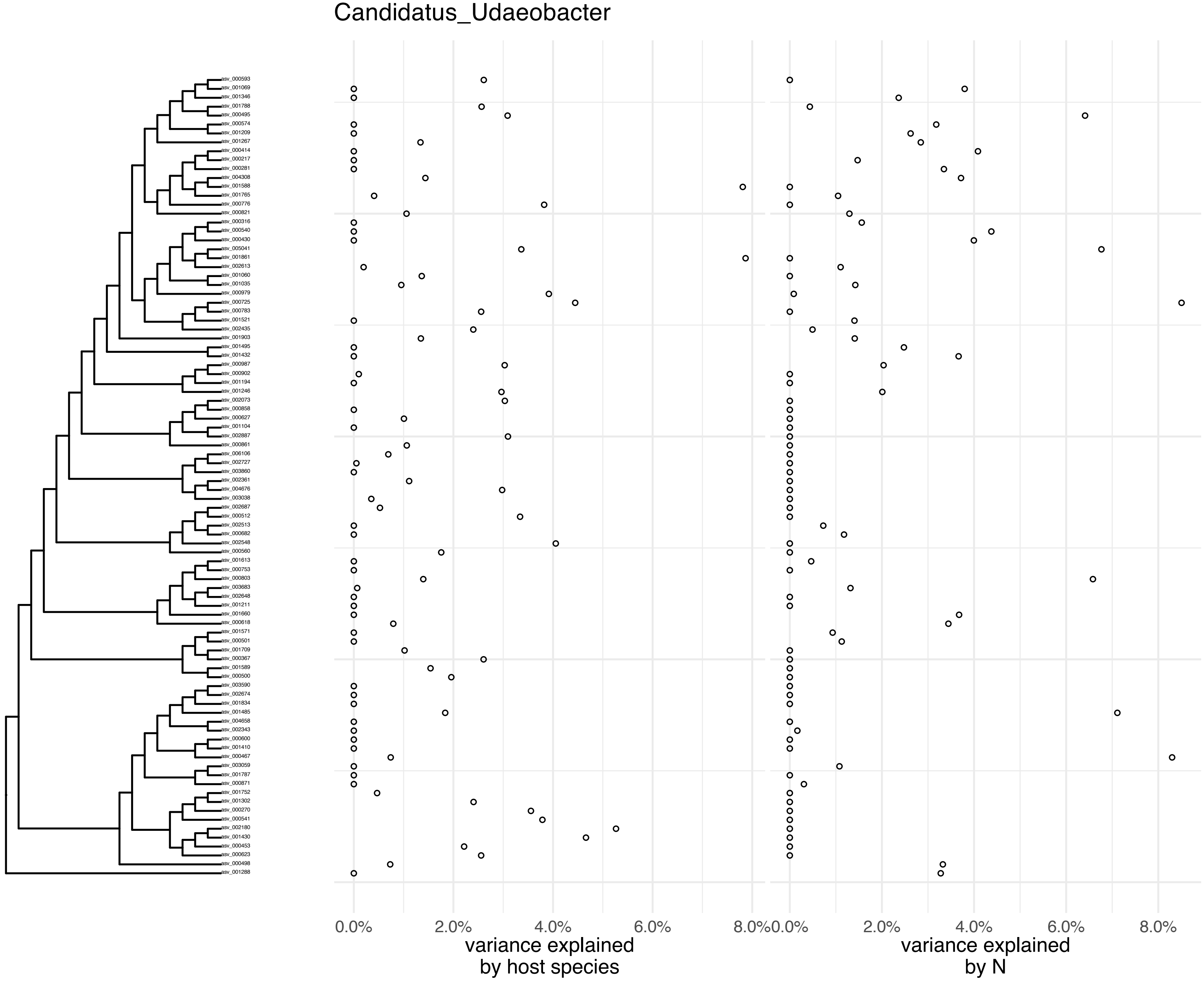

Candidatus_Udaeobacter
asv_000593
asv_001069
asv_001346
asv_001788
asv_000495
asv_000574
asv_001209
asv_001267
asv_000414
asv_000217
asv_000281
asv_004308
asv_001588
asv_001765
asv_000776
asv_000821
asv_000316
asv_000540
asv_000430
asv_005041
asv_001861
asv_002613
asv_001060
asv_001035
asv_000979
asv_000725
asv_000783
asv_001521
asv_002435
asv_001903
asv_001495
asv_001432
asv_000987
asv_000902
asv_001194
asv_001246
asv_002073
asv_000858
asv_000627
asv_001104
asv_002887
asv_000861
asv_006106
asv_002727
asv_003860
asv_002361
asv_004676
asv_003038
asv_002687
asv_000512
asv_002513
asv_000682
asv_002548
asv_000560
asv_001613
asv_000753
asv_000803
asv_003683
asv_002648
asv_001211
asv_001660
asv_000618
asv_001571
asv_000501
asv_001709
asv_000367
asv_001589
asv_000500
asv_003590
asv_002674
asv_001834
asv_001485
asv_004658
asv_002343
asv_000600
asv_001410
asv_000467
asv_003059
asv_001787
asv_000871
asv_001752
asv_001302
asv_000270
asv_000541
asv_002180
asv_001430
asv_000453
asv_000623
asv_000498
asv_001288
0.0%
2.0%
4.0%
6.0%
8.0%
variance explained
by host species
0.0%
2.0%
4.0%
6.0%
8.0%
variance explained
by N

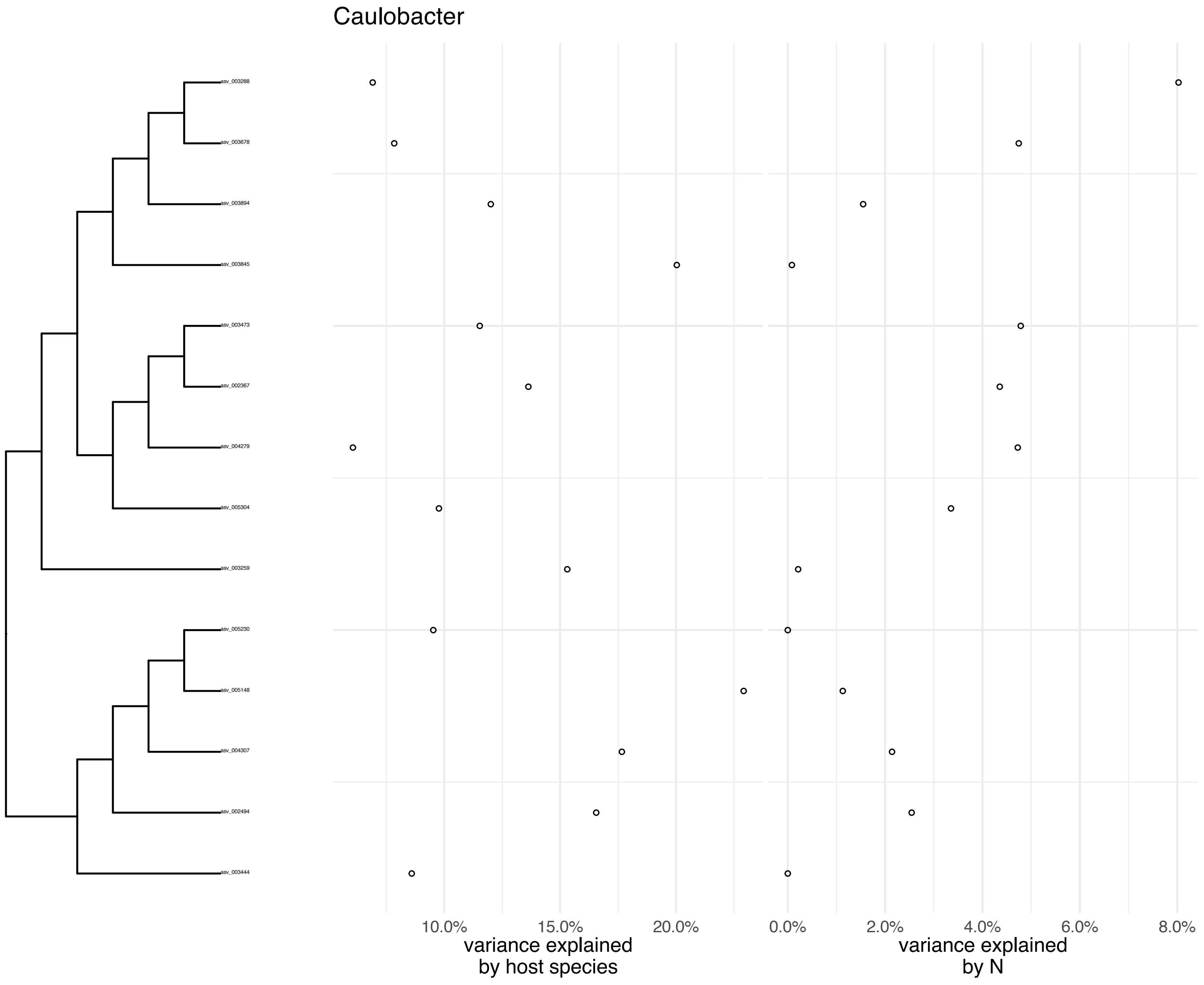
Caulobacter
asv_003288
asv_003678
asv_003894
asv_003845
asv_003473
asv_002367
asv_004279
asv_005304
asv_003259
asv_005230
asv_005148
asv_004307
asv_002494
asv_003444
10.0%
15.0%
20.0%
variance explained by host species
0.0%
2.0%
4.0%
6.0%
8.0%
variance explained by N

Chungangia

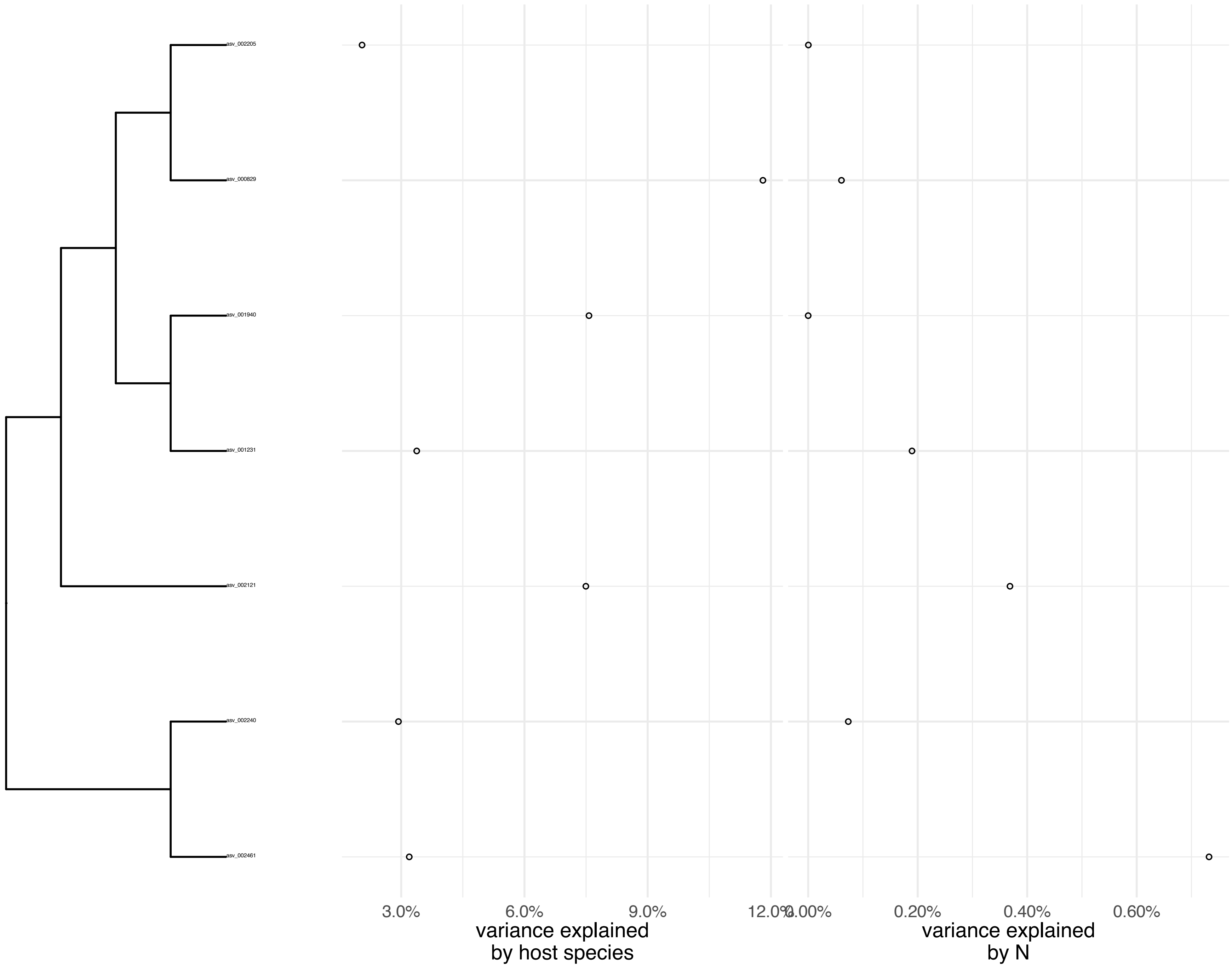
asv_002205
asv_000829
asv_001940
asv_001231
asv_002121
asv_002240
asv_002461
3.0%
6.0%
9.0%
12.0%
variance explained
by host species
0.00%
0.20%
0.40%
0.60%
variance explained
by N

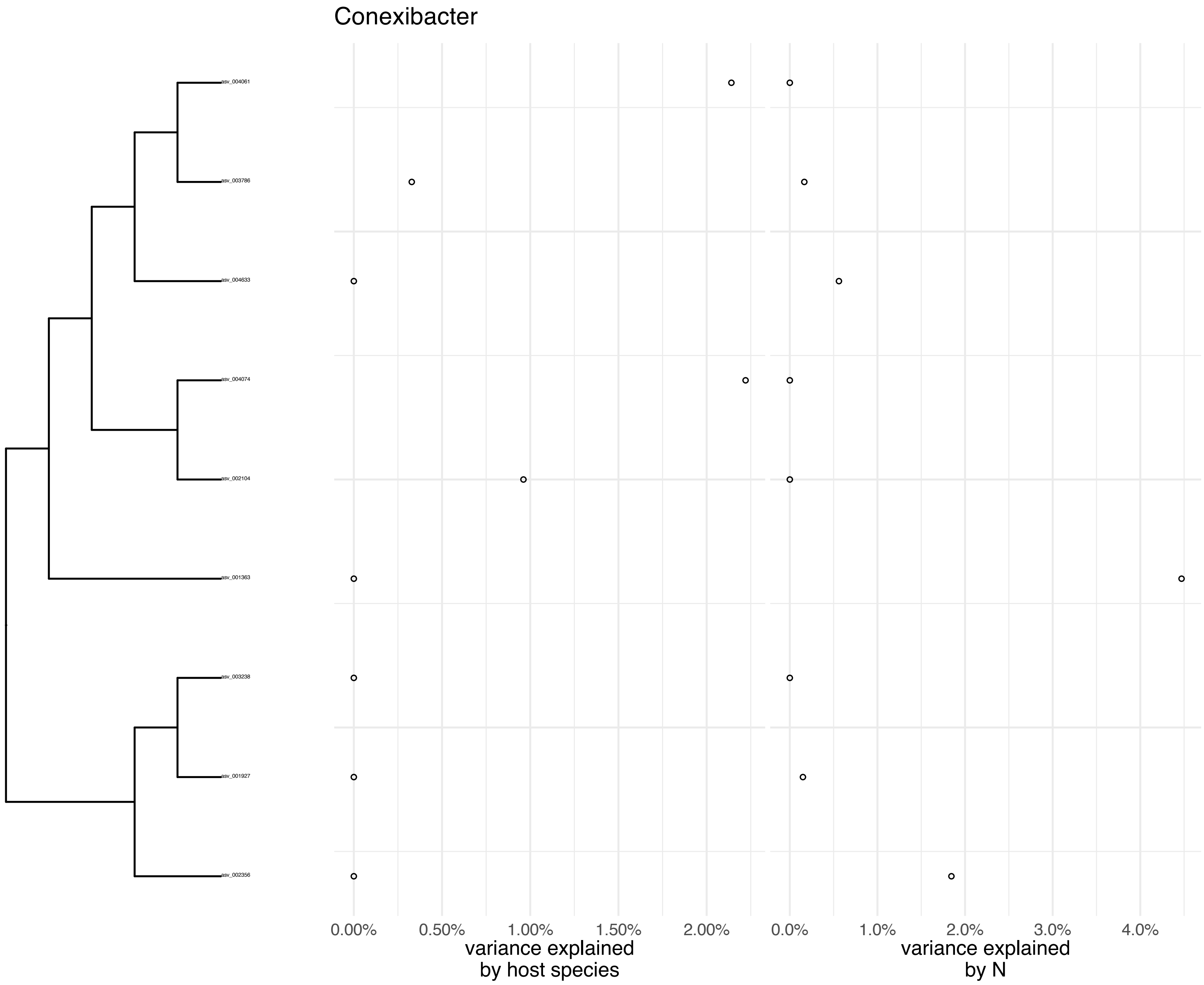
Conexibacter
asv_004061
asv_003786
asv_004633
asv_004074
asv_002104
asv_001363
asv_003238
asv_001927
asv_002356
0.00%
0.50%
1.00%
1.50%
2.00%
variance explained
by host species
0.0%
1.0%
2.0%
3.0%
4.0%
variance explained
by N

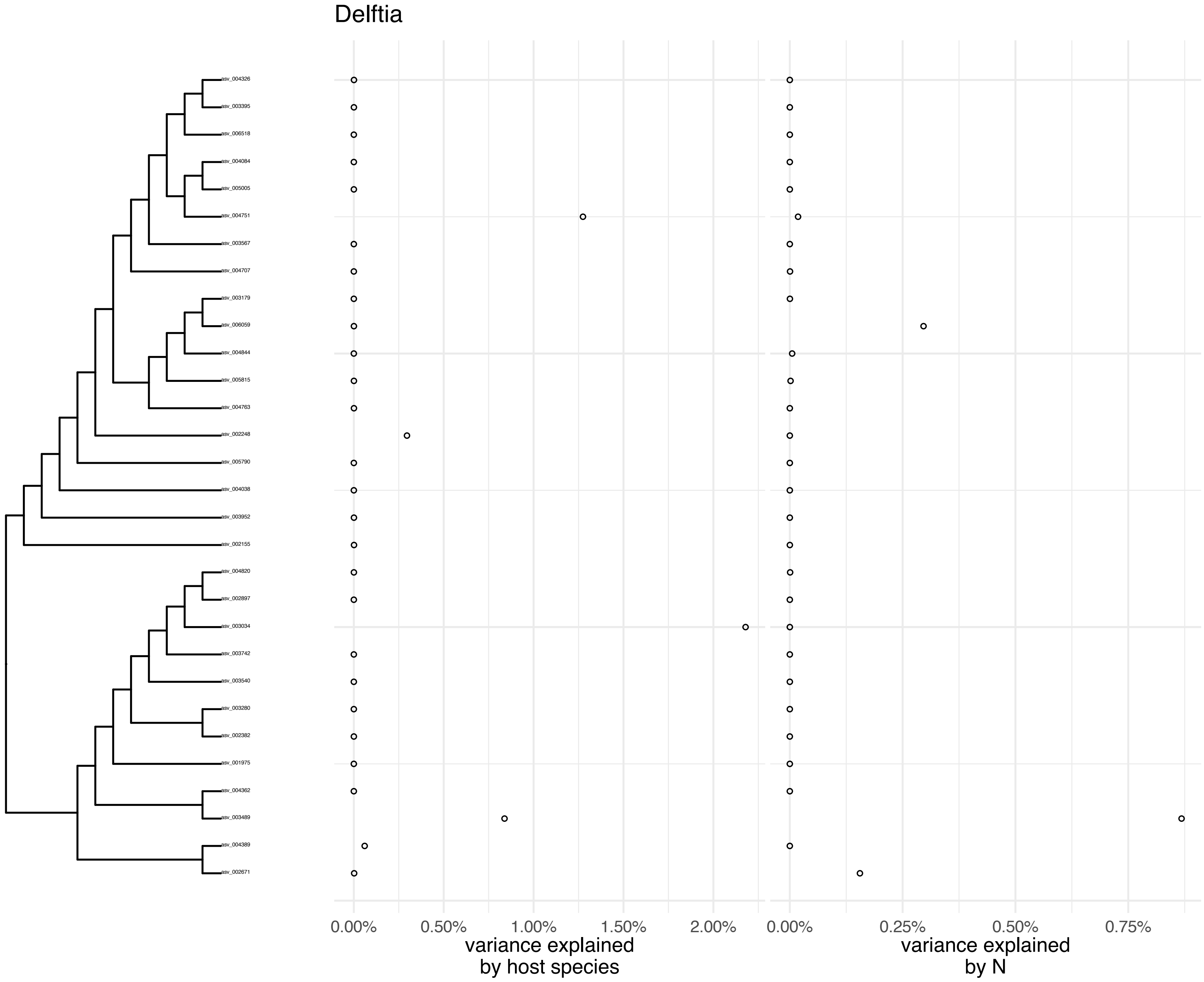

Delftia
asv_004326
asv_003395
asv_006518
asv_004084
asv_005005
asv_004751
asv_003567
asv_004707
asv_003179
asv_006059
asv_004844
asv_005815
asv_004763
asv_002248
asv_005790
asv_004038
asv_003952
asv_002155
asv_004820
asv_002897
asv_003034
asv_003742
asv_003540
asv_003280
asv_002382
asv_001975
asv_004362
asv_003489
asv_004389
asv_002671
0.00%
0.50%
1.00%
1.50%
2.00%
variance explained
by host species
0.00%
0.25%
0.50%
0.75%
variance explained
by N

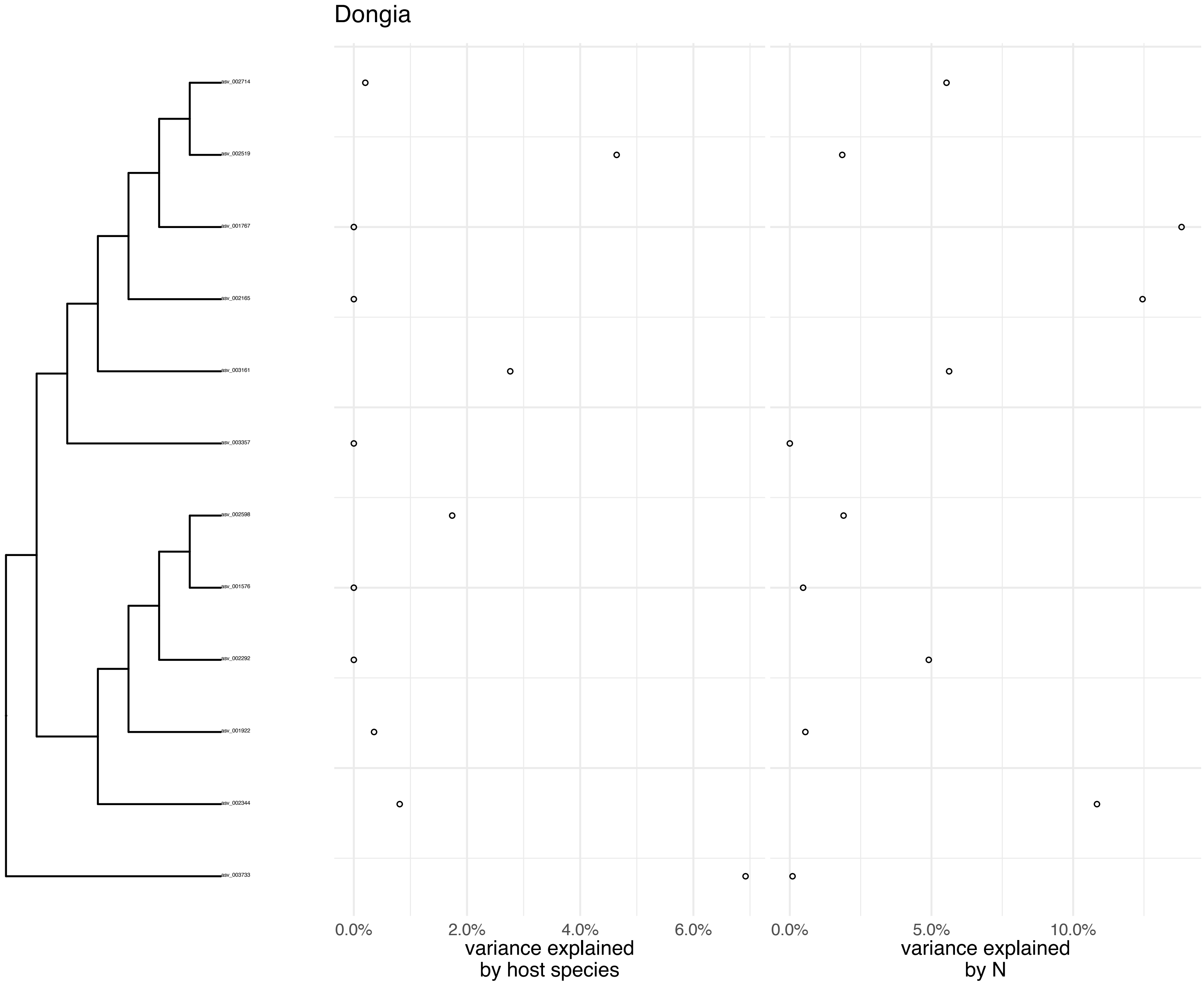
Dongia
asv_002714
asv_002519
asv_001767
asv_002165
asv_003161
asv_003357
asv_002598
asv_001576
asv_002292
asv_001922
asv_002344
asv_003733
0.0%
2.0%
4.0%
6.0%
variance explained
by host species
0.0%
5.0%
10.0%
variance explained
by N

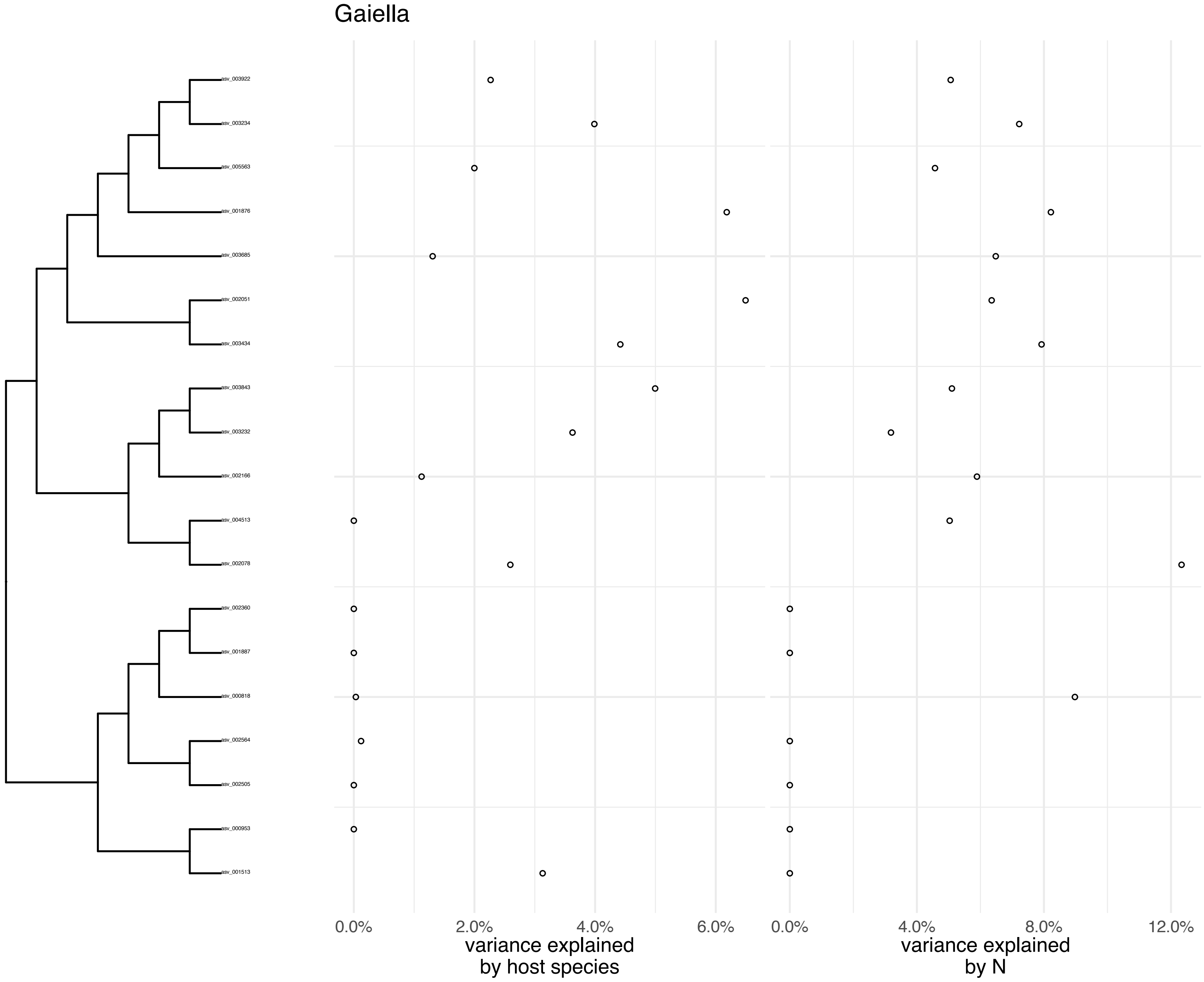
Gaiella
asv_003922
asv_003234
asv_005563
asv_001876
asv_003685
asv_002051
asv_003434
asv_003843
asv_003232
asv_002166
asv_004513
asv_002078
asv_002360
asv_001887
asv_000818
asv_002564
asv_002505
asv_000953
asv_001513
0.0%
2.0%
4.0%
6.0%
variance explained
by host species
0.0%
4.0%
8.0%
12.0%
variance explained
by N

Gemmatimonas

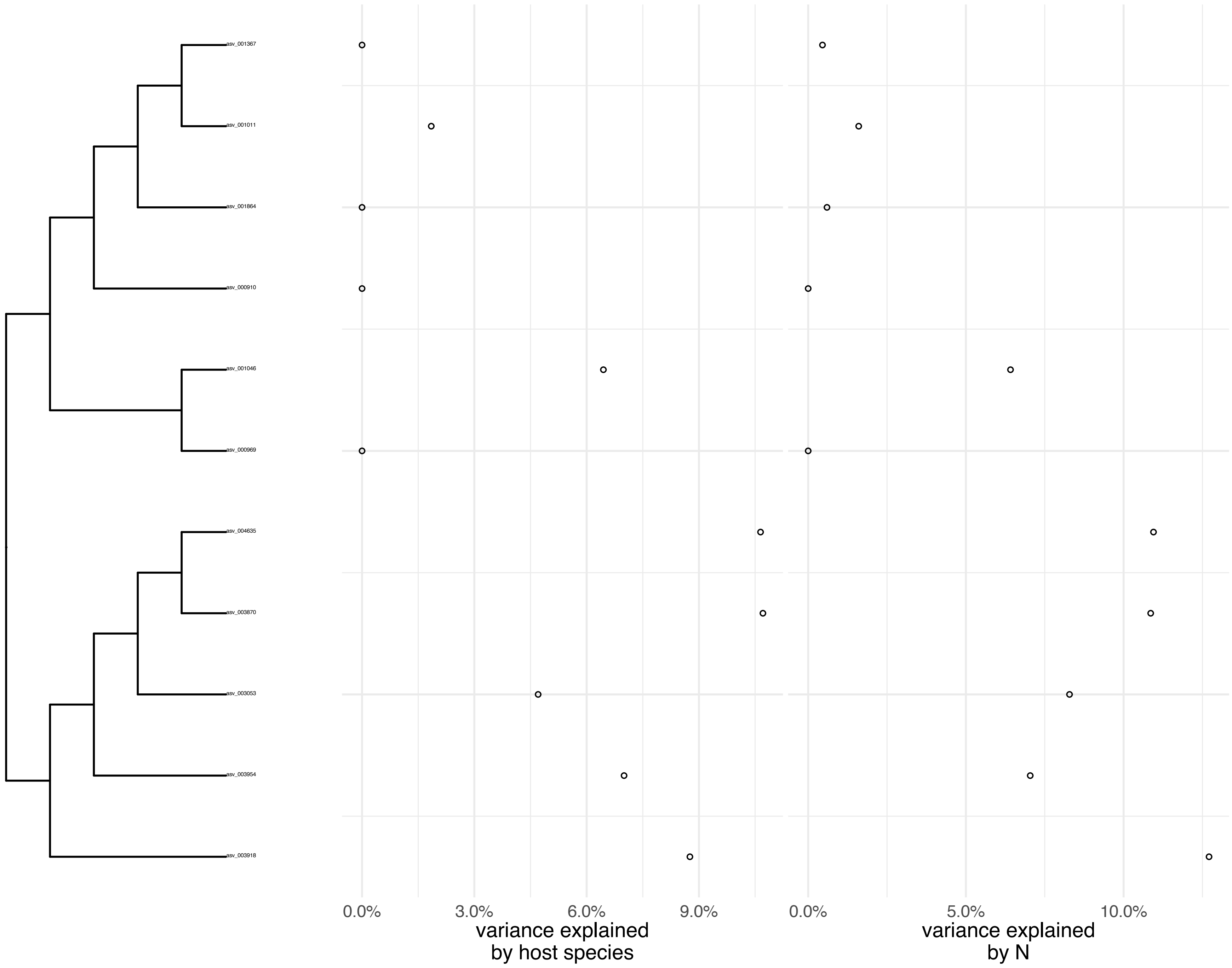

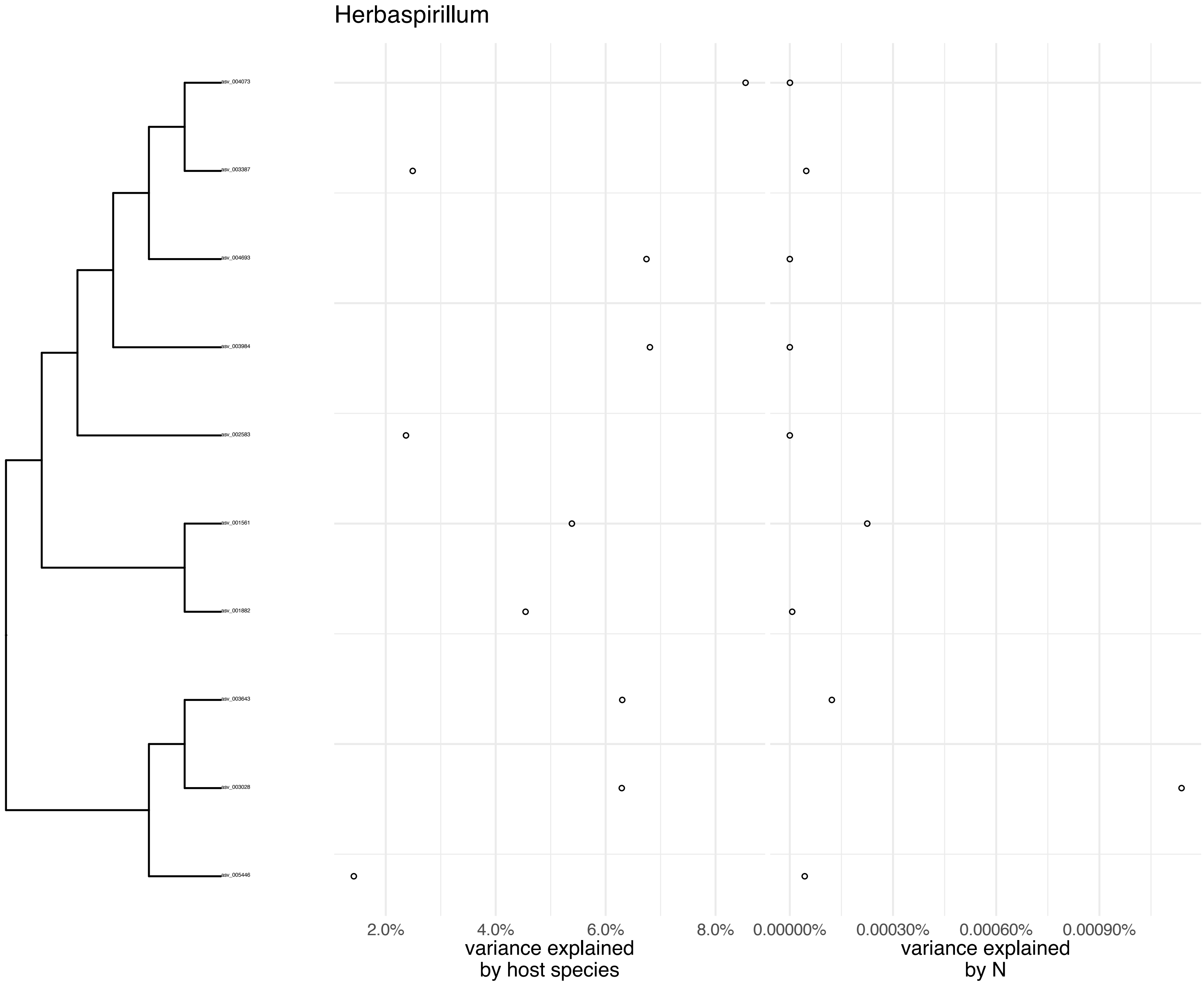
Herbaspirillum
asv_004073
asv_003387
asv_004693
asv_003984
asv_002583
asv_001561
asv_001882
asv_003643
asv_003028
asv_005446
2.0%
4.0%
6.0%
8.0%
variance explained by host species
0.00000%
0.00030%
0.00060%
0.00090%
variance explained by N

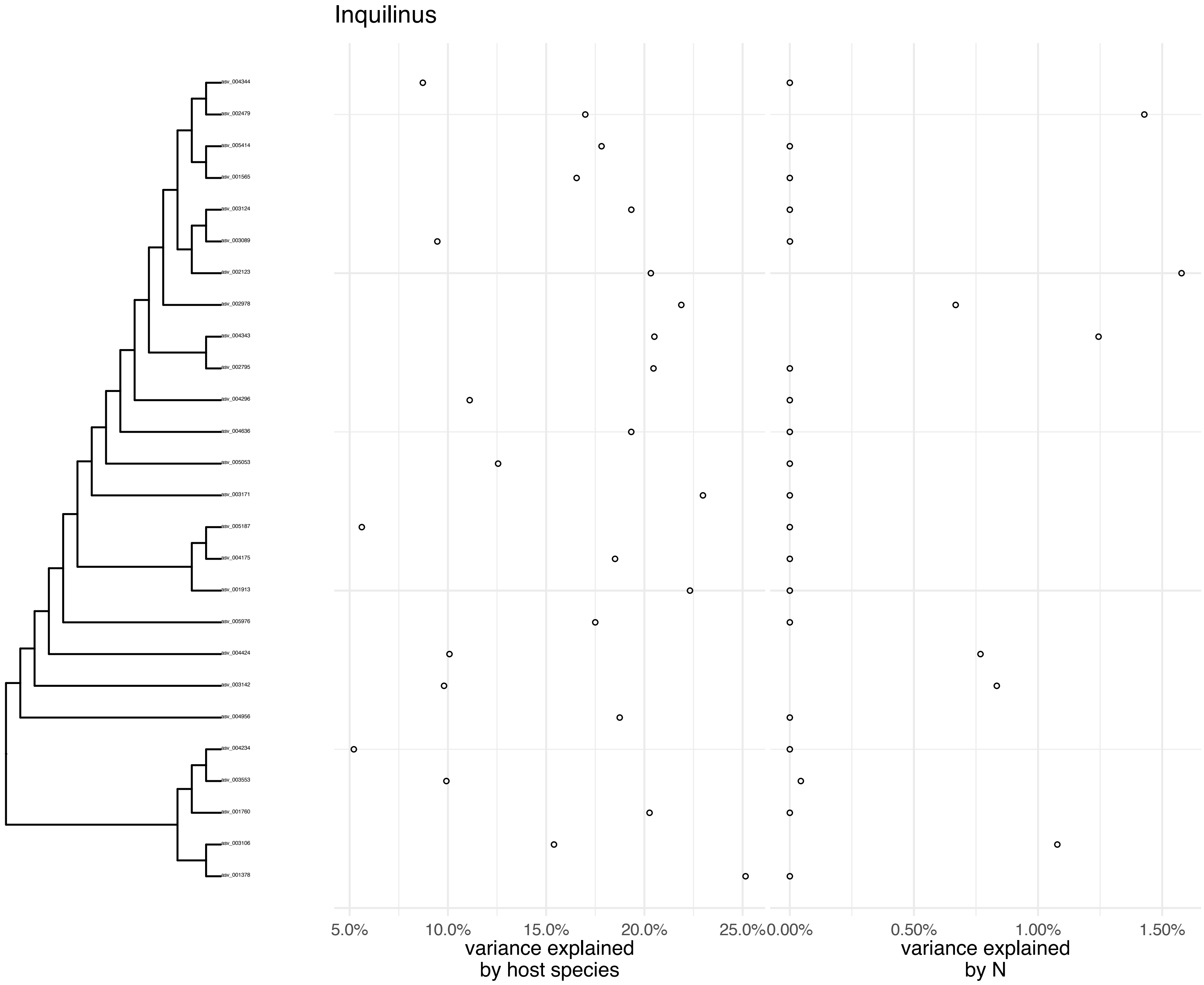
Inquilinus
asv_004344
asv_002479
asv_005414
asv_001565
asv_003124
asv_003089
asv_002123
asv_002978
asv_004343
asv_002795
asv_004296
asv_004636
asv_005053
asv_003171
asv_005187
asv_004175
asv_001913
asv_005976
asv_004424
asv_003142
asv_004956
asv_004234
asv_003553
asv_001760
asv_003106
asv_001378
5.0%
10.0%
15.0%
20.0%
25.0%
variance explained by host species
0.00%
0.50%
1.00%
1.50%
variance explained by N

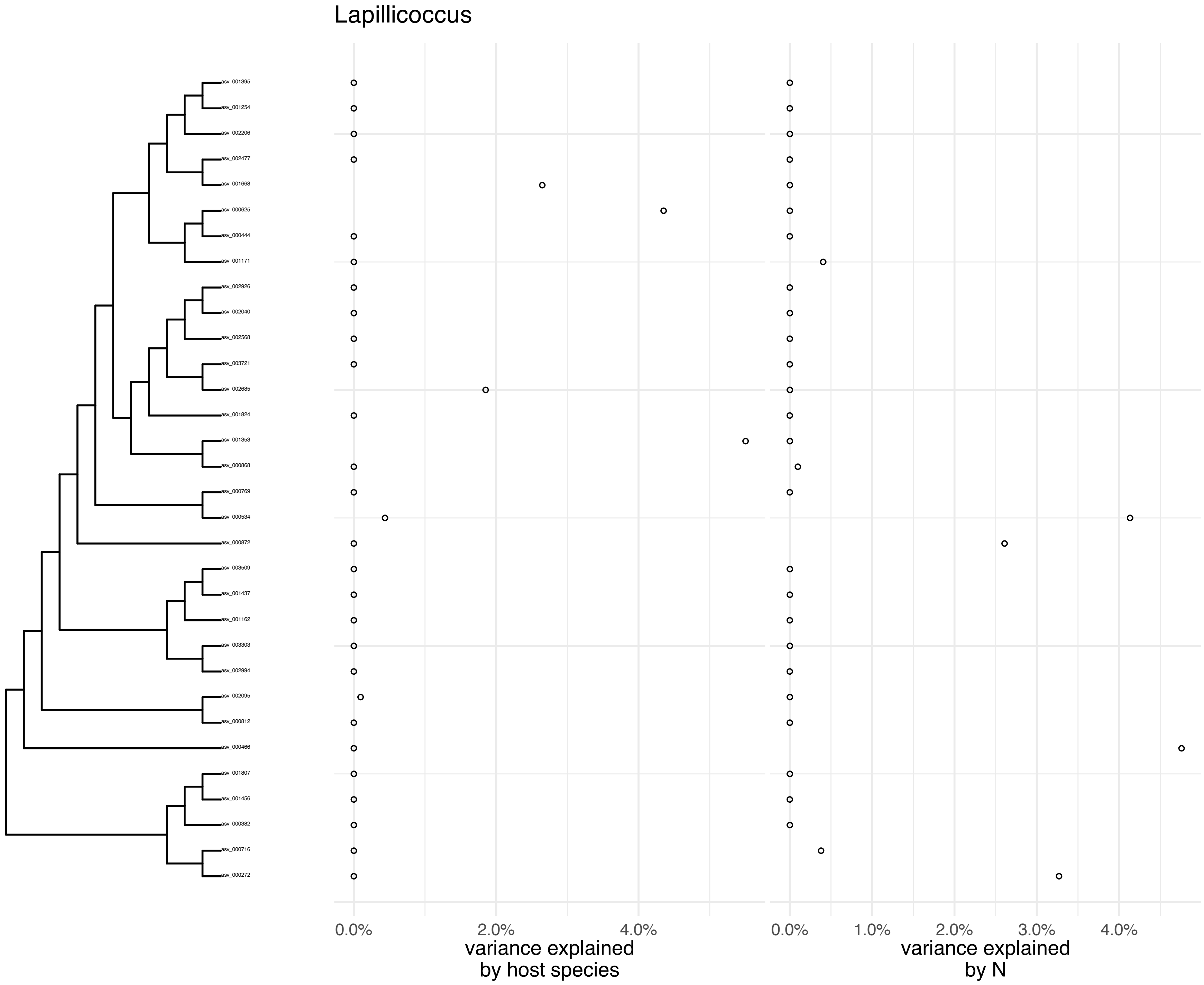
Lapillicoccus
asv_001395
asv_001254
asv_002206
asv_002477
asv_001668
asv_000625
asv_000444
asv_001171
asv_002926
asv_002040
asv_002568
asv_003721
asv_002685
asv_001824
asv_001353
asv_000868
asv_000769
asv_000534
asv_000872
asv_003509
asv_001437
asv_001162
asv_003303
asv_002994
asv_002095
asv_000812
asv_000466
asv_001807
asv_001456
asv_000382
asv_000716
asv_000272
0.0%
2.0%
4.0%
variance explained
by host species
0.0%
1.0%
2.0%
3.0%
4.0%
variance explained
by N

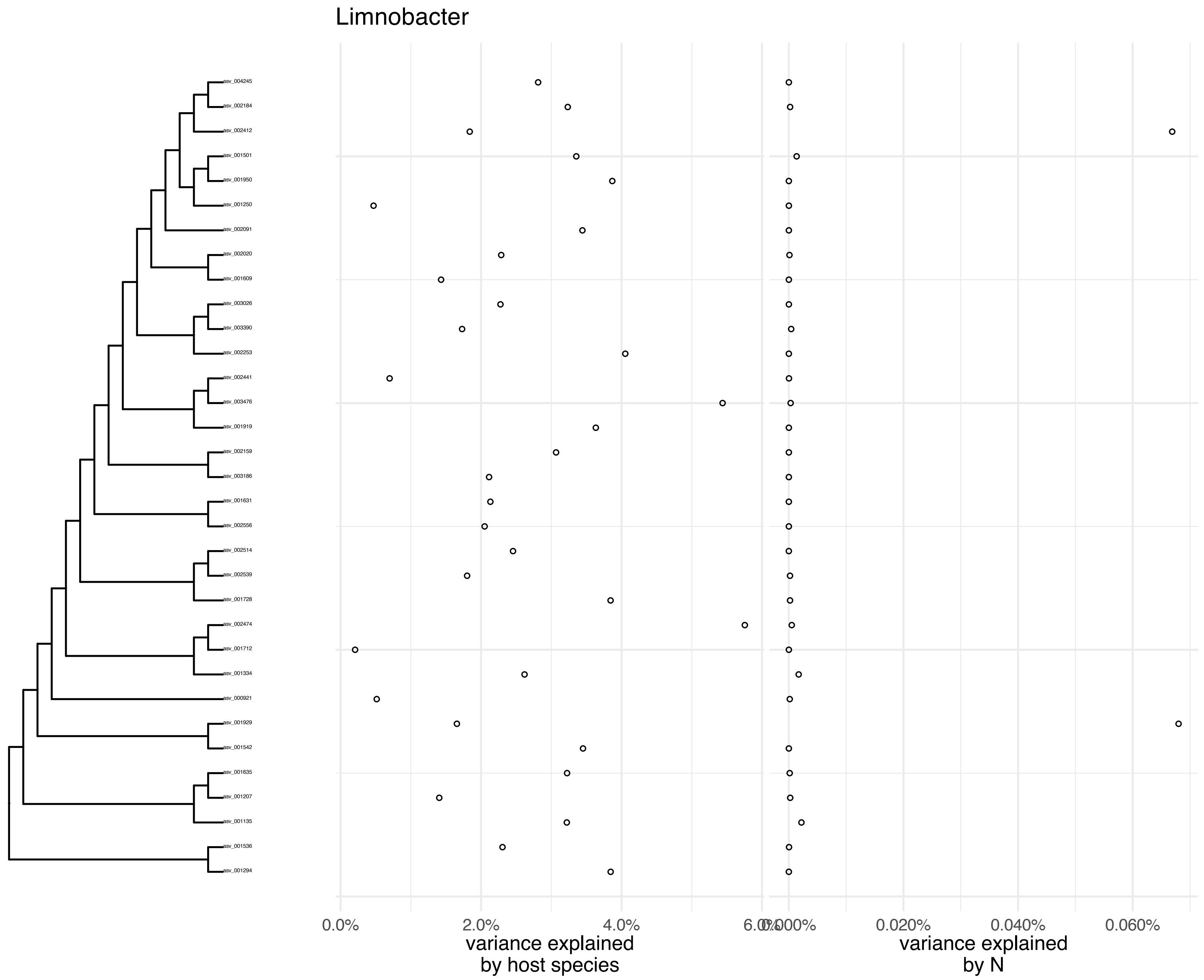

Limnobacter
asv_004245
asv_002184
asv_002412
asv_001501
asv_001950
asv_001250
asv_002091
asv_002020
asv_001609
asv_003026
asv_003390
asv_002253
asv_002441
asv_003476
asv_001919
asv_002159
asv_003186
asv_001631
asv_002556
asv_002514
asv_002539
asv_001728
asv_002474
asv_001712
asv_001334
asv_000921
asv_001929
asv_001542
asv_001635
asv_001207
asv_001135
asv_001536
asv_001294
0.0%
2.0%
4.0%
6.0%
variance explained
by host species
0.000%
0.020%
0.040%
0.060%
variance explained
by N

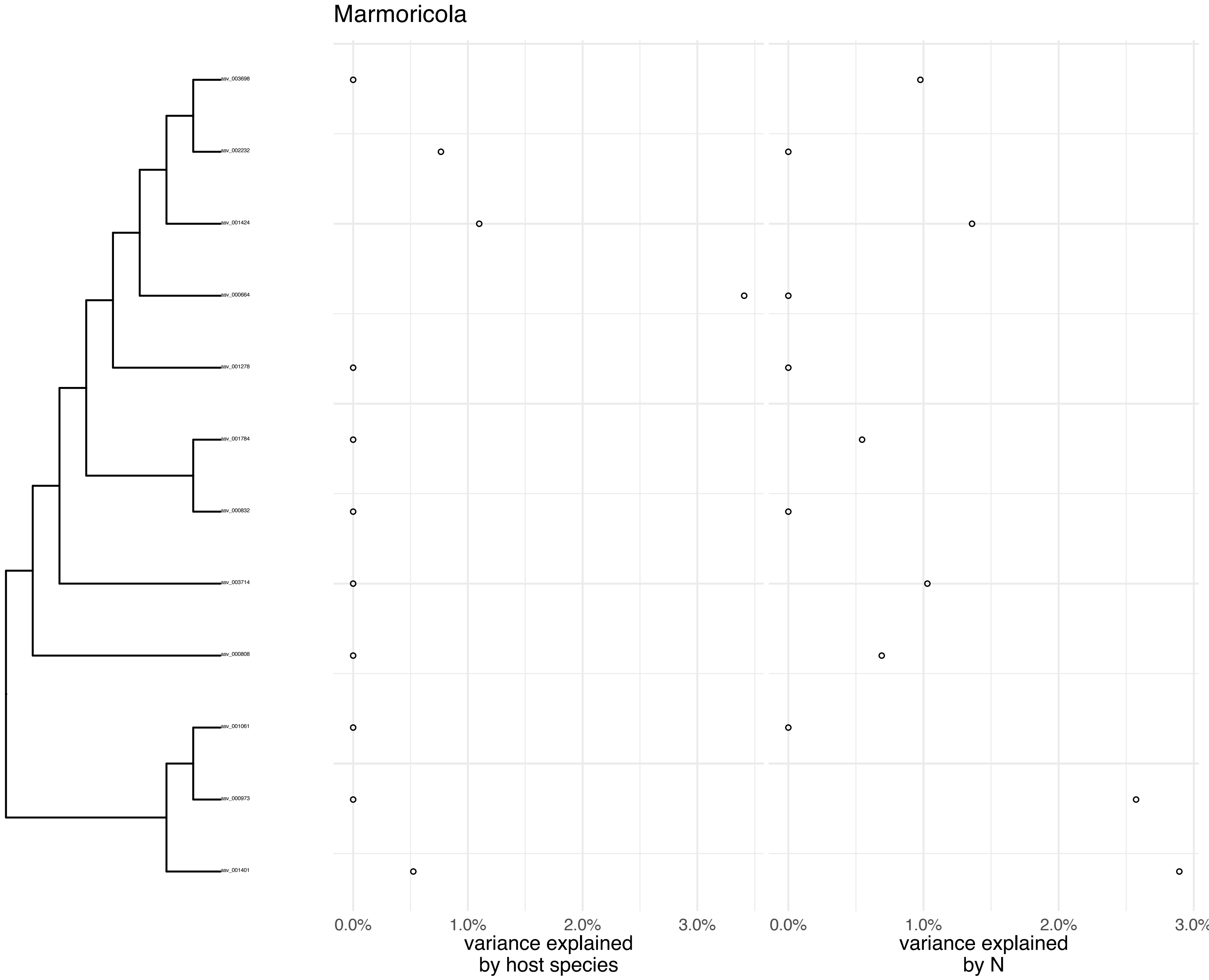
Marmoricola
asv_003698
asv_002232
asv_001424
asv_000664
asv_001278
asv_001784
asv_000832
asv_003714
asv_000808
asv_001061
asv_000973
asv_001401
0.0%
1.0%
2.0%
3.0%
variance explained
by host species
0.0%
1.0%
2.0%
3.0%
variance explained
by N

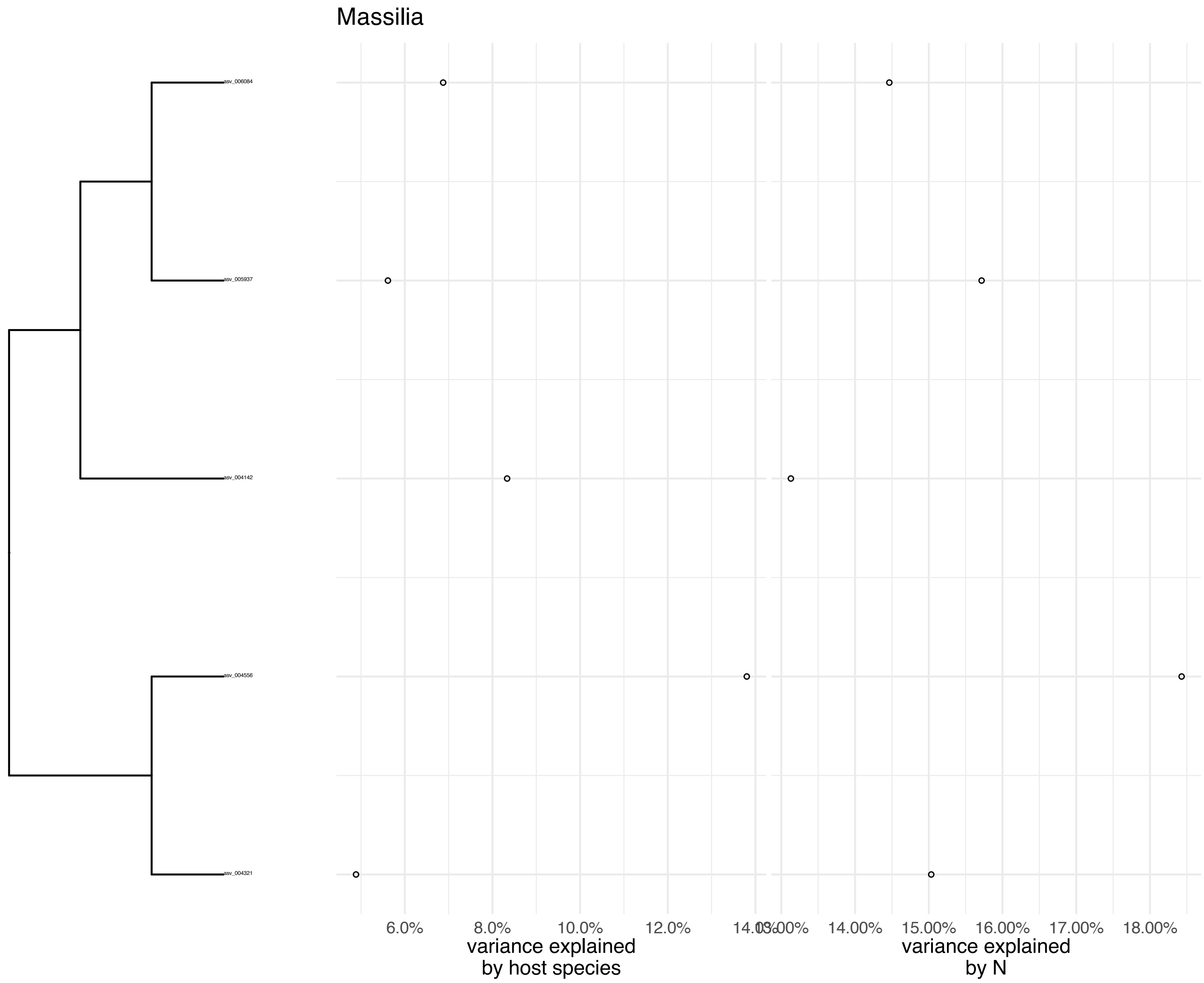

Massilia
asv_006084
asv_005937
asv_004142
asv_004556
asv_004321
6.0%
8.0%
10.0%
12.0%
14.0%
13.00%
14.00%
15.00%
16.00%
17.00%
18.00%
variance explained by host species
variance explained by N

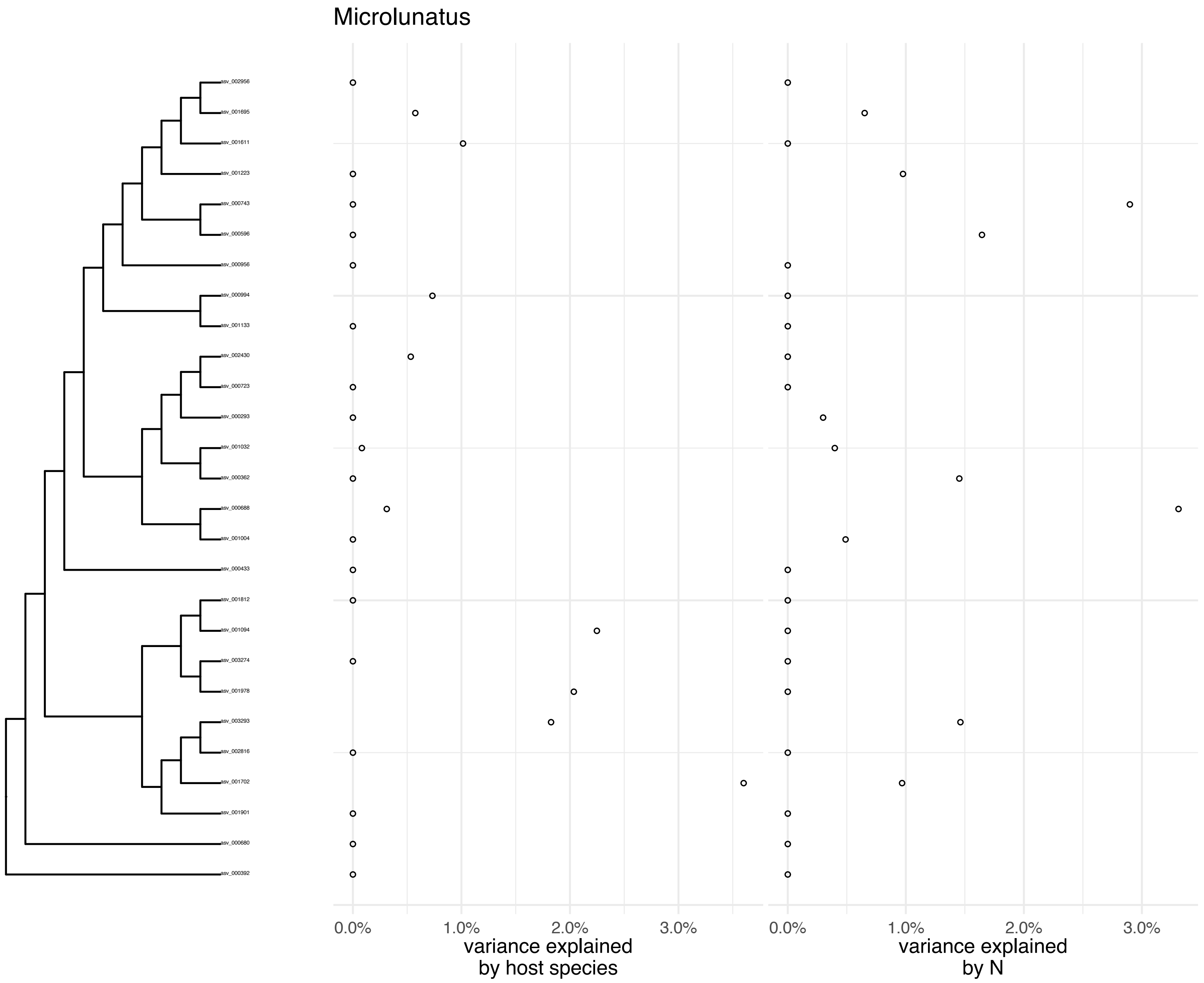
Microlunatus
asv_002956
asv_001695
asv_001611
asv_001223
asv_000743
asv_000596
asv_000956
asv_000994
asv_001133
asv_002430
asv_000723
asv_000293
asv_001032
asv_000362
asv_000688
asv_001004
asv_000433
asv_001812
asv_001094
asv_003274
asv_001978
asv_003293
asv_002816
asv_001702
asv_001901
asv_000680
asv_000392
0.0%
1.0%
2.0%
3.0%
variance explained
by host species
0.0%
1.0%
2.0%
3.0%
variance explained
by N

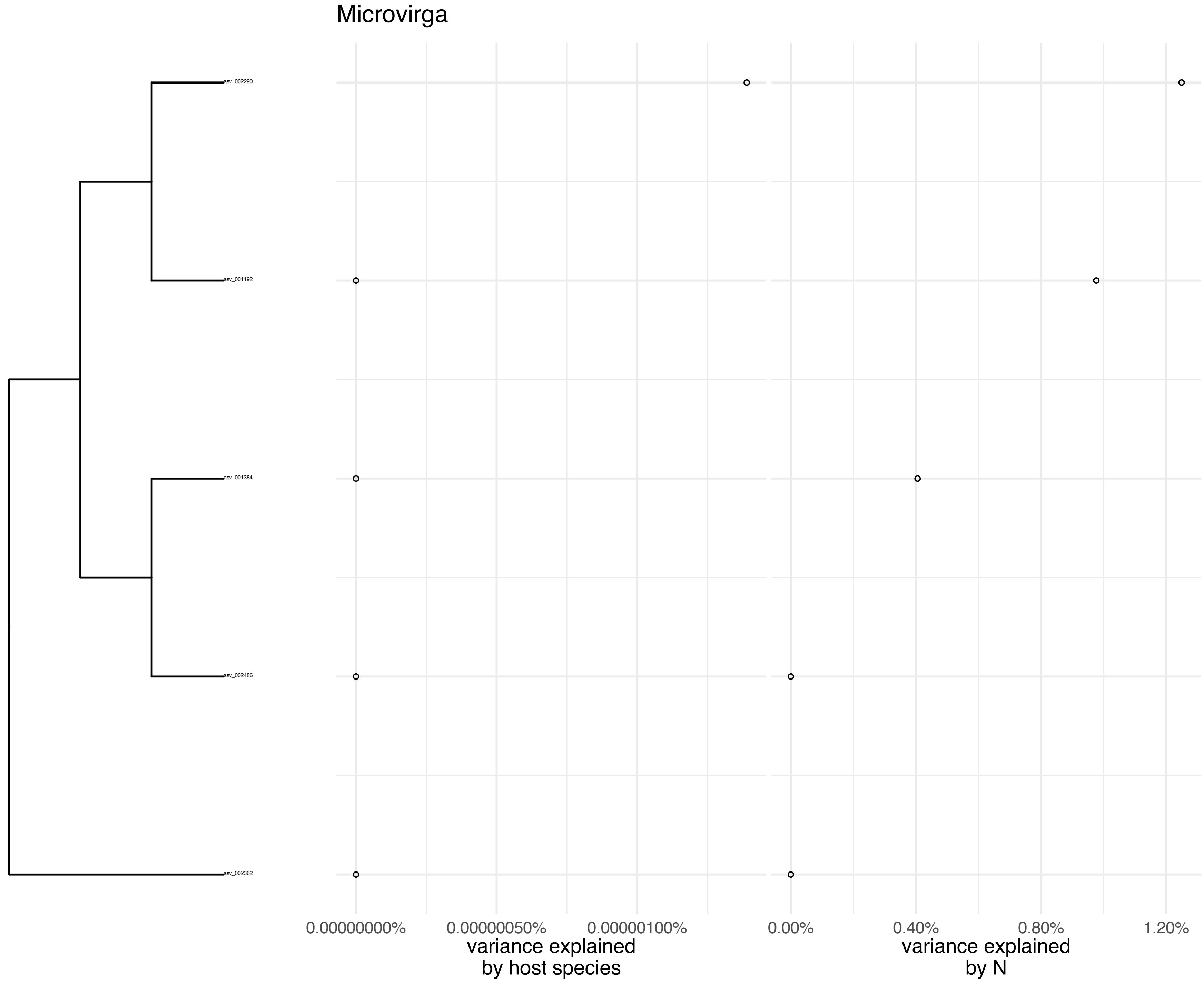
Microvirga
asv_002290
asv_001192
asv_001384
asv_002486
asv_002362
0.00000000%
0.00000050%
0.00000100%
variance explained
by host species
0.00%
0.40%
0.80%
1.20%
variance explained
by N

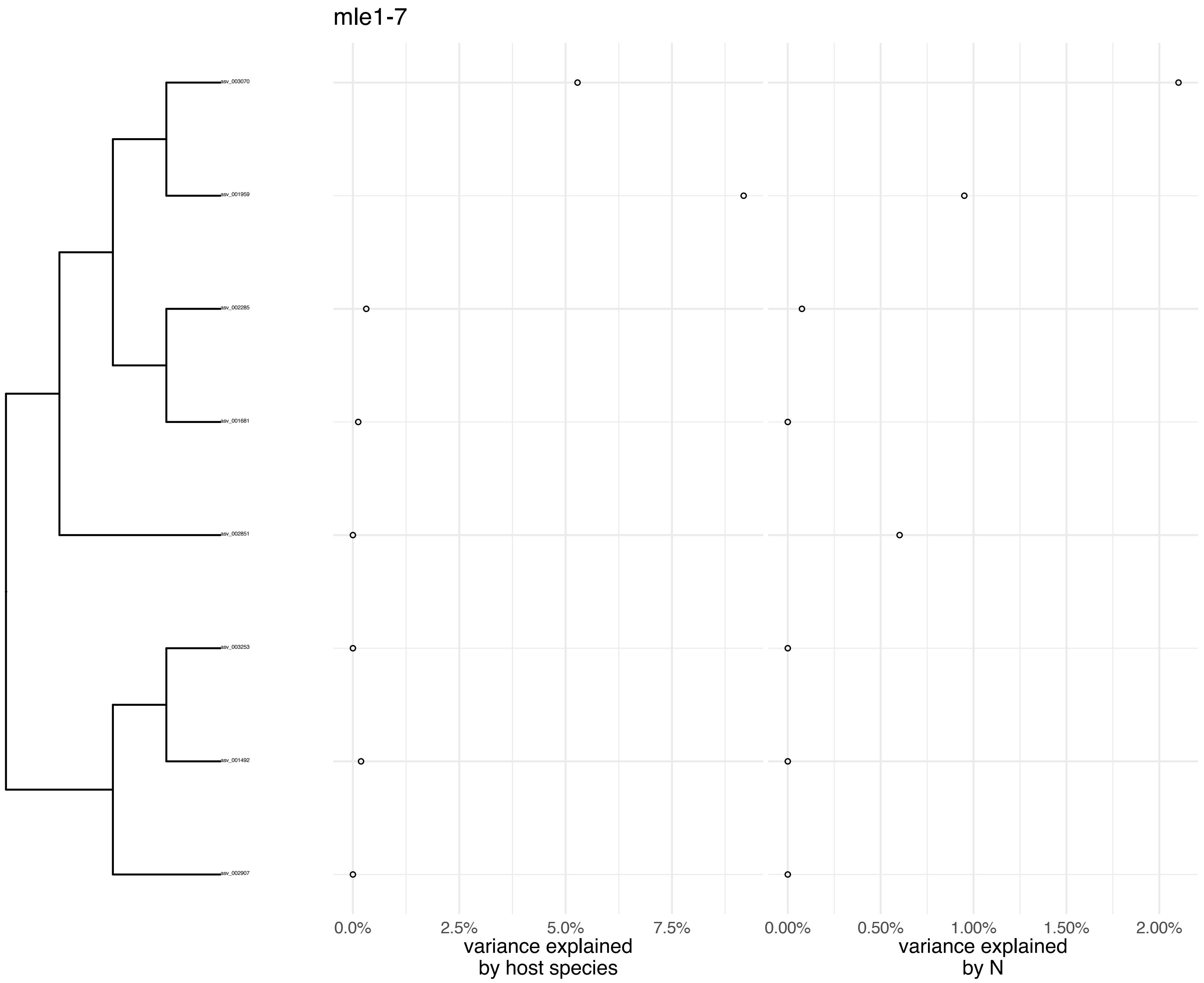
mle1-7
asv_003070
asv_001959
asv_002285
asv_001681
asv_002851
asv_003253
asv_001492
asv_002907
0.0%
2.5%
5.0%
7.5%
variance explained
by host species
0.00%
0.50%
1.00%
1.50%
2.00%
variance explained
by N

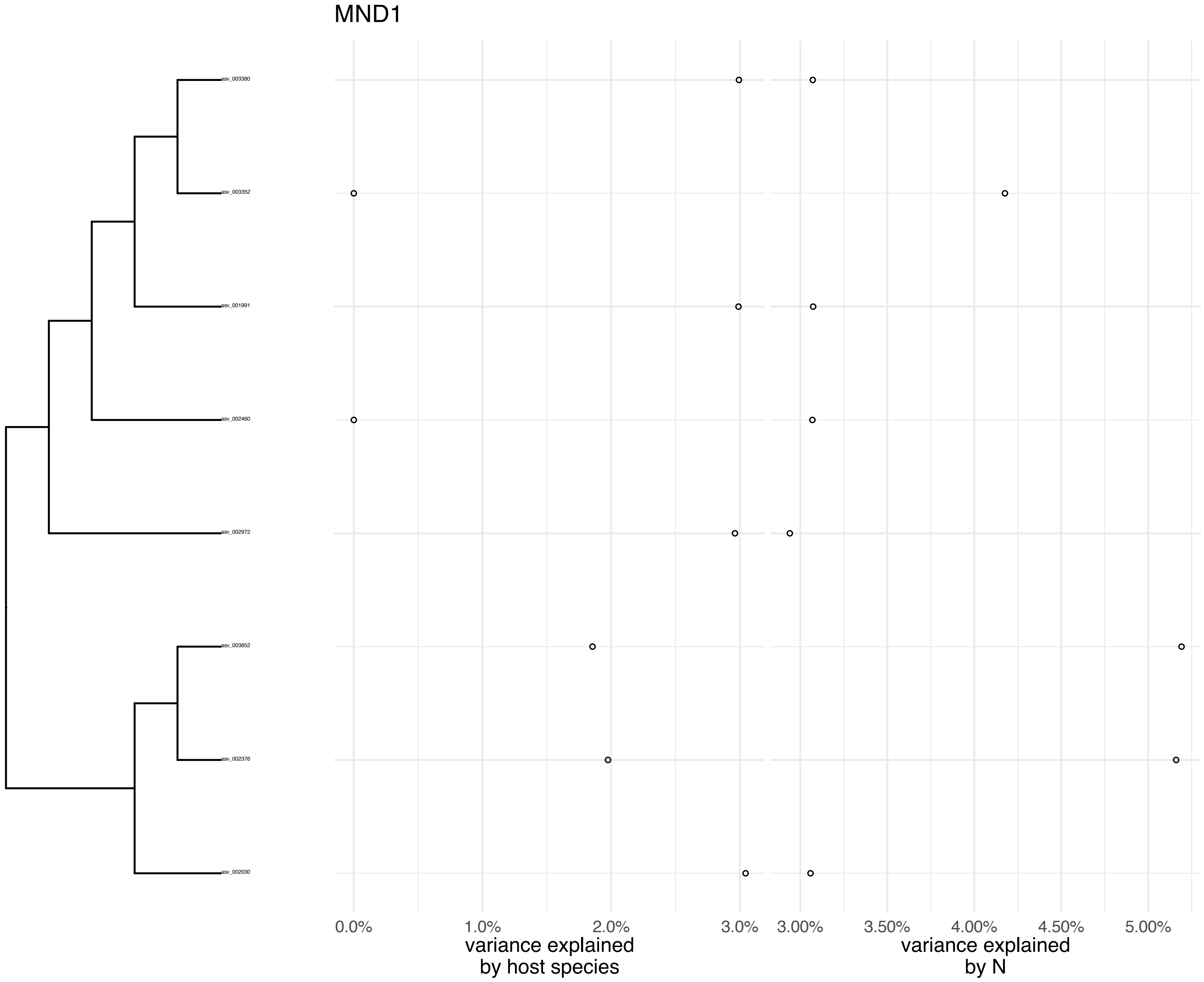
MND1
asv_003380
asv_003352
asv_001991
asv_002460
asv_002972
asv_003852
asv_002376
asv_002030
0.0%
1.0%
2.0%
3.0%
variance explained
by host species
3.00%
3.50%
4.00%
4.50%
5.00%
variance explained
by N

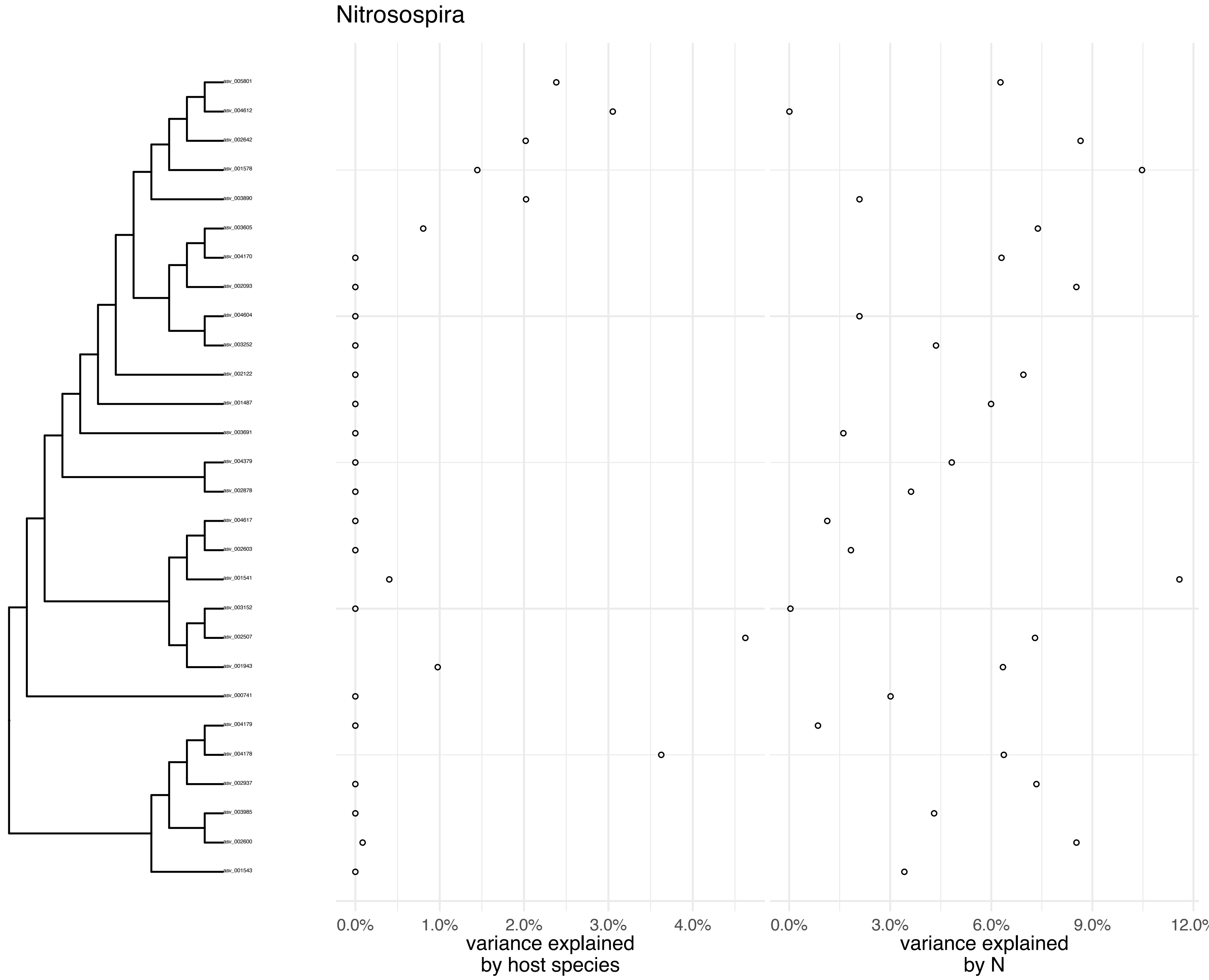

Nitrosospira
asv_005801
asv_004612
asv_002642
asv_001578
asv_003890
asv_003605
asv_004170
asv_002093
asv_004604
asv_003252
asv_002122
asv_001487
asv_003691
asv_004379
asv_002878
asv_004617
asv_002603
asv_001541
asv_003152
asv_002507
asv_001943
asv_000741
asv_004179
asv_004178
asv_002937
asv_003985
asv_002600
asv_001543
0.0%
1.0%
2.0%
3.0%
4.0%
variance explained by host species
0.0%
3.0%
6.0%
9.0%
12.0%
variance explained by N

Nitrospira

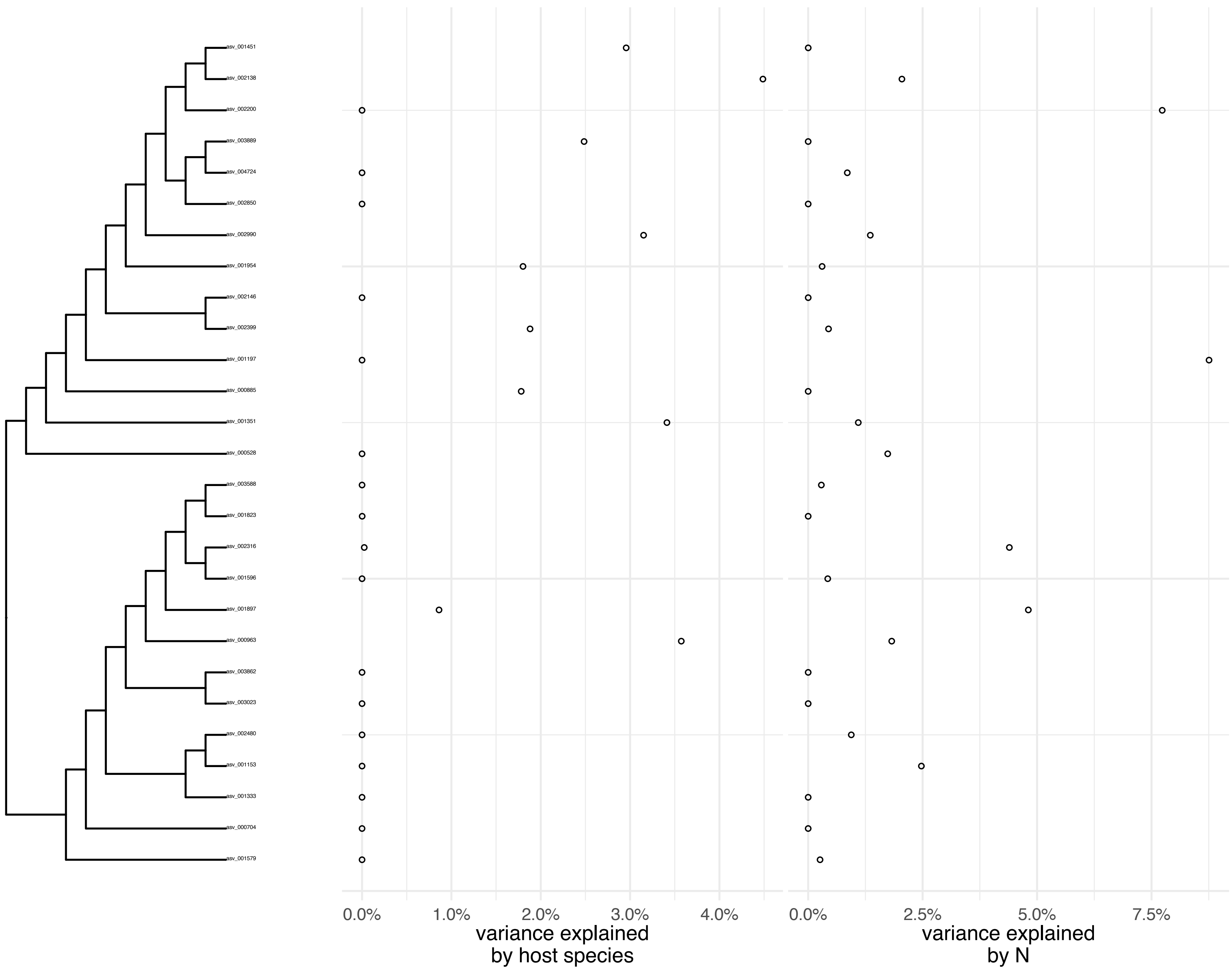
asv_001451
asv_002138
asv_002200
asv_003889
asv_004724
asv_002850
asv_002990
asv_001954
asv_002146
asv_002399
asv_001197
asv_000885
asv_001351
asv_000528
asv_003588
asv_001823
asv_002316
asv_001596
asv_001897
asv_000963
asv_003862
asv_003023
asv_002480
asv_001153
asv_001333
asv_000704
asv_001579
0.0%
1.0%
2.0%
3.0%
4.0%
variance explained
by host species
0.0%
2.5%
5.0%
7.5%
variance explained
by N

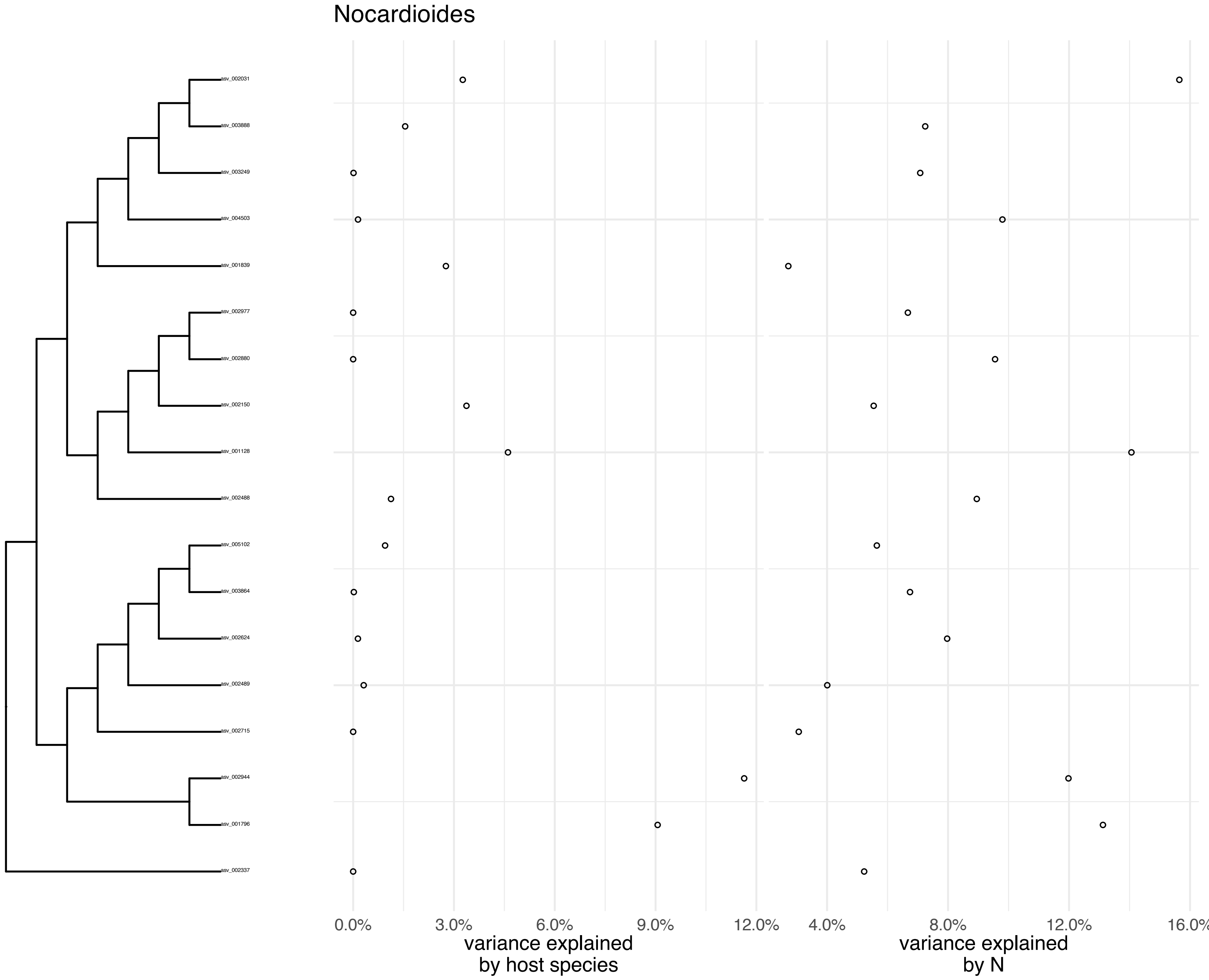
Nocardioides
asv_002031
asv_003888
asv_003249
asv_004503
asv_001839
asv_002977
asv_002880
asv_002150
asv_001128
asv_002488
asv_005102
asv_003864
asv_002624
asv_002489
asv_002715
asv_002944
asv_001796
asv_002337
0.0%
3.0%
6.0%
9.0%
12.0%
variance explained
by host species
4.0%
8.0%
12.0%
16.0%
variance explained
by N

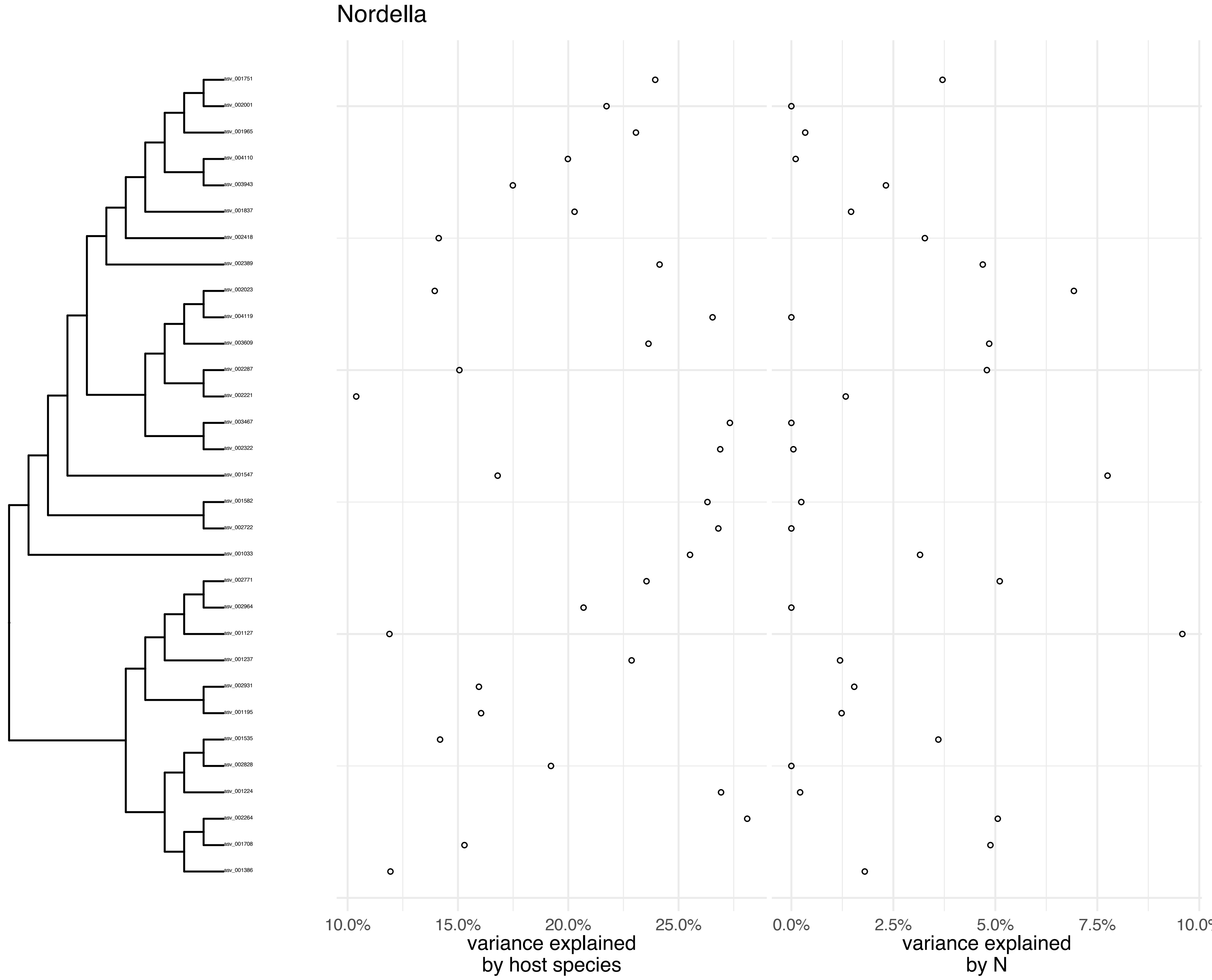
Nordella
asv_001751
asv_002001
asv_001965
asv_004110
asv_003943
asv_001837
asv_002418
asv_002389
asv_002023
asv_004119
asv_003609
asv_002287
asv_002221
asv_003467
asv_002322
asv_001547
asv_001582
asv_002722
asv_001033
asv_002771
asv_002964
asv_001127
asv_001237
asv_002931
asv_001195
asv_001535
asv_002828
asv_001224
asv_002264
asv_001708
asv_001386
10.0%
15.0%
20.0%
25.0%
variance explained by host species
0.0%
2.5%
5.0%
7.5%
10.0%
variance explained by N

Novosphingobium

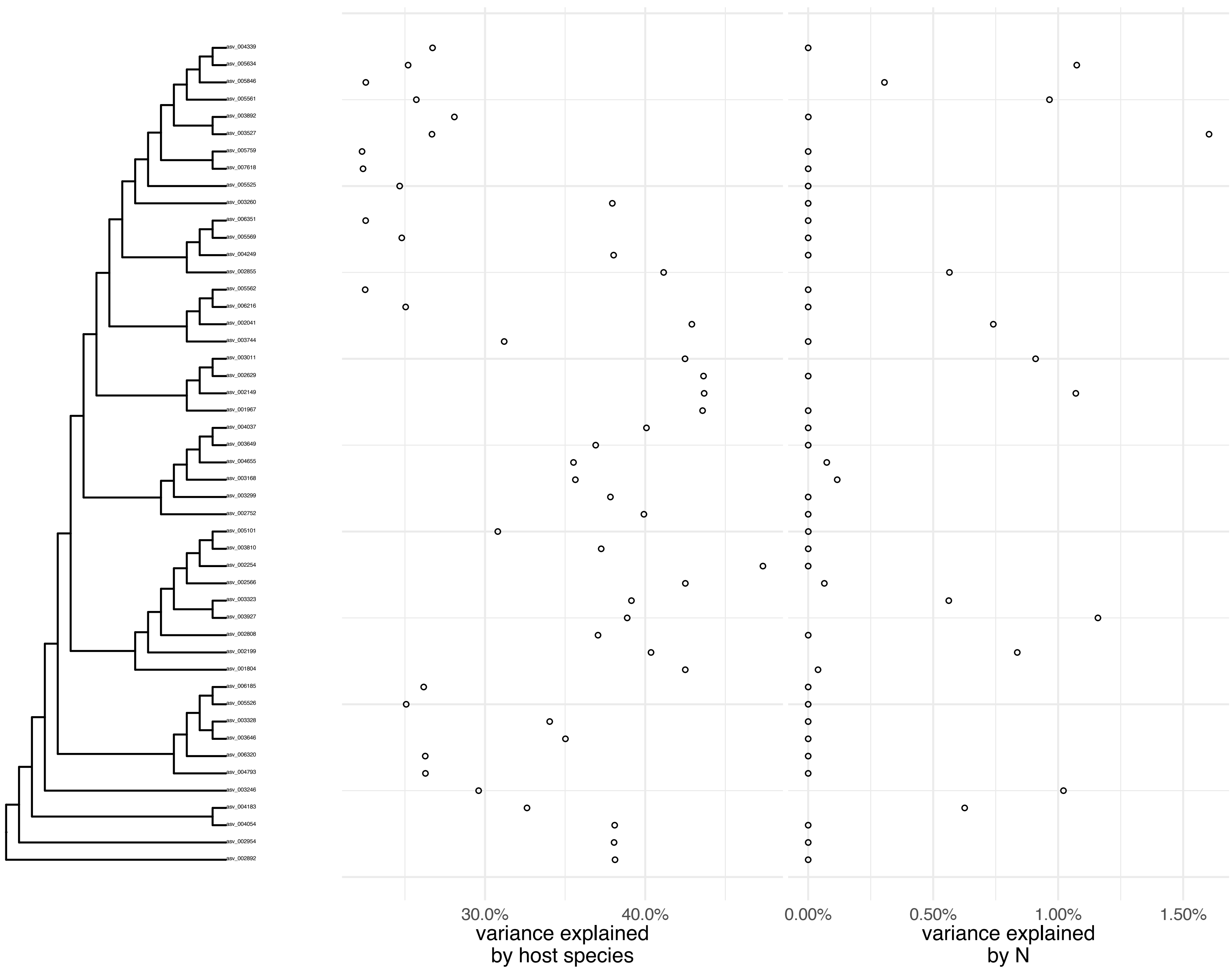

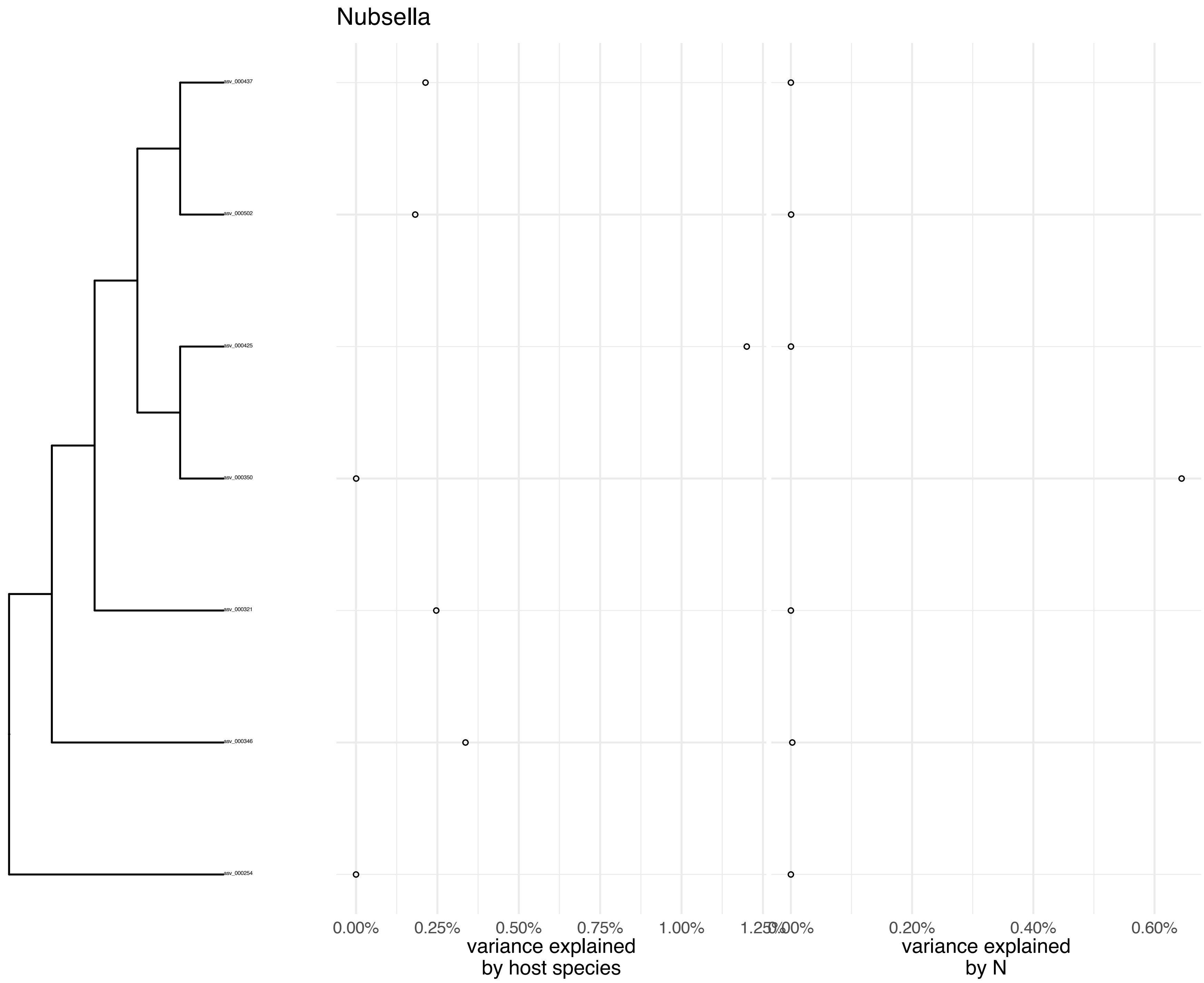
Nubsella
asv_000437
asv_000502
asv_000425
asv_000350
asv_000321
asv_000346
asv_000254
0.00%
0.25%
0.50%
0.75%
1.00%
1.25%
variance explained by host species
0.00%
0.20%
0.40%
0.60%
variance explained by N

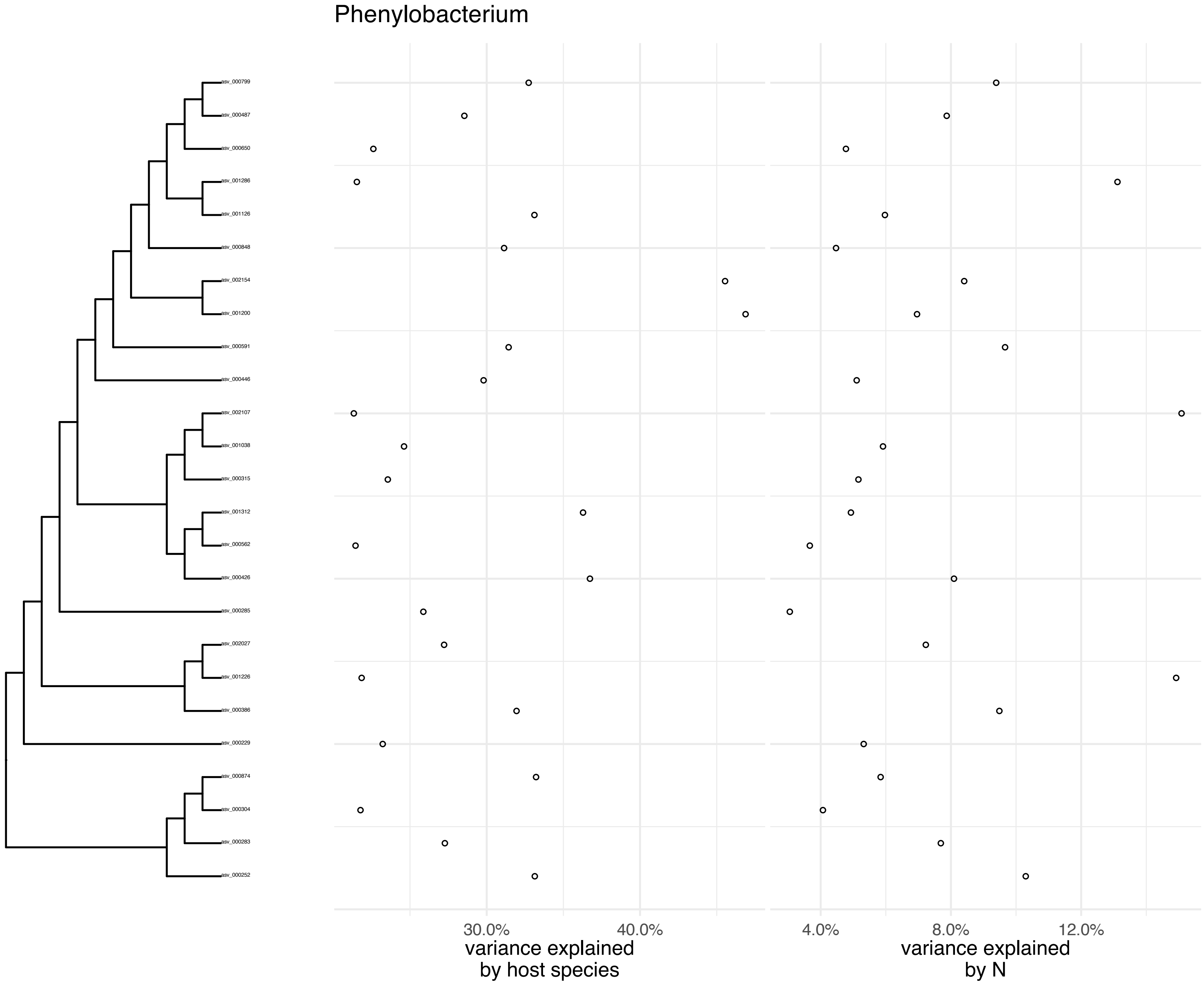
Phenylobacterium
asv_000799
asv_000487
asv_000650
asv_001286
asv_001126
asv_000848
asv_002154
asv_001200
asv_000591
asv_000446
asv_002107
asv_001038
asv_000315
asv_001312
asv_000562
asv_000426
asv_000285
asv_002027
asv_001226
asv_000386
asv_000229
asv_000874
asv_000304
asv_000283
asv_000252
30.0%
40.0%
variance explained
by host species
4.0%
8.0%
12.0%
variance explained
by N

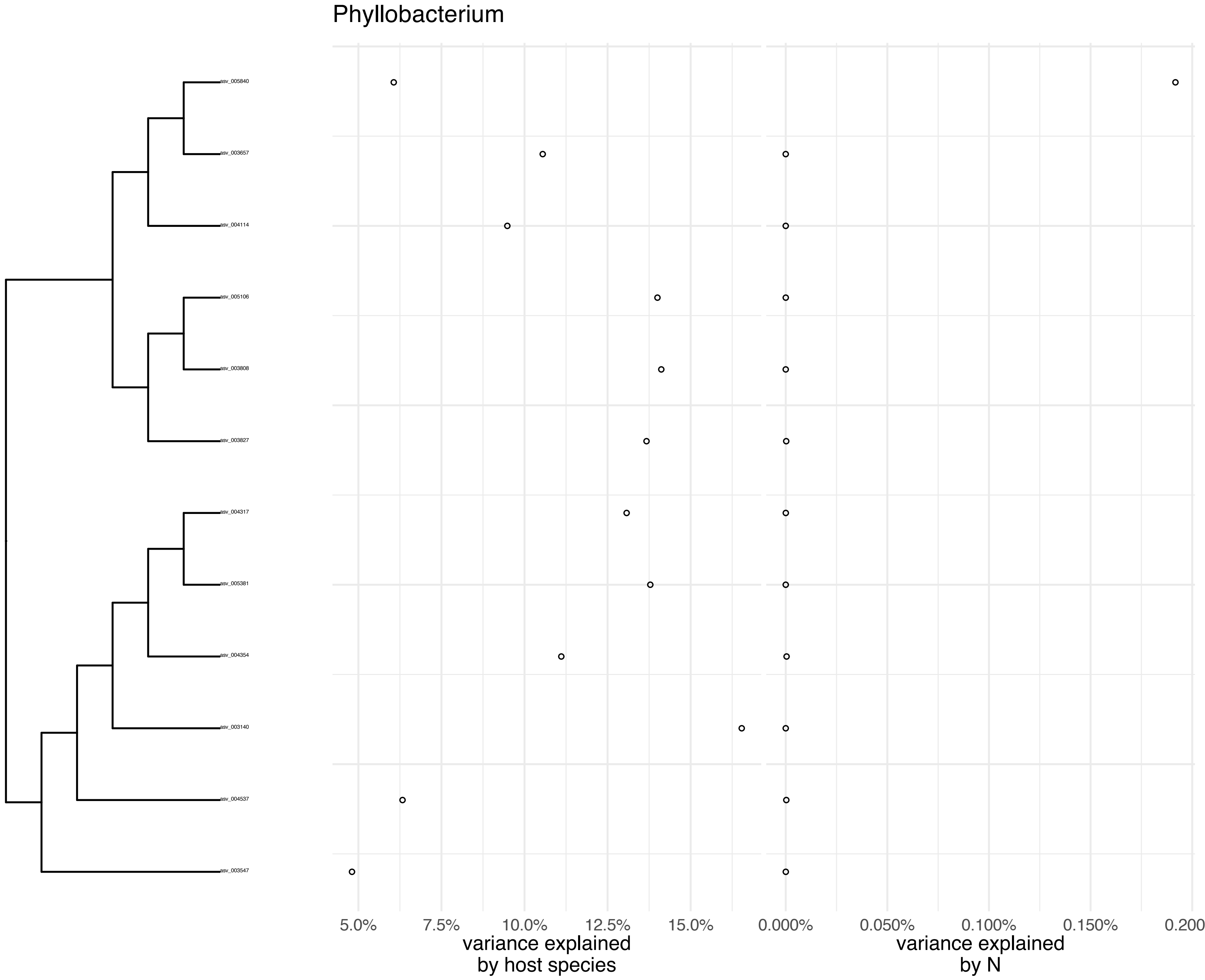
Phyllobacterium
asv_005840
asv_003657
asv_004114
asv_005106
asv_003808
asv_003827
asv_004317
asv_005381
asv_004354
asv_003140
asv_004537
asv_003547
5.0%
7.5%
10.0%
12.5%
15.0%
0.000%
0.050%
0.100%
0.150%
0.200
variance explained
by host species
variance explained
by N

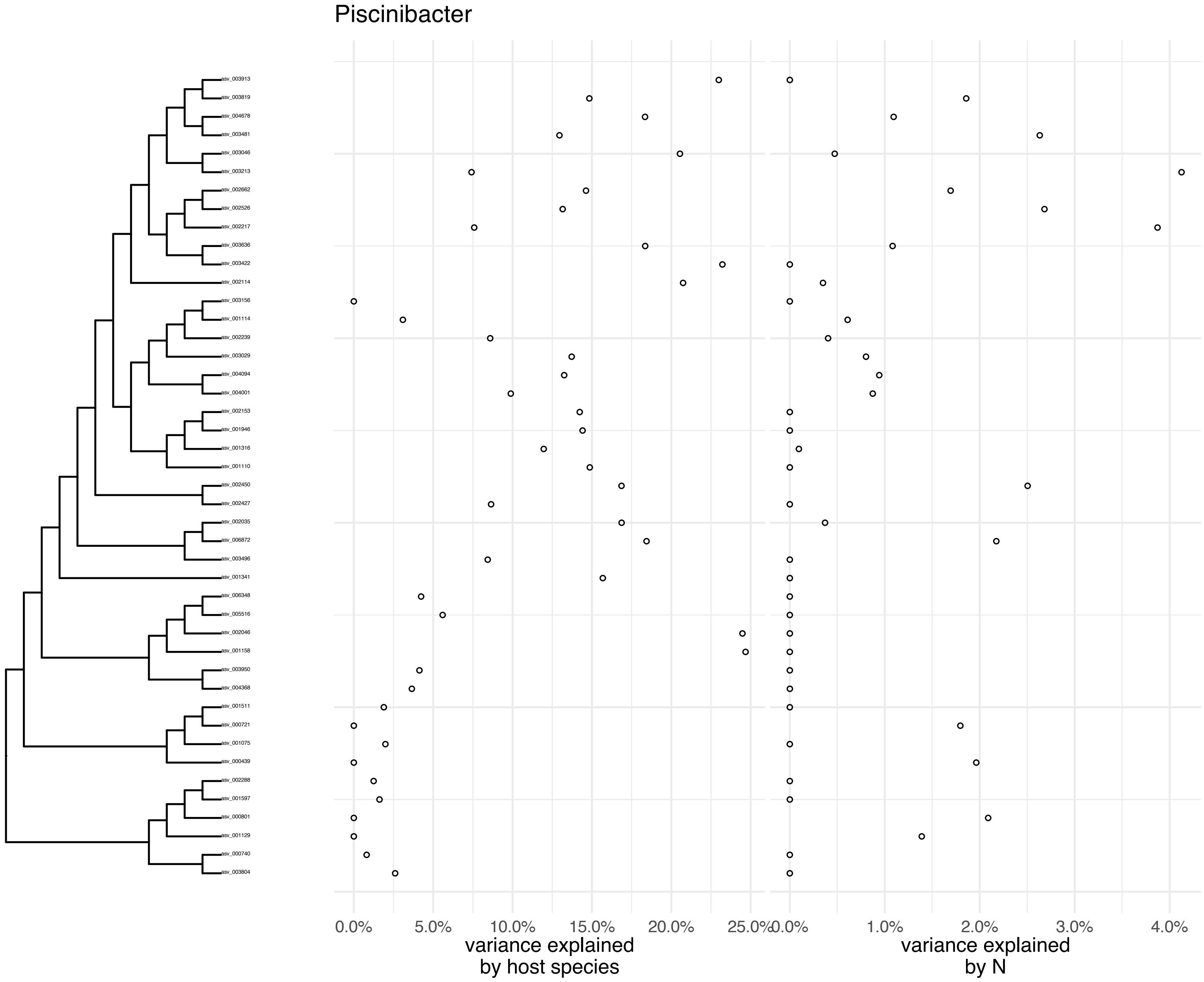

Piscinibacter
asv_003913
asv_003819
asv_004678
asv_003481
asv_003046
asv_003213
asv_002662
asv_002526
asv_002217
asv_003636
asv_003422
asv_002114
asv_003156
asv_001114
asv_002239
asv_003029
asv_004094
asv_004001
asv_002153
asv_001946
asv_001316
asv_001110
asv_002450
asv_002427
asv_002035
asv_006872
asv_003496
asv_001341
asv_006348
asv_005516
asv_002046
asv_001158
asv_003950
asv_004368
asv_001511
asv_000721
asv_001075
asv_000439
asv_002288
asv_001597
asv_000801
asv_001129
asv_000740
asv_003804
0.0%
5.0%
10.0%
15.0%
20.0%
25.0%
variance explained by host species
0.0%
1.0%
2.0%
3.0%
4.0%
variance explained by N

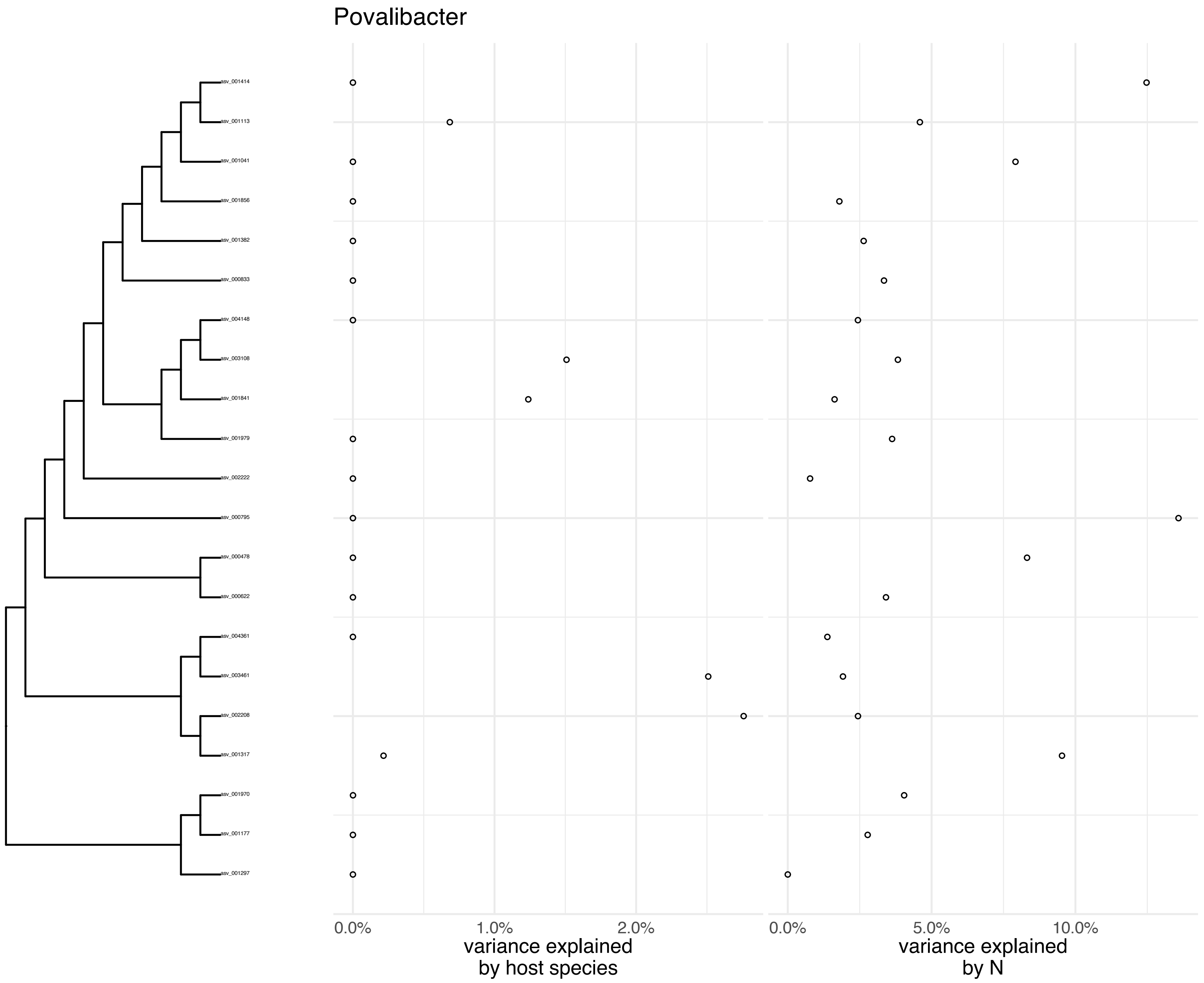
Povalibacter
asv_001414
asv_001113
asv_001041
asv_001856
asv_001382
asv_000833
asv_004148
asv_003108
asv_001841
asv_001979
asv_002222
asv_000795
asv_000478
asv_000622
asv_004361
asv_003461
asv_002208
asv_001317
asv_001970
asv_001177
asv_001297
0.0%
1.0%
2.0%
variance explained by host species
0.0%
5.0%
10.0%
variance explained by N

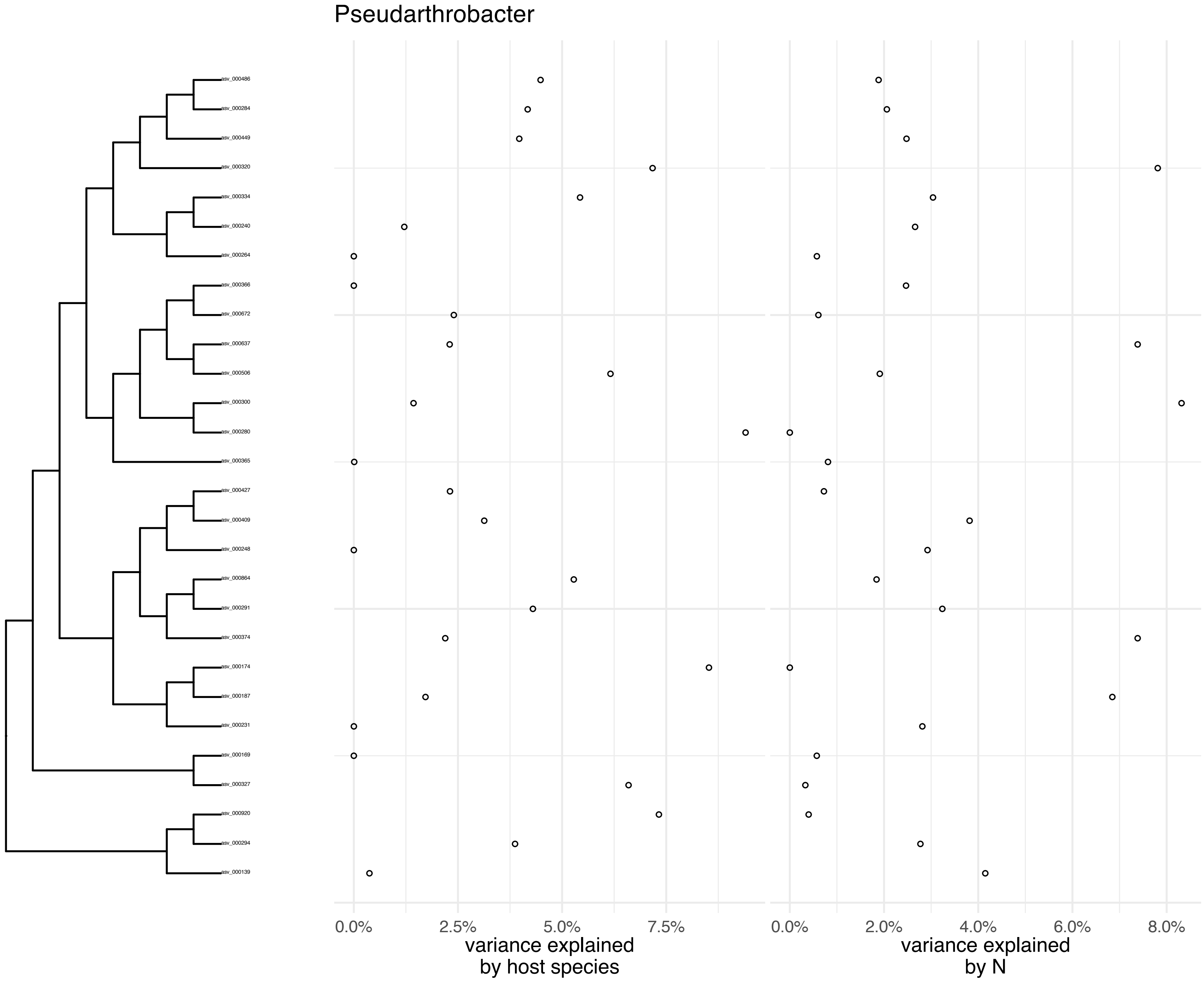

Pseudarthrobacter
0.0%
2.5%
5.0%
7.5%
variance explained
by host species
0.0%
2.0%
4.0%
6.0%
8.0%
variance explained
by N

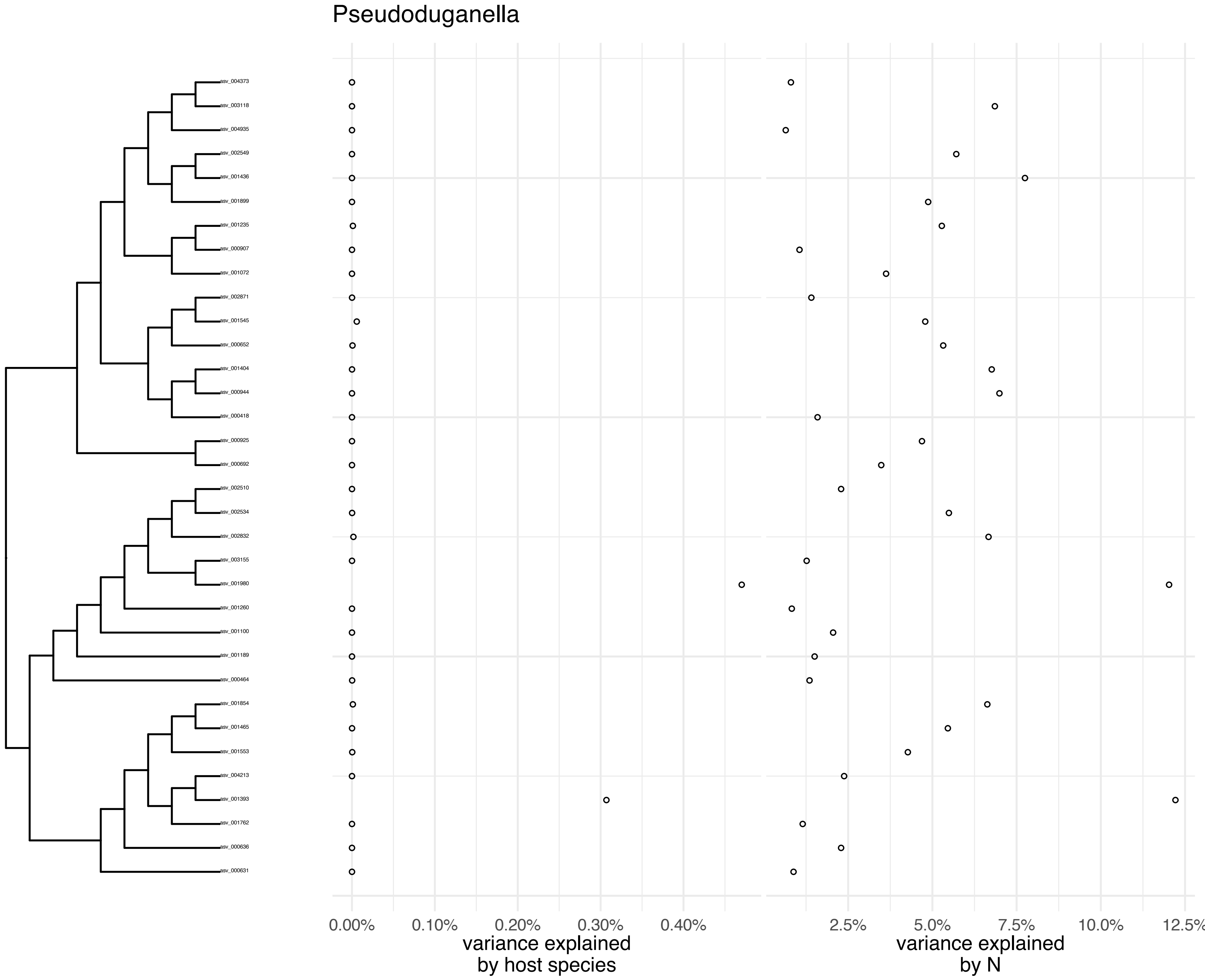

Pseudoduganella
asv_004373
asv_003118
asv_004935
asv_002549
asv_001436
asv_001899
asv_001235
asv_000907
asv_001072
asv_002871
asv_001545
asv_000652
asv_001404
asv_000944
asv_000418
asv_000925
asv_000692
asv_002510
asv_002534
asv_002832
asv_003155
asv_001980
asv_001260
asv_001100
asv_001189
asv_000464
asv_001854
asv_001465
asv_001553
asv_004213
asv_001393
asv_001762
asv_000636
asv_000631
0.00%
0.10%
0.20%
0.30%
0.40%
variance explained
by host species
2.5%
5.0%
7.5%
10.0%
12.5%
variance explained
by N

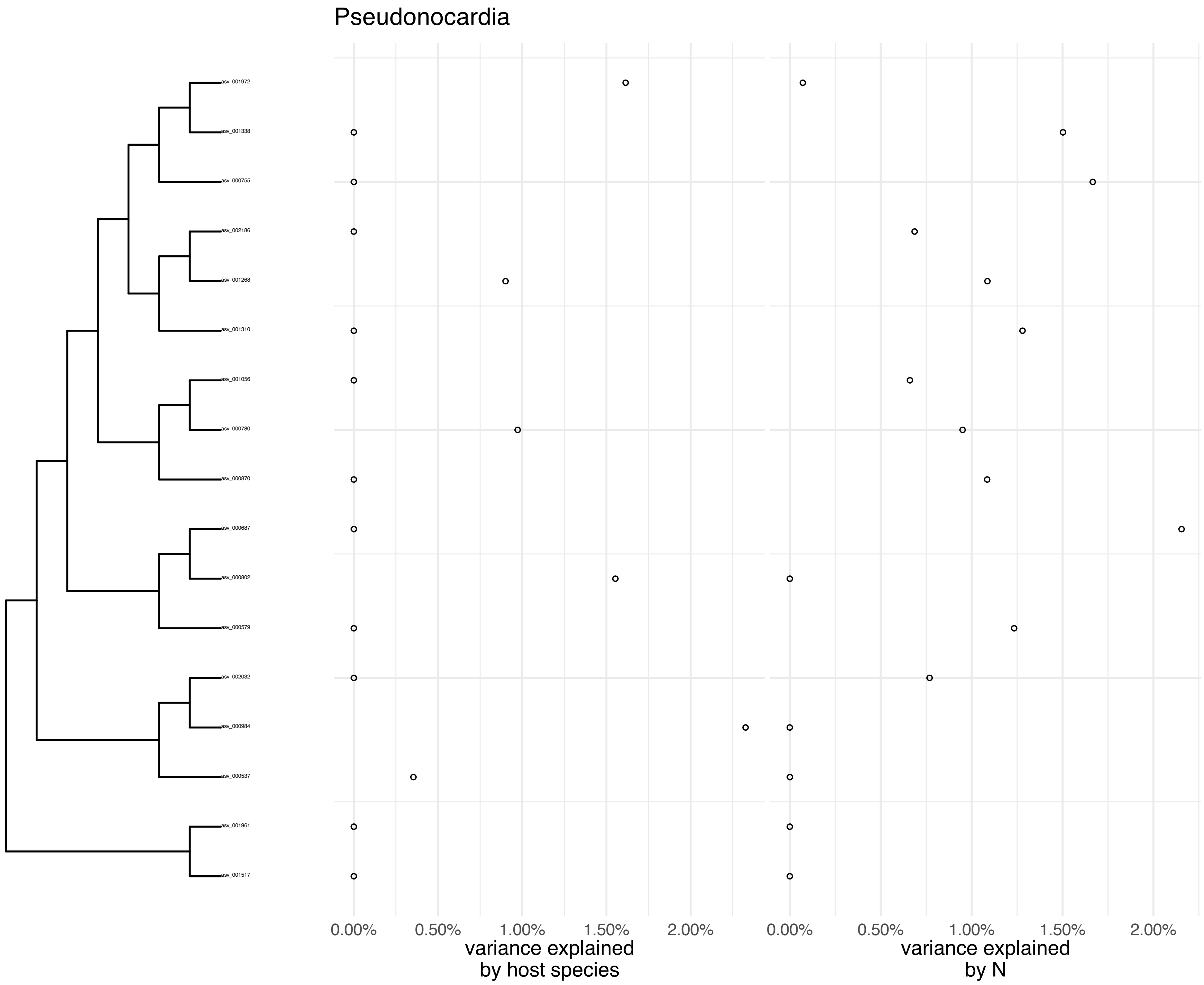

Pseudonocardia
asv_001972
asv_001338
asv_000755
asv_002186
asv_001268
asv_001310
asv_001056
asv_000780
asv_000870
asv_000687
asv_000802
asv_000579
asv_002032
asv_000984
asv_000537
asv_001961
asv_001517
0.00%
0.50%
1.00%
1.50%
2.00%
variance explained by host species
0.00%
0.50%
1.00%
1.50%
2.00%
variance explained by N

### Psychroglaciecola

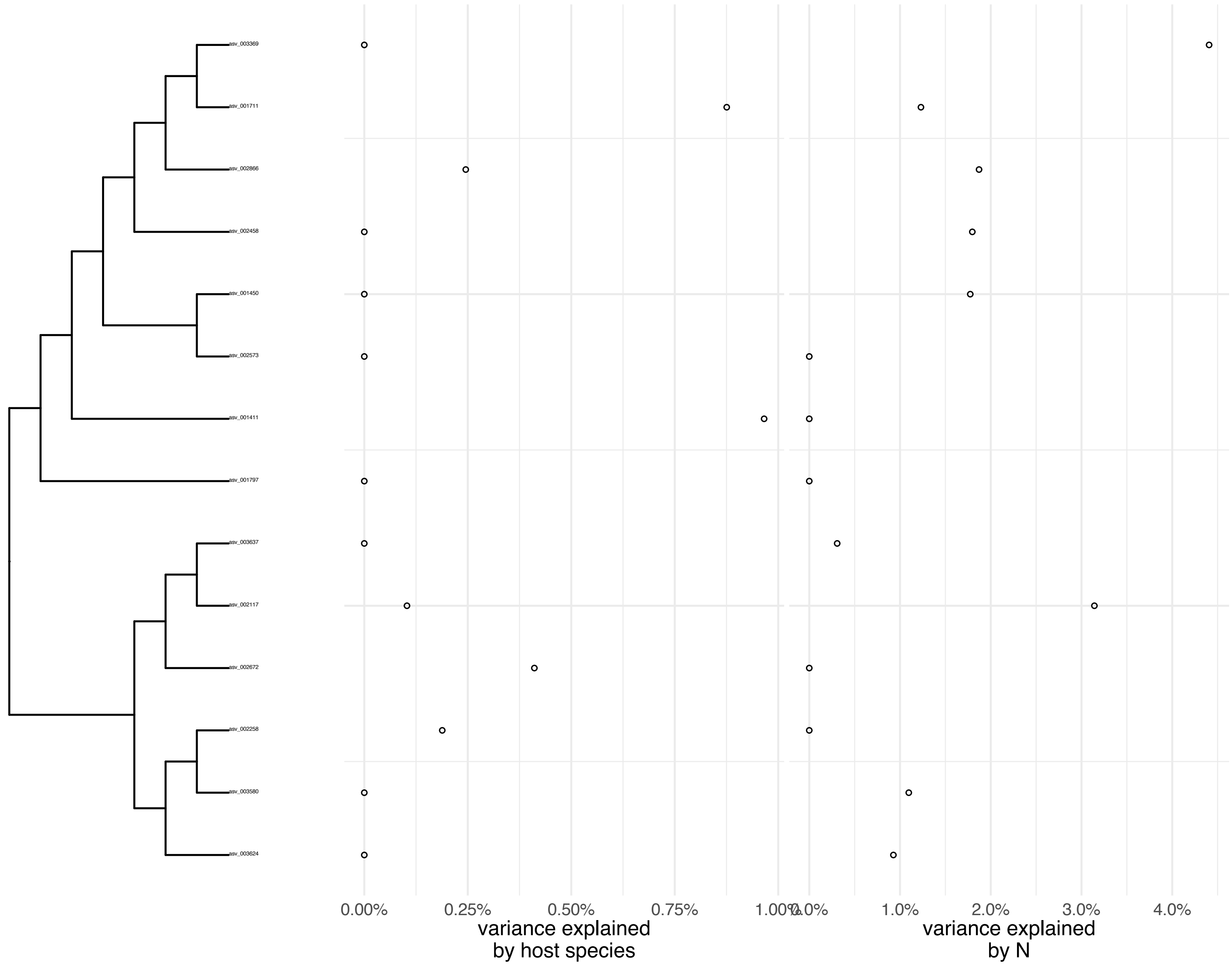

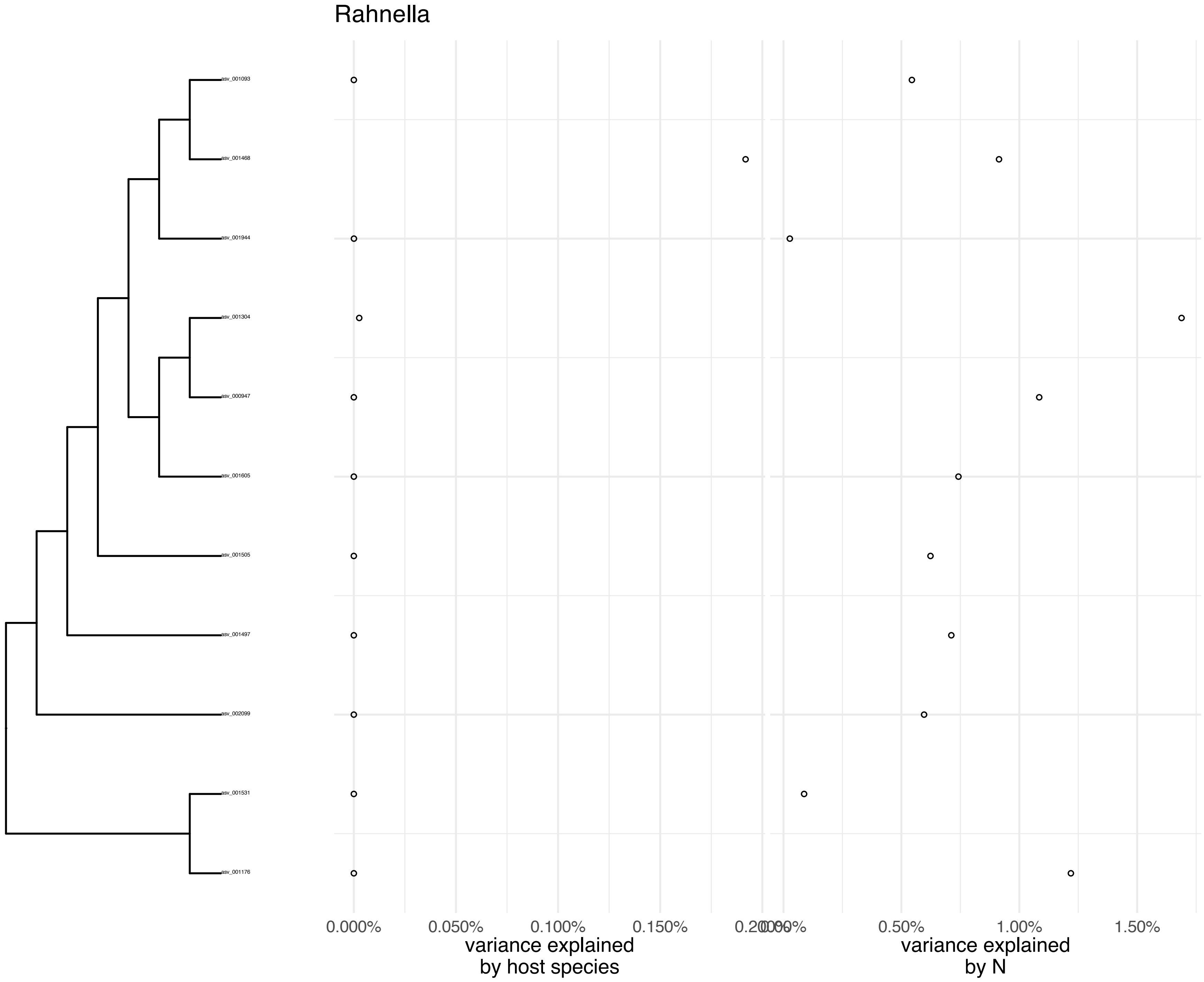

Rahnella
asv_001093
asv_001468
asv_001944
asv_001304
asv_000947
asv_001605
asv_001505
asv_001497
asv_002099
asv_001531
asv_001176
0.000%
0.050%
0.100%
0.150%
0.200%
0.00%
0.50%
1.00%
1.50%
variance explained by host species
variance explained by N

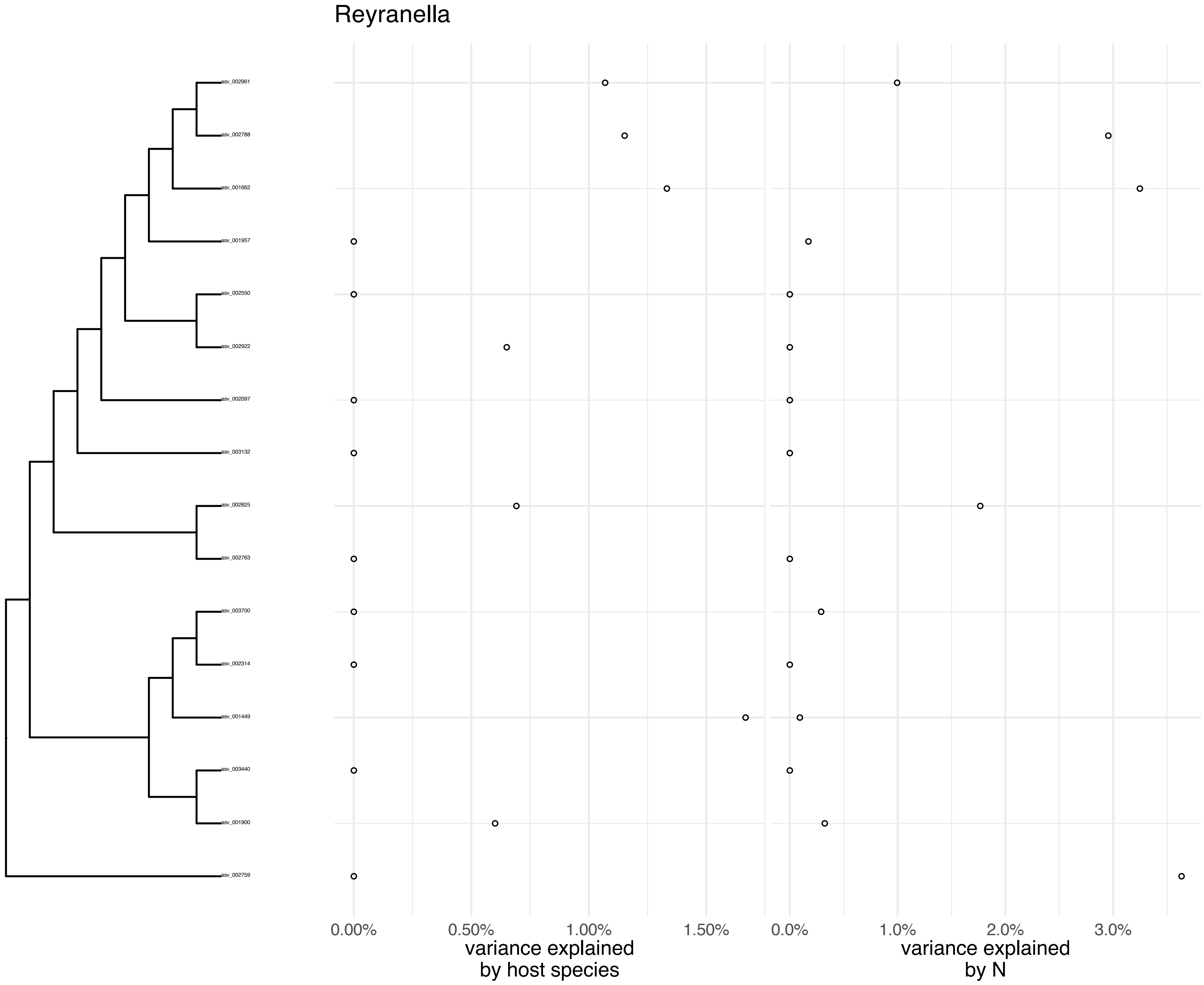

Reyranella
asv_002961
asv_002788
asv_001662
asv_001957
asv_002550
asv_002922
asv_002097
asv_003132
asv_002825
asv_002763
asv_003700
asv_002314
asv_001449
asv_003440
asv_001900
asv_002759
0.00%
0.50%
1.00%
1.50%
variance explained by host species
0.0%
1.0%
2.0%
3.0%
variance explained by N

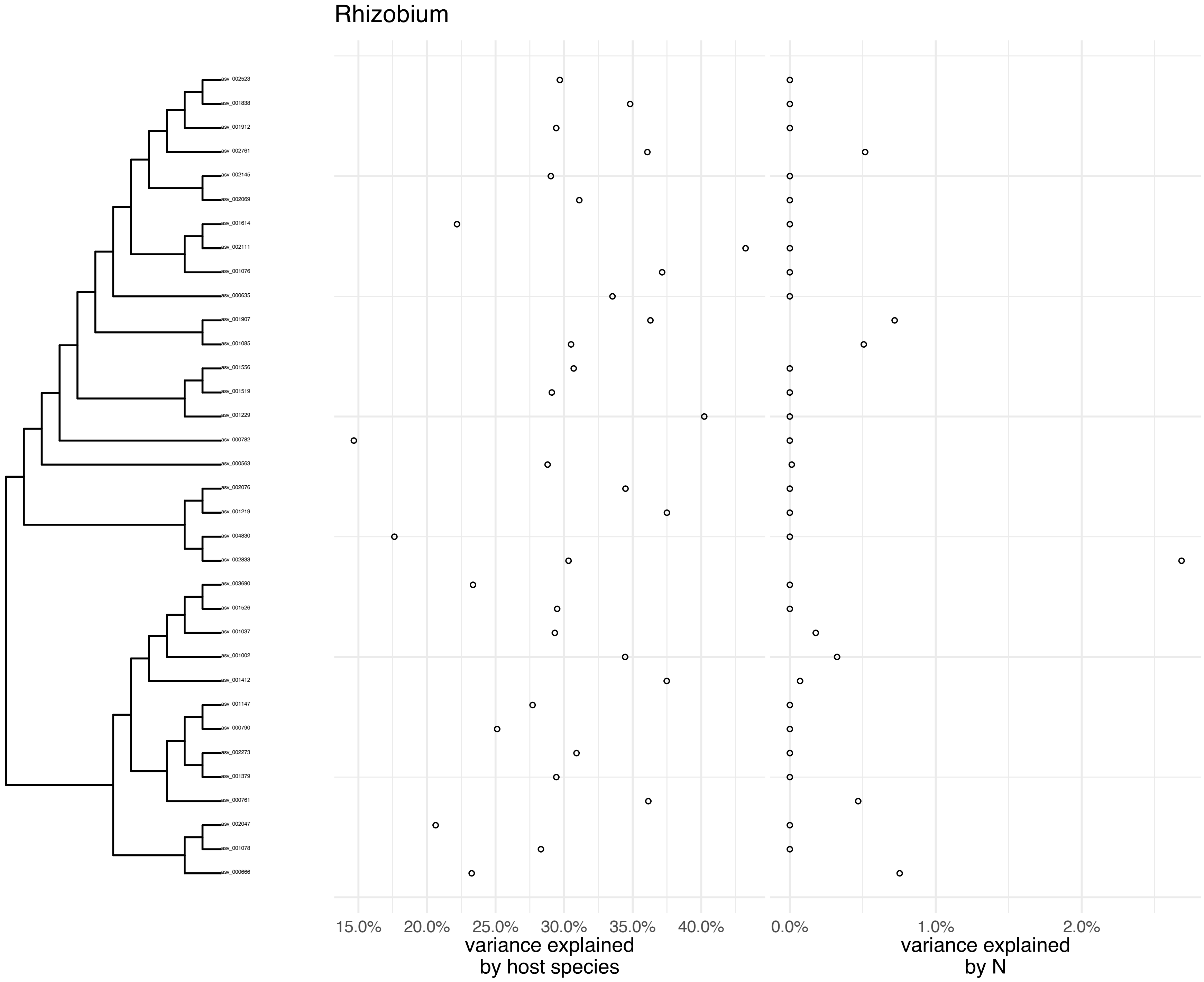
Rhizobium
asv_002523
asv_001838
asv_001912
asv_002761
asv_002145
asv_002069
asv_001614
asv_002111
asv_001076
asv_000635
asv_001907
asv_001085
asv_001556
asv_001519
asv_001229
asv_000782
asv_000563
asv_002076
asv_001219
asv_004830
asv_002833
asv_003690
asv_001526
asv_001037
asv_001002
asv_001412
asv_001147
asv_000790
asv_002273
asv_001379
asv_000761
asv_002047
asv_001078
asv_000666
15.0%
20.0%
25.0%
30.0%
35.0%
40.0%
variance explained
by host species
0.0%
1.0%
2.0%
variance explained
by N

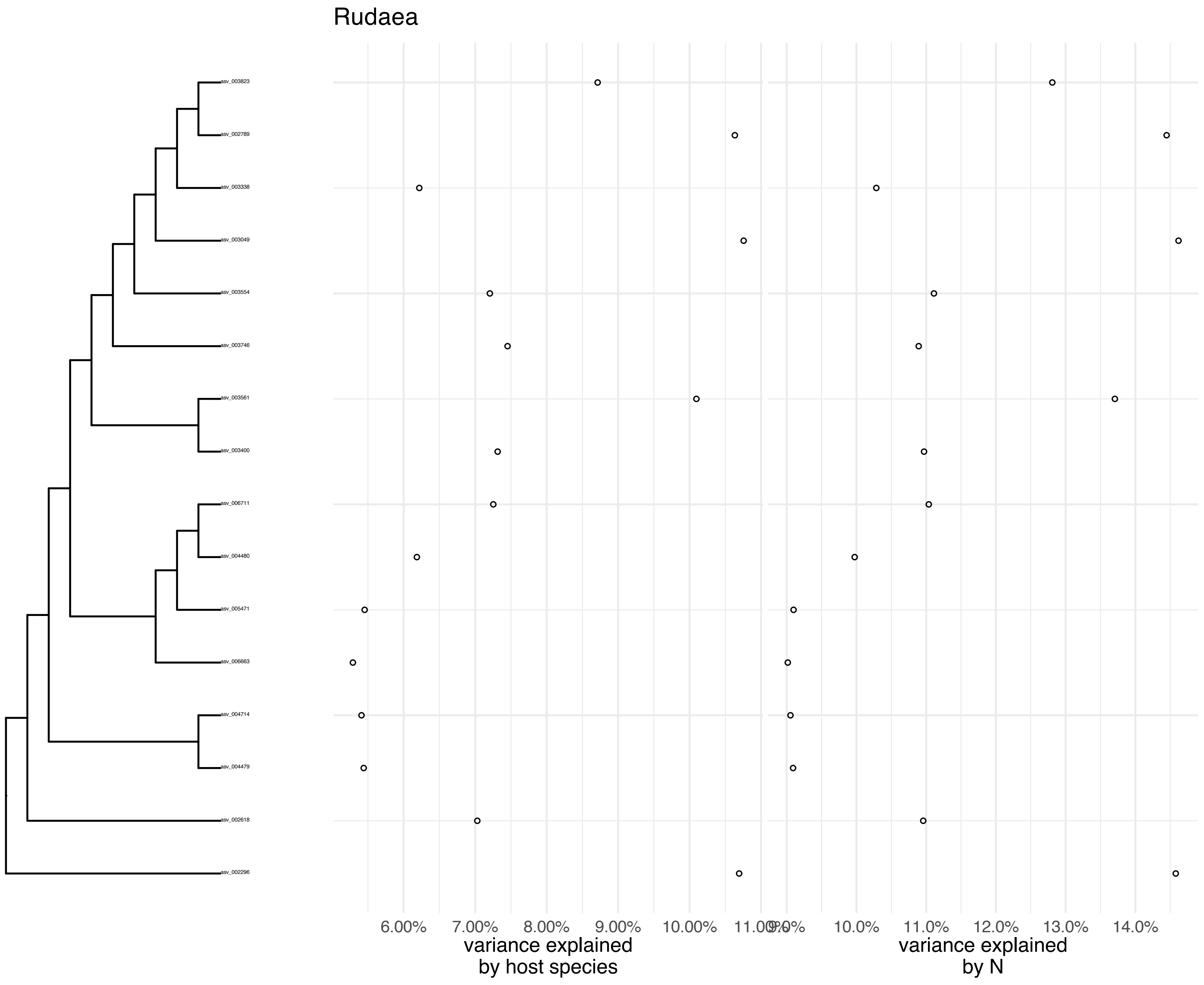
Rudaea
asv_003823
asv_002789
asv_003338
asv_003049
asv_003554
asv_003746
asv_003561
asv_003400
asv_006711
asv_004480
asv_005471
asv_006663
asv_004714
asv_004479
asv_002618
asv_002296
6.00%
7.00%
8.00%
9.00%
10.00%
11.00%
9.0%
10.0%
11.0%
12.0%
13.0%
14.0%
variance explained
by host species
variance explained
by N

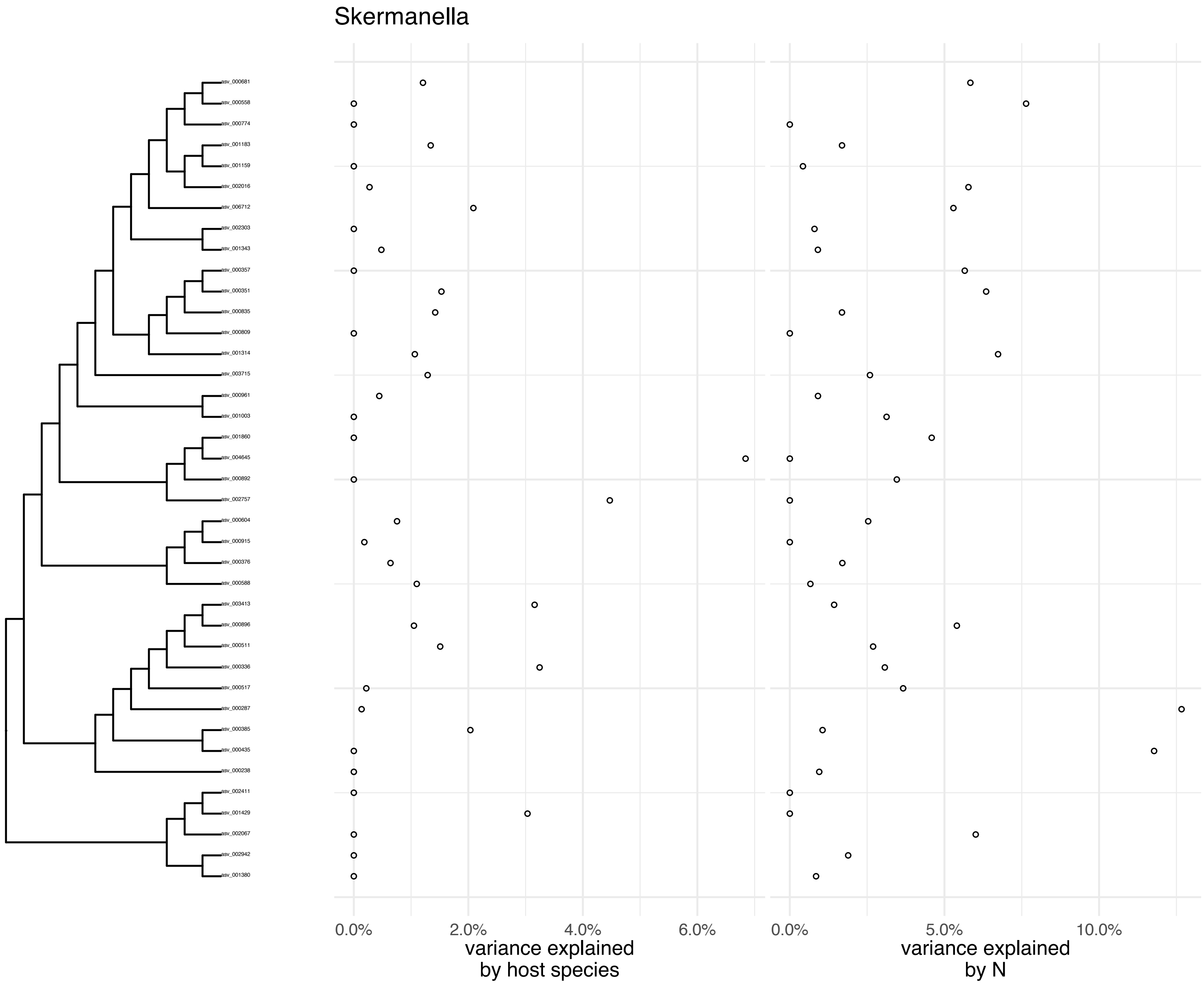
Skermanella
asv_000681
asv_000558
asv_000774
asv_001183
asv_001159
asv_002016
asv_006712
asv_002303
asv_001343
asv_000357
asv_000351
asv_000835
asv_000809
asv_001314
asv_003715
asv_000961
asv_001003
asv_001860
asv_004645
asv_000892
asv_002757
asv_000604
asv_000915
asv_000376
asv_000588
asv_003413
asv_000896
asv_000511
asv_000336
asv_000517
asv_000287
asv_000385
asv_000435
asv_000238
asv_002411
asv_001429
asv_002067
asv_002942
asv_001380
0.0%
2.0%
4.0%
6.0%
variance explained
by host species
0.0%
5.0%
10.0%
variance explained
by N

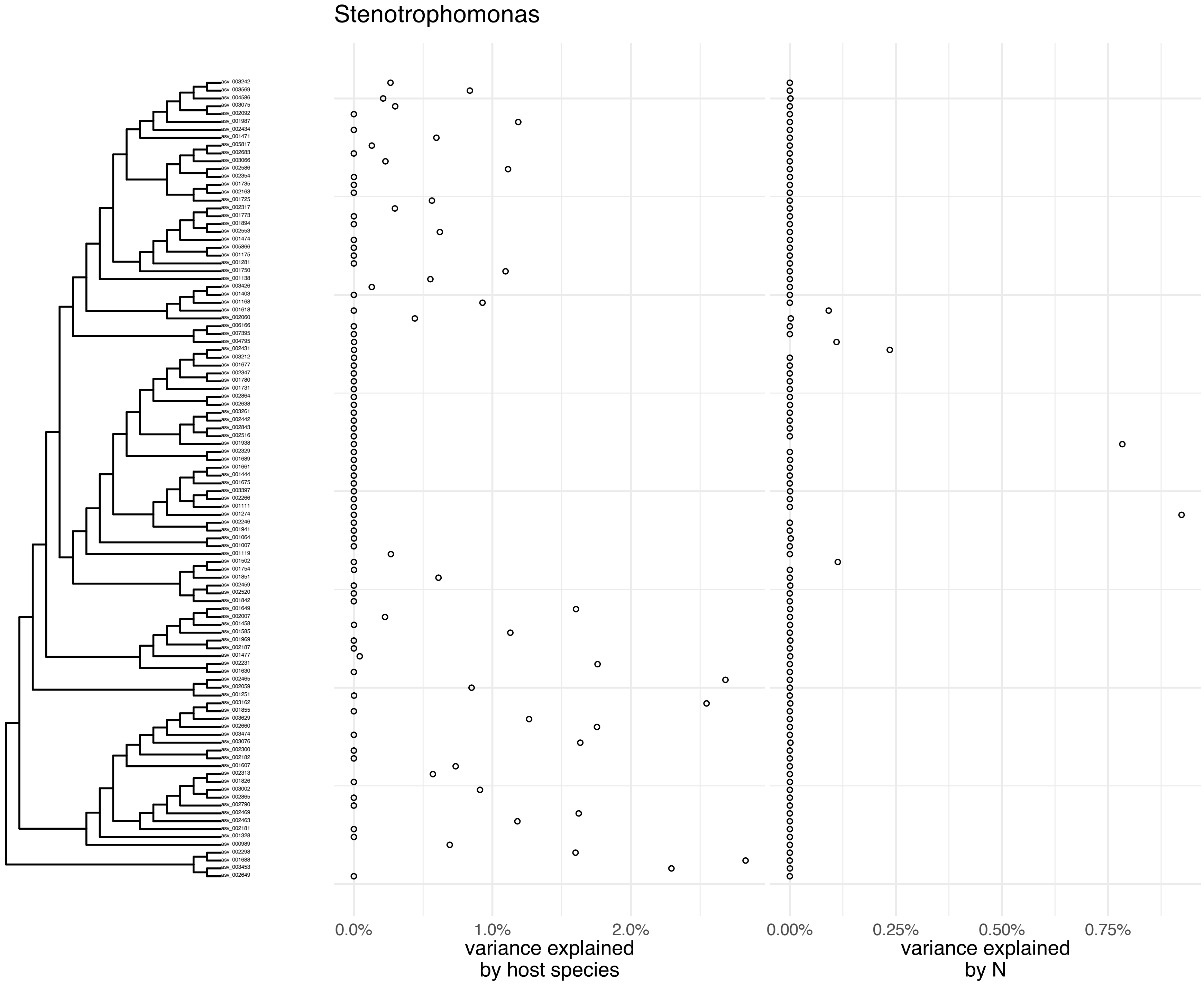
Stenotrophomonas
0.0%
1.0%
2.0%
variance explained
by host species
0.00%
0.25%
0.50%
0.75%
variance explained
by N

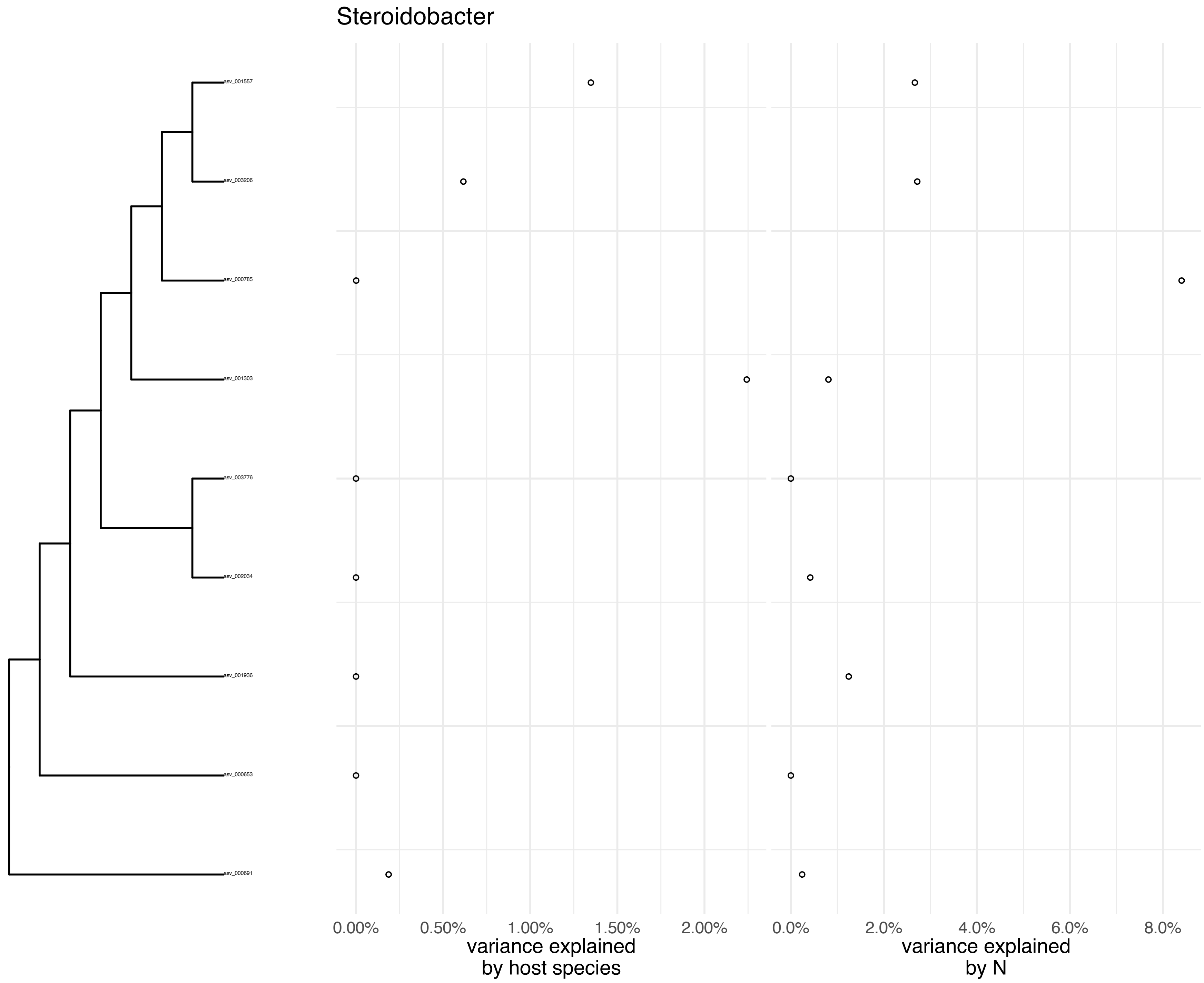
Steroidobacter
asv_001557
asv_003206
asv_000785
asv_001303
asv_003776
asv_002034
asv_001936
asv_000653
asv_000691
0.00%
0.50%
1.00%
1.50%
2.00%
variance explained
by host species
0.0%
2.0%
4.0%
6.0%
8.0%
variance explained
by N

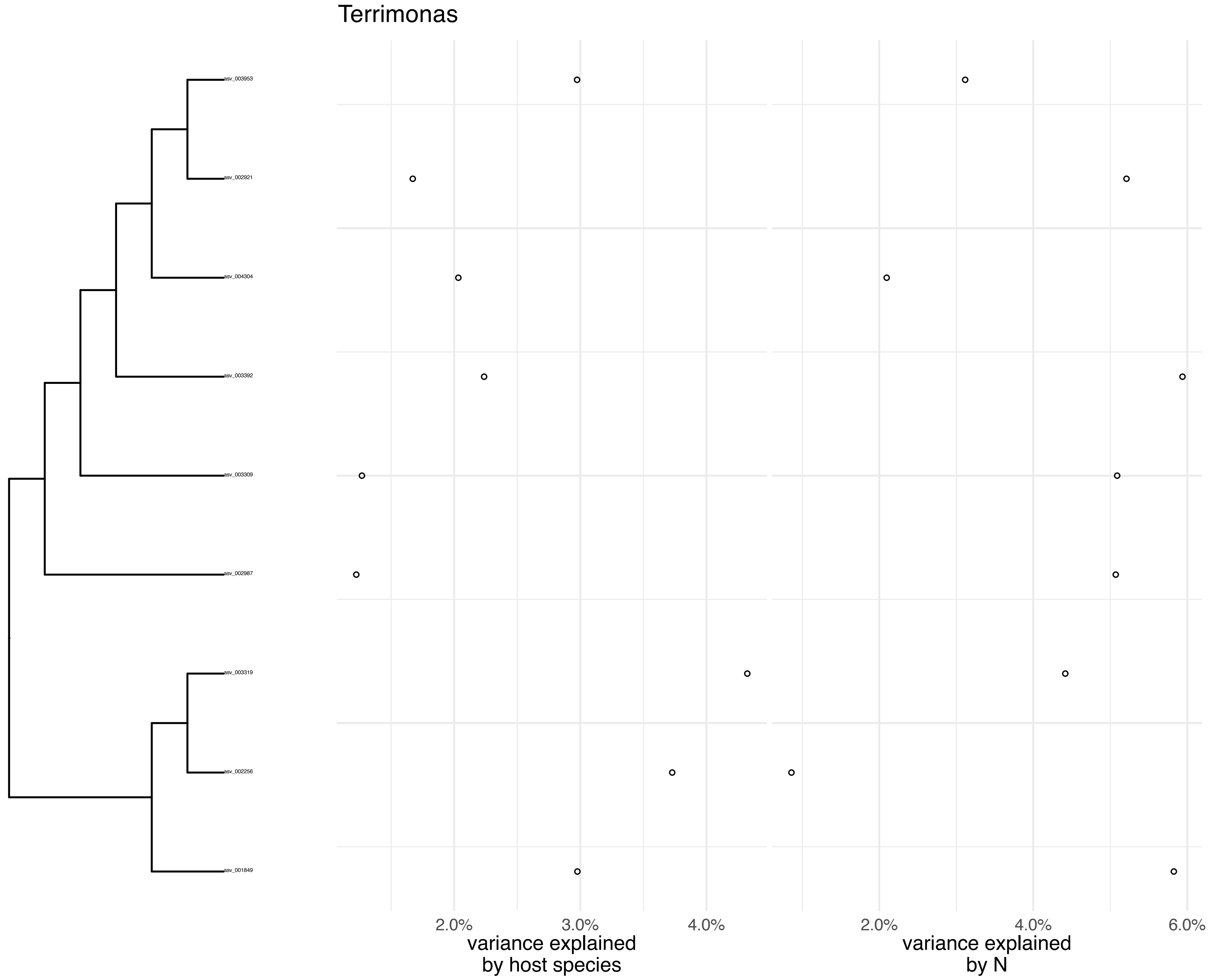
Terrimonas
asv_003953
asv_002921
asv_004304
asv_003392
asv_003309
asv_002987
asv_003319
asv_002256
asv_001849
2.0%
3.0%
4.0%
variance explained
by host species
2.0%
4.0%
6.0%
variance explained
by N

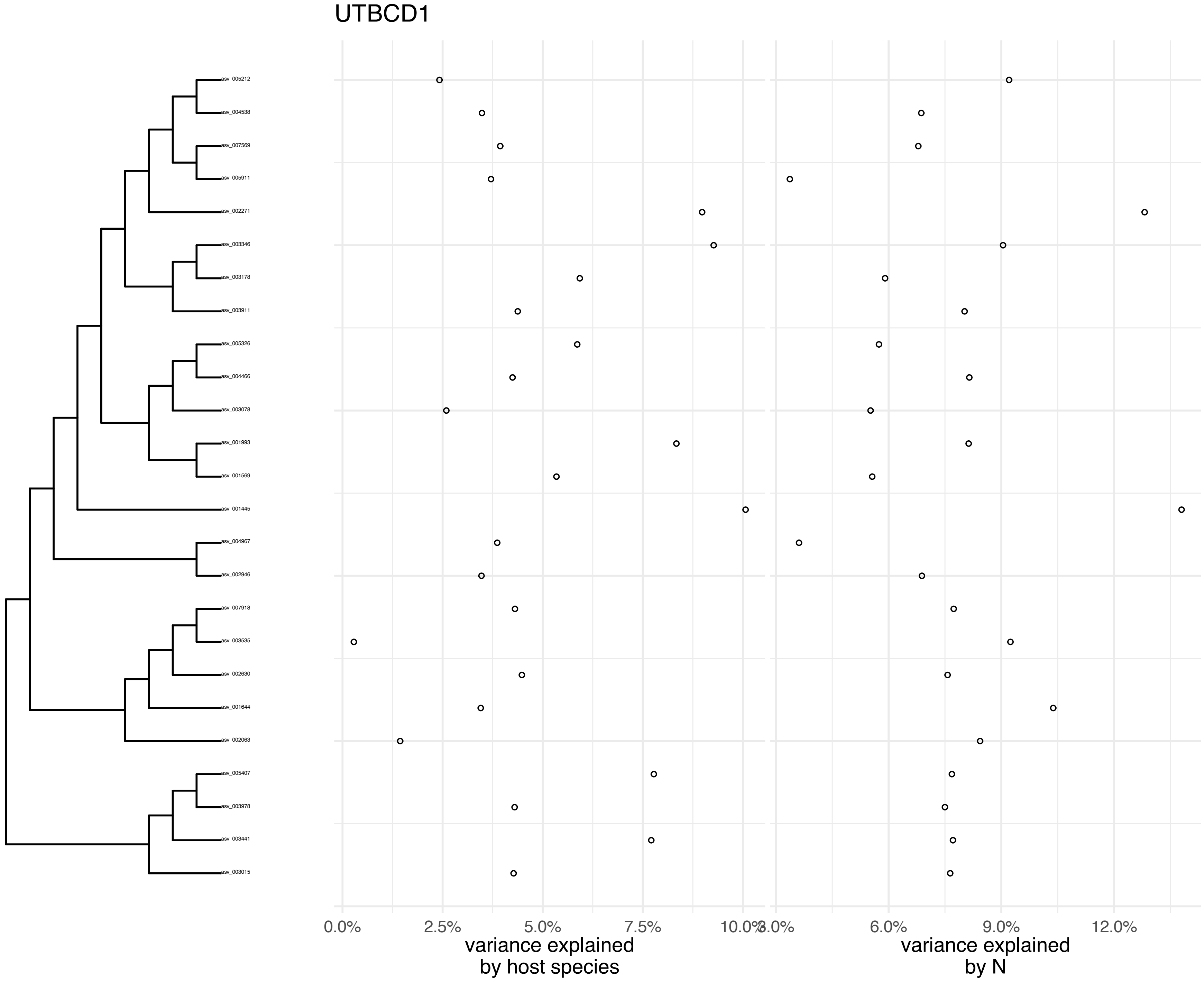
UTBCD1
asv_005212
asv_004538
asv_007569
asv_005911
asv_002271
asv_003346
asv_003178
asv_003911
asv_005326
asv_004466
asv_003078
asv_001993
asv_001569
asv_001445
asv_004967
asv_002946
asv_007918
asv_003535
asv_002630
asv_001644
asv_002063
asv_005407
asv_003978
asv_003441
asv_003015
0.0%
2.5%
5.0%
7.5%
10.0%
3.0%
6.0%
9.0%
12.0%
variance explained
by host species
variance explained
by N

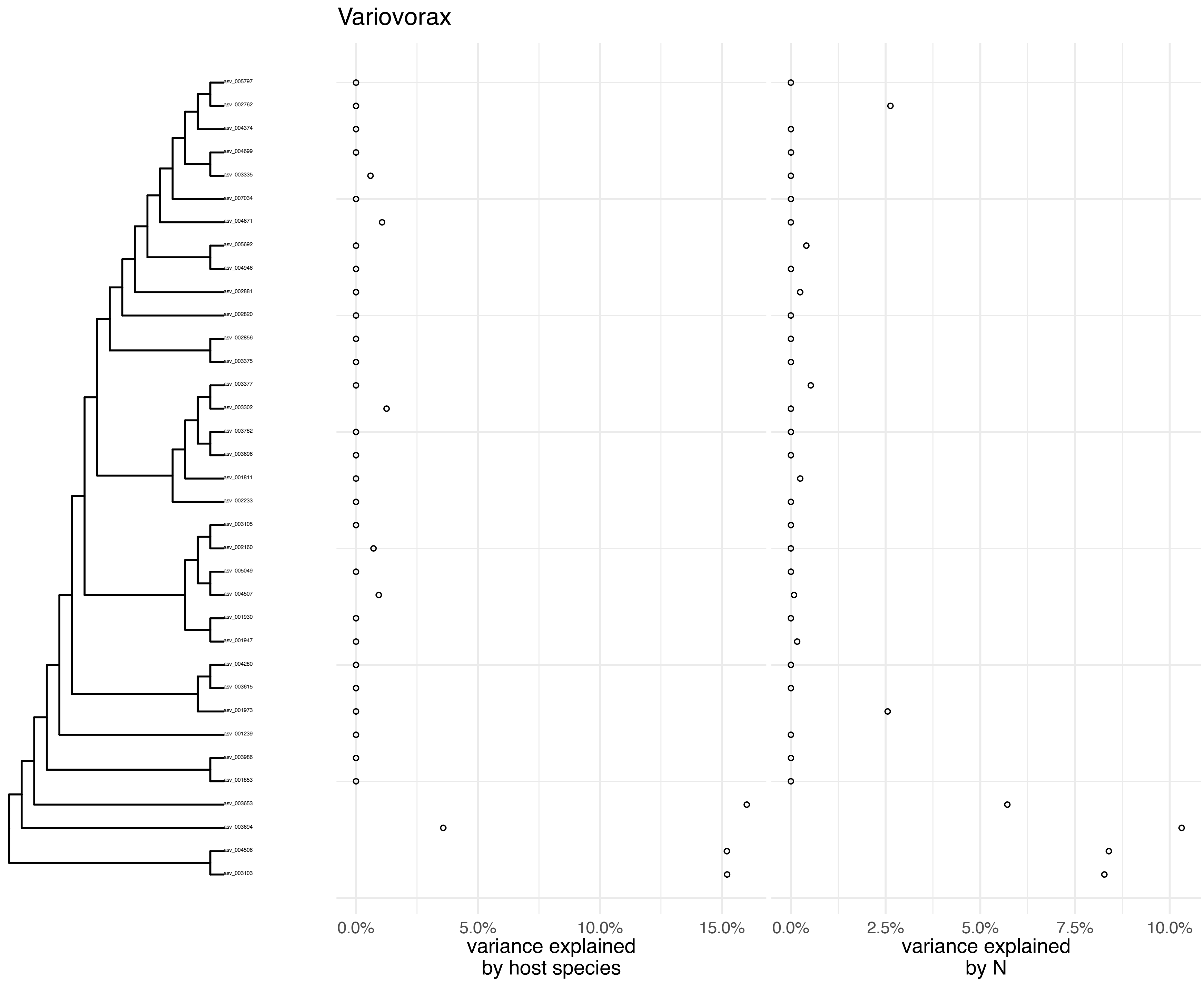

Variovorax
asv_005797
asv_002762
asv_004374
asv_004699
asv_003335
asv_007034
asv_004671
asv_005692
asv_004946
asv_002881
asv_002820
asv_002856
asv_003375
asv_003377
asv_003302
asv_003782
asv_003696
asv_001811
asv_002233
asv_003105
asv_002160
asv_005049
asv_004507
asv_001930
asv_001947
asv_004280
asv_003615
asv_001973
asv_001239
asv_003986
asv_001853
asv_003653
asv_003694
asv_004506
asv_003103
0.0%
5.0%
10.0%
15.0%
variance explained
by host species
0.0%
2.5%
5.0%
7.5%
10.0%
variance explained
by N

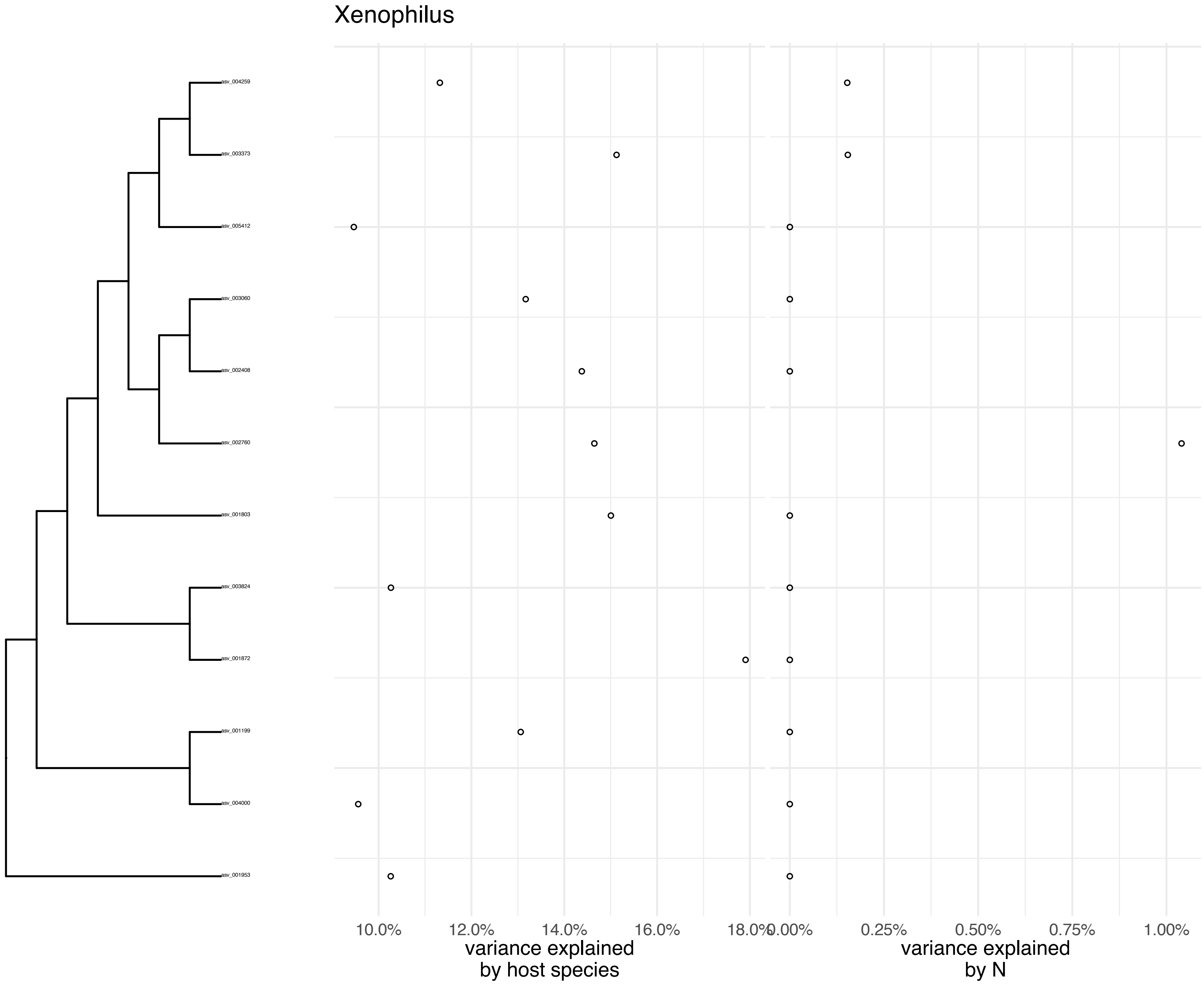
Xenophilus
asv_004259
asv_003373
asv_005412
asv_003060
asv_002408
asv_002760
asv_001803
asv_003824
asv_001872
asv_001199
asv_004000
asv_001953
10.0%
12.0%
14.0%
16.0%
18.0%
0.00%
0.25%
0.50%
0.75%
1.00%
variance explained
by host species
variance explained
by N

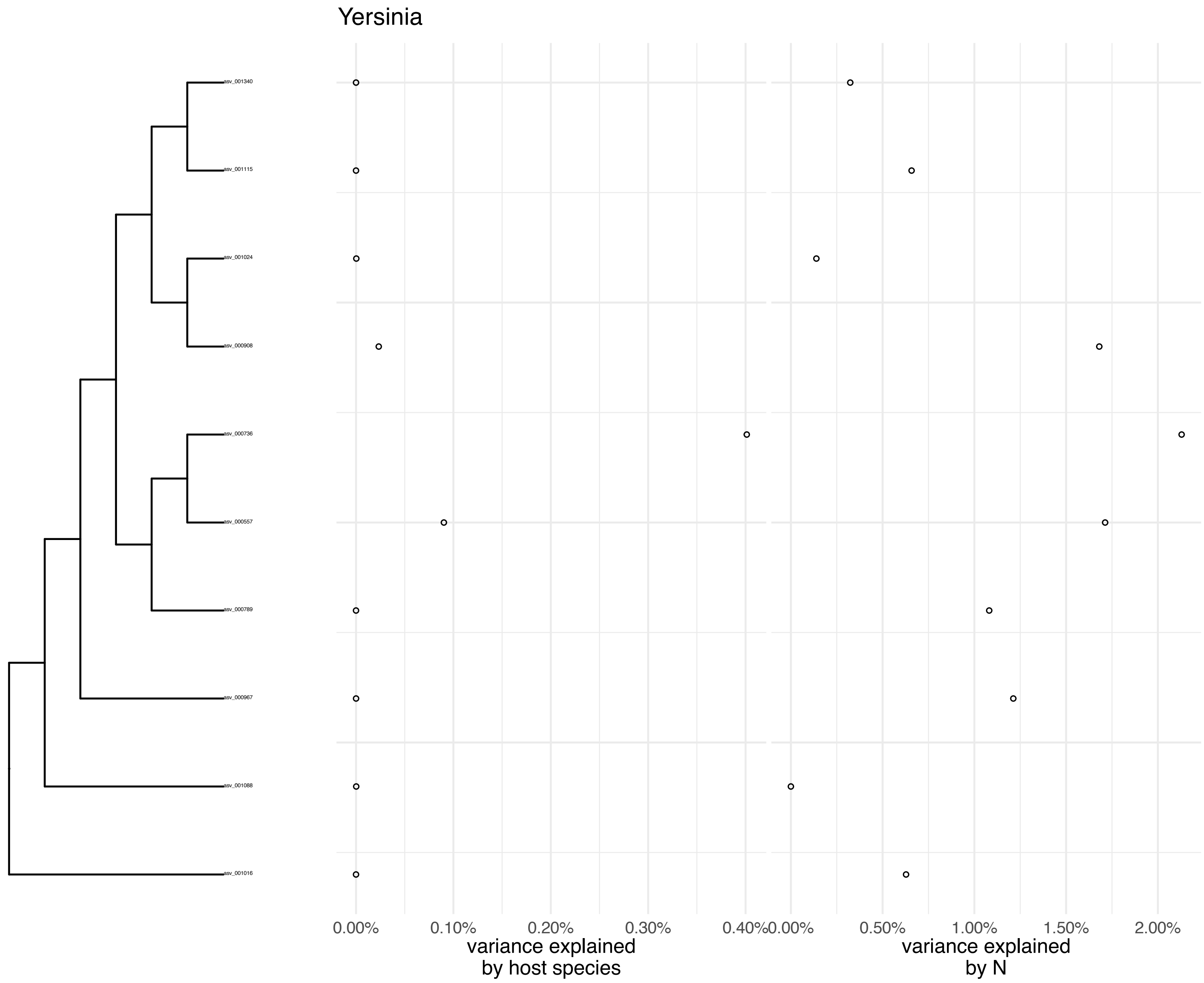
Yersinia
asv_001340
asv_001115
asv_001024
asv_000908
asv_000736
asv_000557
asv_000789
asv_000967
asv_001088
asv_001016
0.00%
0.10%
0.20%
0.30%
0.40%
variance explained
by host species
0.00%
0.50%
1.00%
1.50%
2.00%
variance explained
by N
